# Assessing target genes for homing suppression gene drive

**DOI:** 10.1101/2024.12.06.627146

**Authors:** Xuejiao Xu, Jialing Fang, Jingheng Chen, Jie Yang, Xiaozhen Yang, Shibo Hou, Weitang Sun, Jackson Champer

**Affiliations:** Center for Bioinformatics, Center for Life Sciences, School of Life Sciences, Peking University, Beijing, China 100871

## Abstract

Gene drives are engineered alleles that bias their own inheritance in offspring, enabling the spread of specific traits throughout a population. Targeting female fertility genes in a gene drive can be an efficient strategy for population suppression. In this study, we investigated nine female fertility genes in *Drosophila melanogaster* using CRISPR-based homing gene drives. Employing a multiplexed gRNA approach to prevent the formation of functional resistance alleles, we aimed to maintain high drive conversion efficiency with low fitness costs in female drive carriers. Drive efficiency was assessed in individual crosses and had varied performance across different target genes. Notably, drives targeting the *octopamine β2 receptor* (*oct*) and *stall* (*stl*) genes exhibited the highest drive conversion rates and were further tested in cages. A drive targeting *stl* successfully suppressed a cage population with a high release frequency, though suppression failed in another replicate cage with a lower initial release frequency. Fitness costs in female drive carriers were observed in test cages, impacting the overall efficiency of population suppression. Further tests on the fertility of these lines using individual crosses indicated that some fitness costs were due to maternal deposition of Cas9 combined with new gRNA expression, which would only occur in progeny of drive males when testing split drives with separate Cas9 (when mimicking cages with complete drives) but not for complete drive systems. This could enable success in complete drives with higher maternal Cas9 deposition, even if cage experiments in split drives fail. Our findings underscore the potential and challenges of assessing gene drives for population control, providing valuable insights for optimizing and testing suppression gene drive designs.

## Introduction

Gene drive systems are a class of selfish genetic elements existing in nature, naturally biasing their own inheritance in the population. Initially, homing endonuclease genes were proposed for engineering synthetic gene drives, but target selection was constrained by the specific sequences required for endonuclease activity. The advent of CRISPR-based nucleases, however, has vastly expanded target options through engineered guide RNA (gRNA) sequences, catalyzing the development of diverse gene drive tools^1–5^. These approaches hold promise for ecologically friendly pest control strategies, with potential applications in species conservation, agriculture protection, and disease prevention.

CRISPR-based homing gene drives operate by utilizing Cas9 nuclease to cleave a target site determined by the gRNA, copying the drive element into the target allele through homology-directed repair. This process, known as drive conversion or homing, converts heterozygotes to homozygotes in the germline, ensuring the gene drive is inherited by the majority of offspring. Consequently, the gene drive frequency increases across generations, potentially leading to fixation of the drive allele. Depending on application goals and ecological considerations, gene drives can be designed to either modify or suppress target populations. Modification drives aim to alter populations without significantly impacting their population size, maintaining ecological niche and balance while introducing specific traits (e.g. anti-disease^6,7^ or insecticide susceptibility^8^) in these populations. In contrast, suppression drives are designed to quickly reduce or eliminate target populations, directly preventing agricultural loss^9–11^, disease transmission^12–14^, or other negative effects caused by invasive species^15,16^.

For suppression drives, target genes should be haplosufficient but essential. They can affect both sexes^17^ but will be more powerful when targeting female viability or reproduction. Disrupting these genes results in female mortality or sterility, reducing offspring numbers and ultimately leading to population collapse. This type of drive tends to reach an equilibrium when the drive conversion process is not perfectly efficient, thus placing a genetic load (reflecting suppressive power) on the target population, which can still eliminate the population if it is sufficiently high. A previous study^18^ has demonstrated homing suppression drives respectively targeting three female fertility genes in malaria vector *Anopheles gambiae*. Among these, the drive targeting *nudel* met the minimum requirement to spread in cage populations initially but ultimately failed due to fitness costs in female heterozygotes from somatic Cas9 expression and functional resistant allele formation (see below). A drive in *Drosophila melanogaster* targeting *tra* suffered from even higher fitness costs in females due to somatic Cas9 expression^19^. One of the most successful population suppression gene drives to date targeted the female specific splicing site of the sex determination gene *doublesex* in *A. gambiae*^12^. Despite the promising performance, translating this drive to other species has not always met expectations. For instance, dominant female sterility was observed in both *Drosophila suzukii*^9^ and *D. melanogaster*^20^, and male drive homozygotes in *Anopheles stephensi* exhibited sterility^21^, reducing suppression efficacy. *dsx* is also an important developmental gene, required in many somatic tissues, which can be difficult for Cas9 promoters with somatic expression. These findings underscore the need for further exploration of female fertility or viability targets to broaden the options for effective population suppression.

Resistance alleles, which arise from end-joining repair rather than homology-directed repair following Cas9/gRNA cleavage, present a challenge to gene drive efficacy^22^. They have mismatches in the target sequences and therefore cannot be recognized and cut by Cas9/gRNA. These alleles, if they are “functional” (preserve target gene function), can rapidly spread through target populations under strong selection pressure, outcompeting the suppression drive. For example, in a cage trial of the suppression drive targeting *nudel* in *A. gambiae*, functional resistance outcompeted the drive^23^. Even nonfunctional resistance alleles (that will still be recessive female-sterile) can be problematic for homing suppression drives because they reduce the suppressive power of the drive. This was evident in a homing drive targeting female fertility gene *yellow-G* in *D. melanogaster*, where high fitness costs in female drive heterozygotes and high embryo resistance form maternally deposited Cas9, coupled with a modest drive conversion rate, prevented reduction of the cage population due to insufficient genetic load^24^.

To combat resistance, several strategies have been proposed. To address functional resistance alleles, targeting highly conserved and functionally critical sites will more likely result in nonfunctional alleles. The female-specific exon of the sex determination gene *doublesex* is one such target, where mutations frequently disrupt splicing signals and protein translation in females^9,12,20,21^. gRNA multiplexing has also emerged as a promising method to prevent functional resistance. Simultaneous targeting of multiple sites increases the likelihood of large fragment deletions, and the conversion of all target sites into functional resistance alleles is much less likely compared to single gRNA targets^25–27^. A previous study in *Drosophila* expressed four gRNAs targeting well-spread sites within the target gene, though the drive conversion rate was modest and the homing events were unstable^28^. A more recent approach involved transcribing multiplexed gRNAs from a single U6 promoter with closely spaced target sites, achieving higher conversion rates and eliminating functional resistance^24^.

Revised design can eliminate the requirement for high drive conversion if total cleavage is still high^29^, but other issues can still affect genetic load and prevent population elimination. These include undesired Cas9 activity leading to fitness costs, as well as nonfunctional resistance allele formation. One solution is to enhance Cas9 activity specifically in the germline while minimizing its expression elsewhere, thus allowing wild-type alleles to persist in non-germline tissues, which is required for female fertility. Regulating maternal deposition of Cas9 can also reduce formation of resistance alleles in the embryo, which is often a major source of these. A recent study comparing Cas9 promoters in *D. melanogaster* identified *nanos* and *CG4415* as promising candidates for homing drives, the former lacking significant somatic expression and the latter with very low maternal deposition and only modest somatic expression^30^. Yet, fitness costs based on cage studies are still apparent even in drive heterozygote females utilizing the *nanos* promoter, suggesting that the *yellow-G* target gene may be required in certain ovary cells where *nanos*-Cas9 is expressed^24^. Because different genes are required in different tissues with potentially variable Cas9 expression, it may be possible to select alternate target genes that result in substantially reduced fitness costs.

In this study, we selected nine female fertility or viability genes and tested them using split homing gene drives to identify promising candidates with enhanced efficiency. We combined several gRNA lines, each containing four multiplexed gRNAs, with Cas9 lines under different germline promoters to evaluate drive efficiency. The most promising target genes were further assessed in cage studies. Additionally, fecundity and fertility were evaluated through single-pair crosses to investigate the causes of fitness costs. Our findings highlight potential factors limiting suppression drive efficacy and identify promising candidates for the next generation of efficient suppression gene drives.

## Methods

### Target genes selection and protein alignment

In this study, we focused on targeting female-specific genes for population suppression. We utilized the *Drosophila* database (https://flybase.org) to identify genes annotated as mutant female sterile or lethal, with an expression profile biased towards somatic tissues in female adults, ideally excluding germline expression. The gene ID of selected target genes are listed in Table S1.

Homologs of the target genes in several globally important pest species were identified by BLASTing *Drosophila* protein sequences against the NCBI database (https://blast.ncbi.nlm.nih.gov/Blast.cgi) and aligned with ClustalW. These pest species, including *Ceratitis capitata*, *Spodoptera frugiperda*, *Plutella xylostella*, *Aedes aegypti*, *Anopheles gambiae*, *Culex quinquefasciatus*, *Leptinotarsa decemlineata*, *Rhynchophorus ferrugineus*, *Periplaneta americana* and *Blattella germanica*, belong to different insect orders and cause significant damage to human health, endangered species, and agricultural production (Table S2).

### gRNA targets design

For designing gRNA target sites in *Drosophila*, we used the online tool CHOPCHOP (https://chopchop.cbu.uib.no/) to select targets with higher predicted cleavage activity. Four gRNA targets were designed for each gene, primarily targeting the 5’ coding region of essential domains. The exception was *tra*, for which two constructs containing different gRNA sets were generated. This multiplexing approach was intended to fully disrupt the target genes and prevent the formation of functional resistance.

### Expression profiling

Transcriptome data of target genes were downloaded from Flybase to analyze their expression profiles across different developmental stages (Flybase dataset: modENCODE_mRNA-Seq_development) and adult cell types (Flybase dataset: scRNAseq_2022_FCA)^31^. The developmental stages dataset included data from egg, larva, prepupa, pupa, adult male, and adult female, each containing various developmental time points. The adult cell types dataset was derived from single-cell transcriptomes, encompassing various cell types (e.g., neurons, somatic gonad, and germline cells). The downloaded RPKM data was first processed with a log2 transformation, defined as log2(RPKM+1), and then clustered and plotted using TBtools software^32^.

### Plasmid construction

A similar design was employed for constructing gRNA-expressing plasmids targeting different genes. Each gRNA construct contained a DsRed marker under the control of the 3xP3 promoter, a tRNA-linked gRNA cassette driven by *D. melanogaster* U6:3 promoter, and homology arms for homology-directed repair. A step-by-step cloning method, as described in a previous study^24^, was applied, and all plasmids used in this study were constructed via Gibson assembly. The reagents, including restriction enzymes, Q5 polymerases, and Hifi assembly mix, were purchased from New England Biolabs, while oligonucleotides were synthesized by BGI. Constructed plasmids were cleaned up with the ZymoPure Midiprep Kit from Zymo Research for embryo injection and confirmed by Sanger sequencing. All the final plasmid sequences are provided on Github (https://github.com/jchamper/Suppression-Targets).

### Fly rearing and transformation

All flies were reared according to previously described protocols^30^. In brief, flies were fed with artificial cornmeal medium in vials or bottles under 25°C±1°C with a 14/10h day/night cycle. All work with genetically modified insects was performed following regulations approved by the biosafety office at Peking University.

To generate transgenic lines, a mixture of 500 ng/μL donor plasmid and 500 ng/μL *nanos-*Cas9 helper plasmid was injected into *w*^1118^ flies. The embryo injection was conducted by Unihuaii Company. After injection, the founder flies were crossed with *w*^1118^, and their offspring were screened for red fluorescence, indicating successful transformations. Transgenic drive lines were maintained as a mixture of heterozygotes and homozygotes due to the sterility of female homozygotes.

### Fly crosses

For general drive efficiency assessment, drive carriers were crossed to Cas9 lines under the control of either *nanos* (BHDaaN)^33^ or *CG4415* (SNc9XnGr or SNcc9XnG, the latter on chromosome 2L instead of 2R like the other Cas9 genes) promoters^30^. Their offspring, heterozygous for both drive and Cas9, were then outcrossed to *w*^1118^. Each cross vial contained two males and four females unless specifically stated. The generated progeny were phenotyped for red and green fluorescence to assess the inheritance of drive and Cas9 alleles. For the control cross, drive heterozygous flies were crossed to *w*^1118^ in the absence of Cas9. Their offspring were phenotyped for red fluorescence to calculate drive inheritance.

### Cage study

Due to the nature of split drives, drive and Cas9 alleles tend to segregate after crossing. Therefore, to conduct a cage study designed to mimic the release of a complete drive in a wild population, we first integrated Cas9 alleles into the population genomic background and then released the drive into this population (with drive individuals also containing Cas9). Specifically, drive males were outcrossed to Cas9 homozygous females for several generations to generate fly lines heterozygous for the drive and homozygous for Cas9. Males of this line were crossed with Cas9 homozygous females for two days and then removed from the bottle. The mated females, together with females that were mated with Cas9 only males, were equally distributed into eight food bottles for a two-day oviposition period, after which the females were discarded, and the bottles, without plugs, were placed into 25 x 25 x 25 cm cages. Flies were allowed to grow and fly inside the cages. Subsequently, food bottles were replaced with new ones after 12 days, and flies were allowed to lay eggs, which was recorded as the G0 generation. Only one day of oviposition was conducted in the following generations, resulting in a 13-day cycle for each generation. After oviposition, adults were frozen and collected for phenotyping, while egg-containing bottles were returned to the cages to continue the cycle of non-overlapping generations.

### Analysis of cage performance

After collecting drive carrier frequency for each generation of each cage, we employed a maximum likelihood approach to quantify drive fitness costs. This model was similar to the model used in previous studies^24,30,34^, with a simplifying assumption of a single gRNA at the drive allele site. Here, all the target alleles would be converted into non-functional resistance alleles if they could not be converted into drive alleles in the germline (based on relatively high embryo resistance, we assumed that there would be few or no remaining wild-type alleles after Cas9 cleavage). The drive conversion rates and embryo resistance formation rates in males and females were set based on drive efficiency tests in individual vials. The female heterozygote fitness and effective population size were inferred by the model.

### Fecundity and fertility test

Three different cross schemes were conducted to investigate the fertility of drive carriers. The biparental cross mimicked the fertility performance in our cage experiment. Specifically, drive males were outcrossed to Cas9 homozygous females for several generations to produce males heterozygous for the drive and homozygous for Cas9. These males were then crossed to Cas9 homozygous females and their offspring, including drive and non-drive individuals, were outcrossed to Cas9 homozygotes for egg number and egg viability assessment.

The maternal/paternal Cas9 cross schemes were set up by crossing drive heterozygous males/females (without Cas9) to Cas9 homozygous females/males to generate flies heterozygous for both drive and Cas9 alleles or non-drive Cas9 heterozygotes. These flies were then crossed to *w*^1118^, and the egg number and egg viability of their offspring was recorded.

In the above cross schemes, Cas9 homozygotes or *w*^1118^ for each target gene cross were respectively collected from the same parental bottles to minimize the batch effect caused by stochastic factors. Each cross vial contained a single pair of male and female as a biological replicate. Flies were moved to new vials every day, and their egg numbers were counted over three consecutive days. Progeny were reared until emerging as adults and then phenotyped. To assess the embryo resistance allele formation rate, drive females from the drive mothers were randomly collected and outcrossed to *w*^1118^ males to assess their fertility. These females were then sequenced to confirm their genotype.

To reduce batch effects between different individuals, in each cross scheme, drive males, drive females, control males, and control females were all collected from the same parental cross (drive heterozygote crossed with Cas9 homozygote), ensuring they had the same parental effects. Additionally, because all the files were reared in the same bottle before being collected for fertility assessment, the potential effect from various population density and food characteristics was minimized. This means that in the biparental Cas9 cross, the control flies likely had one nonfunctional resistance allele at the target site, and in the paternal/maternal Cas9 cross, the control flies had one copy of Cas9. However, this was considered necessary due to the wide variance in fertility and especially fecundity between flies from different batches. Our target genes were selected to be haplosufficient, and Cas9 alone is unlikely to have substantial fitness effects^35^, so these controls likely had similar performance to wild-type flies.

### Phenotyping and genotyping

Flies anesthetized with CO_2_ or frozen at -20°C were phenotyped. Morphological observations were made using a stereomicroscope. Their fluorescence was confirmed using the NIGHTSEA adapter SFA-GR for DsRed and SFA-RB-GO for EGFP. The percentages of flies carrying DsRed or EGFP were recorded as the inheritance rates of drive or Cas9 alleles, respectively, in single-vial crosses.

To address potential batch effects (each vial was considered an independent batch), potentially biasing rate, and error estimates, we analyzed data as reported in previous studies^24,30^. We fit a generalized linear mixed-effects model with a binomial distribution (maximum likelihood, Adaptive Gauss-Hermite Quadrature, nAGQ = 25). This approach accounts for variance between batches, resulting in slightly different parameters and increased standard error estimates compared to pooling all individual progeny from different vials. The analysis was performed with R program (3.6.1) and supported by the lme4 (1.1-21) and emmeans (1.4.2) packages.

To identify resistance alleles formed in the germline or embryos, drive or non-drive flies were individually ground, and their genomic DNA was extracted using DNAzol (Thermo Fisher) according to the manufacturer’s protocol. A pair of oligonucleotides were designed for each gene to amplify a 210∼600 bp product covering the gRNA target region. This product was gel-cleaned, sequenced by Sanger sequencing, and analyzed with Benchling (https://www.benchling.com/). The oligonucleotide sequences are provided in GitHub (https://github.com/jchamper/Suppression-Targets).

## Results

### Target selection and expression profiling

Nine genes annotated as recessive and essential for female viability/fertility were selected from Flybase as suppression drive targets, including *intersex* (*ix*), *nudel* (*ndl*), *NADPH oxidase* (*nox*), *Octopamine β2 receptor* (*oct*), *stall* (*stl*), *transformer* (*tra*), *virilizer* (*vir*), *defective chorion* (*dec*) and *sex-lethal* (*sxl*). Seven of these genes are located in autosomes while *dec* and *sxl* are on the X chromosome. Among these genes, *sxl*, *tra*, and *vir* are required for sex-determination or dosage compensation, whereas the others play important roles in female-specific development and reproduction (Table S1).

Most of these genes, with the exception of *dec* and *tra*, have homologs across various insect species, including major agricultural pests and disease vectors. The *tra* gene has been identified in *Drosophila melanogaster* and two cockroach species, *Periplaneta americana* and *Blattella germanica*, while *dec* appeared to be restricted to *D. melanogaster* and *Ceratitis capitata*. Several conserved regions were found in these proteins, suggesting conserved functions and that these regions would be potentially suitable for gRNA targeting as part of a pest control strategy across diverse insect species (Table S2).

We analyzed the potential functions and expression profiles of these genes at different developmental stages and in adult cell types (Table S1 and Figure S1). Transcriptome data from Flybase shows that *ndl*, *dec*, and *stl* share a similar expression profile, being highly expressed in female adults. This is consistent with their critical roles in eggshell formation, maternally affected egg patterning, and ovarian follicle development. *ix*, *sxl*, *tra* and *vir* are clustered together, with expression spread throughout developmental stages and slightly higher in early and middle-stage embryos, likely due to their significant roles in sex determination and differentiation. Most of these genes show higher expression in female adults than male adults, except for *oct*, which exhibits slightly higher expression in males. Although *oct* is important for female ovulation and fertilization, its expression is observed from the late embryo stage, peaks at the late pupa stage, and decreases at the adult stage. In adults, *ndl*, *nox*, *tra*, *ix*, and *vir* show similar expression patterns, while *oct*, *stl*, *dec* and *sxl* are clustered together. At the cellular expression level, *ndl*, *nox*, *tra*, *ix*, and *vir* exhibit similar expression patterns, while *oct*, *stl*, *dec*, and *sxl* display a comparable pattern. In general, these genes have lower, but not absent, expression in female and male germline cells compared to other somatic cell types. Although the expression profiles suggest potential roles of these genes in males, their primary functions are likely more pronounced in females.

### Drive construct design and transformation

In this study, gRNA constructs were knocked into the female fertility genes to generate drive lines, which were then combined with existing Cas9 lines in a “split drive” design. Each gRNA construct consists of two homology arms flanking target sites for homology-directed repair, a DsRed marker driven by the eye-specific 3xP3 promoter for phenotyping, and a set of four gRNAs under the control of U6:3 promoter (Figure 1A). To avoid potential recombination caused by repetitive genomic promoter sequences of multiple U6-gRNA cassettes, tRNAs were placed at the start of the gRNA transcripts and between gRNAs, allowing gRNAs to be expressed in a single transcript and spliced out to create mature gRNAs. The purpose of this gRNA multiplexing design was to prevent formation of functional resistance alleles, and also to potentially compensate for low-efficiency gRNAs, thus increasing cut rates and drive conversion efficiency.

**Figure 1.**
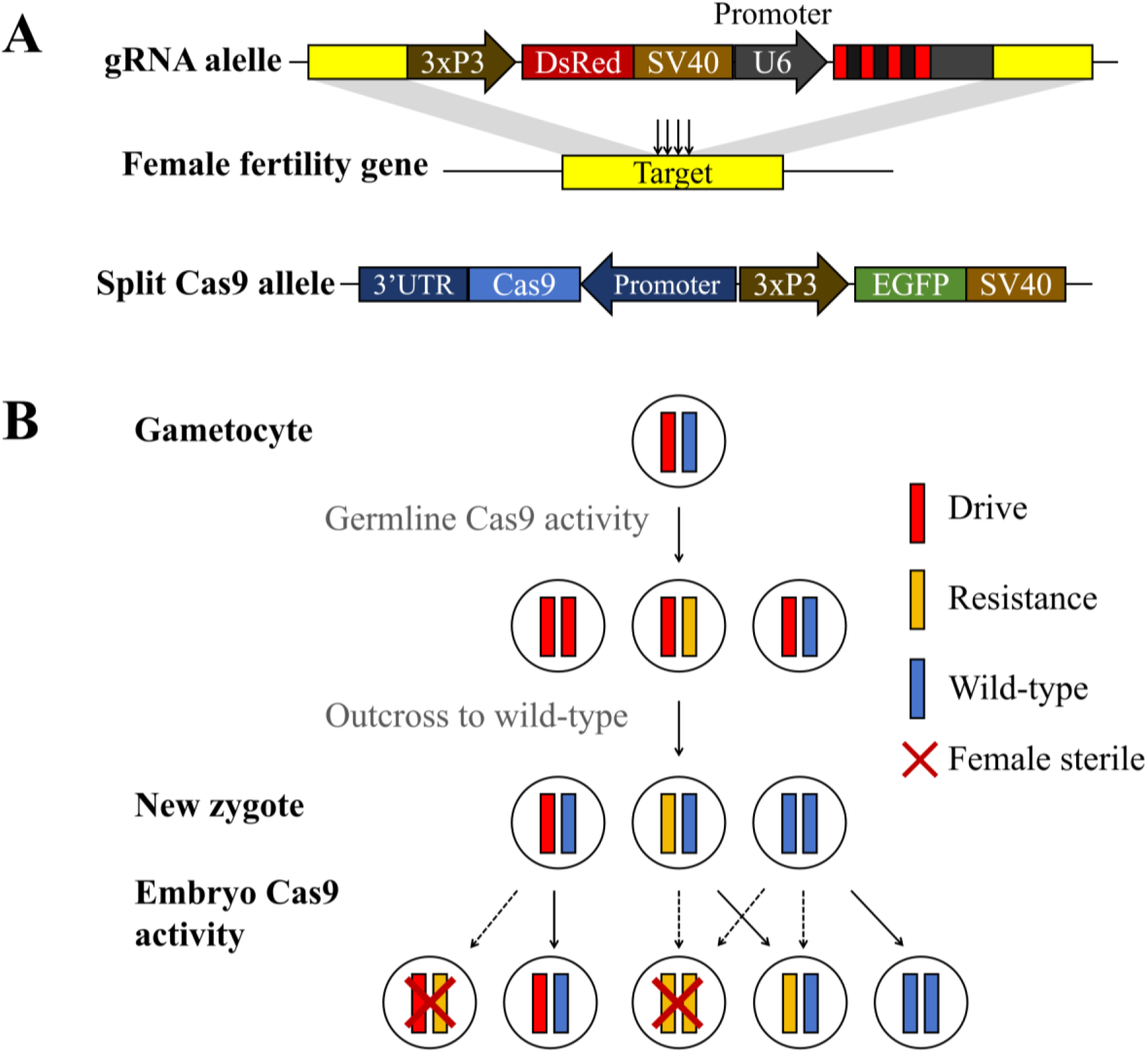
Female fertility homing suppression drive construct design and mechanism. (**A**) The gRNA (drive) construct was inserted into a female fertility gene, containing the 3xP3-DsRed-SV40 marker and a set of 4-gRNA under the control of U6 promoter to target the insertion site. A split Cas9, under the control of a germline promoter (*nanos* or *CG4415*), was provided at an unlinked site together with 3xP3-EGFP-SV40. (**B**) In the germline, wild-type alleles are cut by Cas9/gRNA and converted into drive alleles by homology-directed repair or resistance alleles via end-joining. Maternally deposited Cas9 and gRNA can also cut and disrupt the wild-type allele in early embryos. Females carrying any combination of drive or non-functional resistance alleles will be sterile, while a single copy of wild-type is sufficient to preserve female fertility. However, somatic Cas9 express can partially or completely convert the wild-type allele to a drive or resistance allele in drive/wild-type heterozygotes.

In these drives, target sites are cleaved, and subsequent homing or resistance allele formation occurs in germline cells in the presence of gRNA and Cas9 (after crossing gRNA lines to Cas9 lines), while targets in somatic cells ideally remain intact. When flies carrying both gRNA and Cas9 are crossed with wild-type, their offspring inherits either the gRNA “drive” allele, a disrupted resistance allele, or a wild-type allele from the drive parent. However, these heterozygotes or wild-type homozygotes could also undergo additional cleavage from the maternally deposited Cas9, converting any wild-type alleles (including from the non-drive parent) into resistance alleles. In drive heterozygote individuals with Cas9, it may be possible for additional cleavage to occur in somatic cells, depending on Cas9 expression. Because null alleles for these target genes are recessive, females lacking wild-type alleles (carrying only drive and/or nonfunctional resistance alleles) will be sterile (Figure 1B).

To ensure the disruption of gene function in resistance alleles, two specific criteria were adopted. First, all gRNA target sites were chosen either at the start or in the middle of essential conserved domains (Figure 2) to reduce the chance that resistance alleles lacking frameshifts were functional ^36^. Additionally, multiplexing gRNAs were applied to further reduce functional resistance alleles generated in a single site. The four gRNA sites are placed closely together, with the length between two outer gRNAs no larger than 128 bp, to maximize drive conversion rates^25^. Note that two gRNA constructs (*tra-v1* and *tra-v2*) were generated to target *tra* with different gRNA sets, while only one construct was designed for each of the other genes.

**Figure 2.**
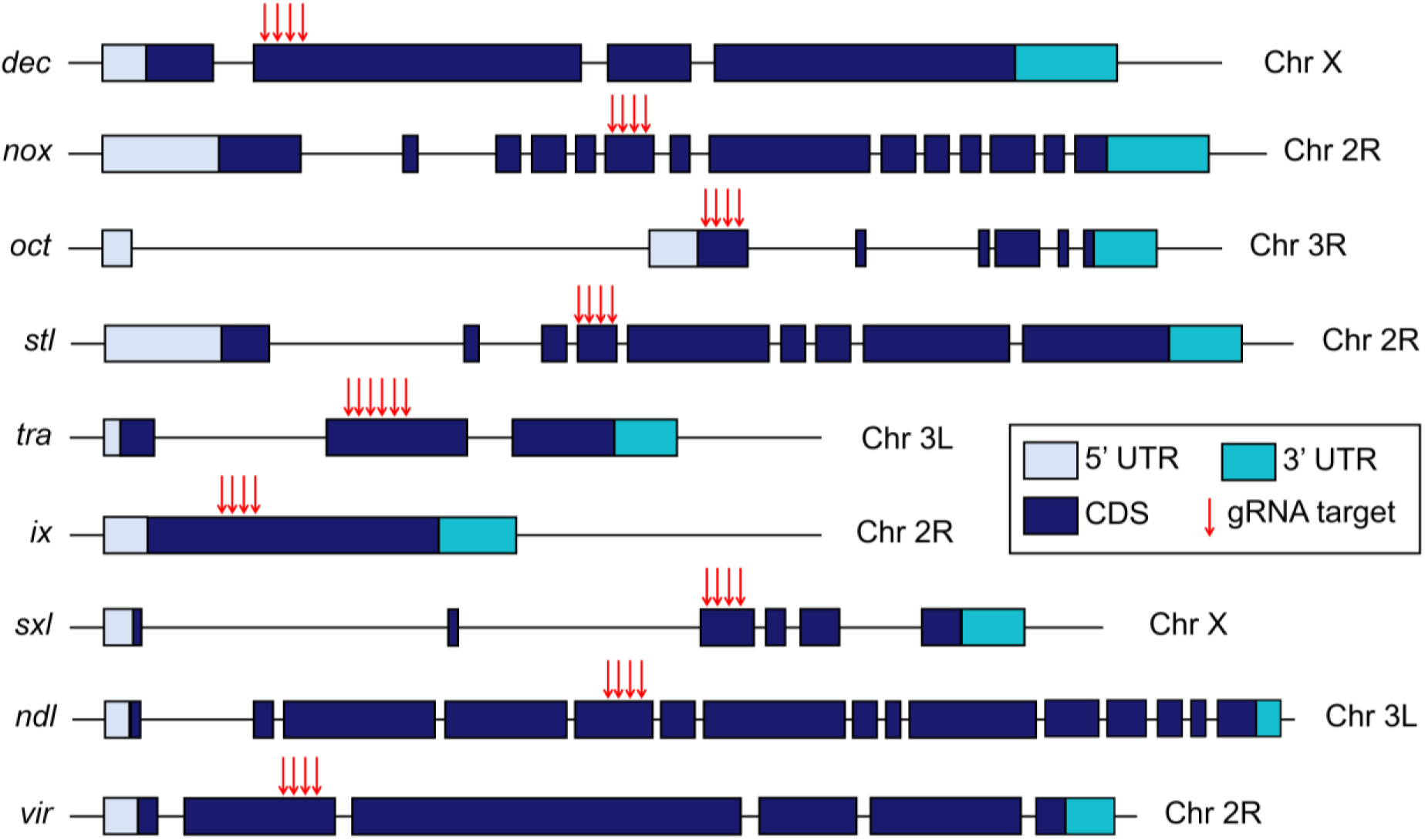
Genomic structure of target genes and gRNA target sites. The untranslated region (UTR) and coding sequence (CDS) are shown with colored rectangles, while introns and intergenic regions are shown with black lines. gRNA target sites are indicated with red arrows. Gene elements are not drawn exactly to scale.

We inserted the drive construct into the target site of each gene and generated transgenic lines through embryonic injection. Drive heterozygotes and homozygous flies, confirmed with genotyping, were randomly collected and respectively crossed to *w*^1118^ files to assess their fertility. The results showed that all drive heterozygotes, as well as male homozygotes with drives targeting *nox*, *oct*, *stl*, *ndl*, and *ix* were fertile, while female homozygotes for these drives were sterile, consistent with expectations based on FlyBase annotations. Notably, we were unable to generate viable male or female homozygous adults for the drive targeting *vir*, nor were we able to generate any female homozygotes for *ix*, suggesting potential lethality during early developmental stages of these homozygotes. While female lethality is not likely to substantially affect suppression effectiveness, male homozygous lethality would reduce suppressive power similarly to X-linked targets (and haplodiploid species).

### Drive efficiency assessment

To assess the drive efficiency, flies heterozygous for both drive and Cas9 alleles (receiving at least one of these alleles from a male parent) were crossed to the *w*^1118^ line, after which the drive inheritance in their offspring were recorded (Figure 3, Data Set S1). The results showed varied drive performance targeting different genes. When combined with *nanos-*Cas9, the drive inheritance rates were usually higher than the 50% Mendelian expectation, except for drive females targeting *vir* and *sxl*, which were slightly lower than 50%. This could be explained by potential fitness costs from the drive insertion at these two genomic sites, or potentially an important function of these two genes in the germline, where they will be disrupted by the drive or resistance allele. The drive lines targeting *oct* and *stl* showed the highest drive inheritance rates in both females (93.2%±0.8% and 94.0%±1.0%, respectively) and males (91.0%±2.0% and 89.2%±1.1%, respectively). The drive inheritance rates of constructs targeting *ndl* and *nox* were moderate, achieving around 84-88%. When targeting *tra*, different drive inheritance rates were observed between two constructs with different gRNA sets (61%-66% for *tra-v1* and 81%-83% for *tra-v2*), possibly due to varied gRNA cleavage efficiency. Notably, when targeting *ix*, the fertility of drive females was greatly reduced, and fewer offspring were produced, likely due to the fitness cost from embryo activity of Cas9/gRNA forming recessive lethal alleles. Because both *dec* and *sxl* are X-linked genes, we only assessed drive conversion in females. However, only the drive line targeting *dec* showed a biased drive inheritance in their offspring (73.3%±2.0%).

**Figure 3.**
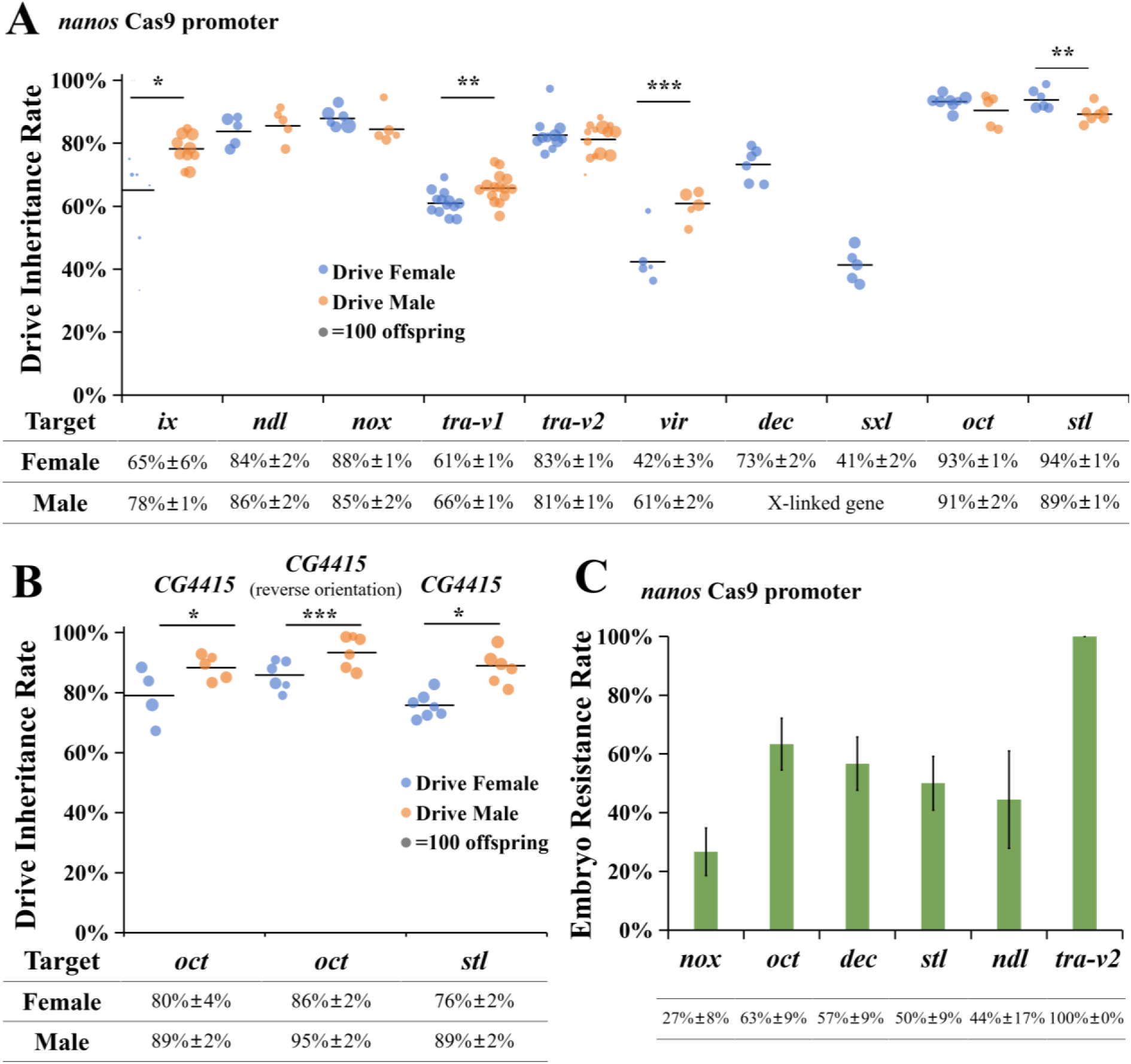
Drive inheritance rates with different combinations of drive and Cas9 lines. (**A**) Drive inheritance with Cas9 driven by the *nanos* promoter. (**B**) Drive inheritance with Cas9 driven by the *CG4415* promoter. *CG4415* (reverse orientation) indicates that Cas9 has the reverse orientation relative to the fluorescence marker. It is at nearly the same site as the *nanos*-Cas9 drive (chromosome 2R), but the other *CG4415*-Cas9 is on chromosome 2L. Drive inheritance rate is the percentage of offspring with DsRed fluorescence from crosses between drive heterozygotes (which are also heterozygotes for Cas9) and *w*^1118^ flies. Each dot represents the offspring from a single vial, and the size of dots is proportional to the total number of offspring. Drive inheritance rates between males and females in each target gene are compared. \**p<*0.05, \*\**p<*0.01 or \*\*\**p<*0.001 (z test). (**C**) Embryo resistance rate indicates the proportion of sterile drive female offspring derived from a mother that had the drive.

As a negative control, we examined the inheritance rates of drives by crossing heterozygous drive carriers with *w*^1118^ flies in the absence of Cas9. Generally, this resulted in no bias in drive inheritance, except for the drive targeting *dec*, which was somewhat lower than the 50% Mendelian expectation (41.3%±2.6%, *p=*0.0012, z test) (Data Set S2). This outcome was likely due to a fitness cost associated with the insertion of the drive construct or haploinsufficiency of *dec* in drive carriers.

Our previous study showed that the *CG4415* promoter combined with the *nanos* 3’UTR performed better with a suppression drive targeting the female fertility gene *yellow-G*, achieving slightly higher drive conversion and much lower embryo resistance allele formation than when Cas9 used *nanos* regulatory elements^30^. To assess if this was consistent across different targets, we respectively combined the *oct* and *stl* drive lines with the same two Cas9 lines (at nearly the same genomic insertion site) using the *CG4415* promoter and *nanos* 3’UTR, one of which has the orientation of the Cas9 gene reversed. When targeting *oct*, both *CG4415-* Cas9 lines showed significantly lower though still fairly high female drive conversion rates compared to the *nanos-*Cas9 line (*p<*0.0001 and *p*=0.003, z test), although male drive conversion rates were not affected (*p=*0.8098 and *p=*0.3797, z test). While targeting *stl*, a lower drive conversion rate was observed in females (*p<*0.0001, z test) but not males (*p*=0.8665, z test). These are consistent with the finding in the previous study that suggested somewhat lower germline expression of *CG4415*-Cas9 than *nanos*-Cas9^30^, but support the potential use of these promoters with our new drive lines.

We also compared the drive performance between drive males and females in each group. The results showed higher drive conversion rates in males than females for *ix*, *tra-v1*, and *vir* with *nanos-*Cas9. This is consistent with our previous study targeting another female fertility gene *yellow-G*^24^. A similar trend was also observed for drives targeting *oct* and *stl* when Cas9 was under the control of *CG4415* promoter. However, when targeting *stl* with *nanos-*Cas9, more offspring inherited drive alleles from drive females than males. Because the drive inheritance rates in *stl* drive males and drive females were both high, the observed difference may have been caused by random factors in test vials. In addition, no significant differences were found between drive males and drive females in other groups (Data Set S1).

### Resistance allele formation

Embryo resistance formation rates of drives were also assessed by crossing drive females to non-drive males (Figure 3C). Because mutation of most target genes does not show observable phenotypes, the measurement of embryo resistance of these lines was based on the percentage of sterile drive females derived from drive mothers (which should be caused by embryo resistance in nearly all cases, though this might produce a slight overestimate due to natural sterility). The embryo resistance rates of drives targeting *oct*, *dec*, *stl* and *ndl* were moderate to high, (63.3%±8.8%, 56.7%±9.1%, 50.0%±9.1%, 44.4%±16.56%, respectively), while the rate of *nox* drive was relatively lower (26.7%±8.1%). Sequencing showed mutations in targets in all of these sterile females.

The exception was for the *tra-v2* drive, which as noted, had 100% sterility. When *tra* drive only (without Cas9) flies were intercrossed, phenotyping and genotyping showed that drive homozygous females were morphologically male and sterile, while phenotypes of heterozygous males and females were identical to *w*^1118^ flies. Notably, in the offspring of *tra* drive female crosses, more male-like flies were observed than females (Data Set S1). Some of these male-like flies exhibited slightly bigger body sizes like wild-type females, and they were genetically females based on genotyping. These flies were recorded as “strong masculinization”. Additionally, there were some flies displaying an intermediate phenotype between male and female (patchy dorsal fragment), which were recorded as “mild masculinization” (Figure S2). Ten genetically female flies with the drive and strong masculinization and mild masculinization phenotypes were individually crossed to *w*^1118^, and none of them produced offspring, suggesting sterility. Sequencing confirmed the presence of embryo resistance alleles in them, though wild-type was always present as well for the ones with mild masculinization phenotype. Note that these masculinized flies represented all the genetically female offspring of drive female crosses, with the small number from male drive crosses (with a strong masculinization phenotype) likely misidentified due to the difficulty of separating strong masculinization females and wild-type males.

Additionally, the fraction of fertile drive female controls (derived from drive father) was 90-100% (Data Set S5). Therefore, the sterility observed in the above lines was most likely caused by embryo resistance. However, note that moderate or heavy mosaic embryo resistance could still cause sterility, so the rate of “complete” embryo resistance (mutations in all cells due to cleavage in early embryo cells) is likely lower than our reported values. This is particularly apparent in the *tra* drive, in which only the stronger masculinization phenotype had any chance of being a complete embryo resistance allele. As a performance measurement, this is therefore not entirely complete. Some females that remain fertile may still have enough mosaic cleavage to have reduced fertility. On the other hand, males that experience enough mosaic cleavage to render females sterile may be unaffected and still be able to perform normal drive conversion in the germline^33^.

### Suppression drive cage experiments

We selected some of the drive lines showing superior performance in small-scale drive efficiency tests (*nox*, *oct*, and *stl*) and further assessed them in cage experiments. To set up cages, lines heterozygous for drive allele and homozygous for Cas9 allele were generated and released into cage populations containing a homozygous Cas9 background. The drive carrier frequency and total fly number of each generation were tracked. The drive carrier frequency was expected to increase over several generations, reaching high equilibrium frequency and causing population suppression after forming high numbers of sterile female homozygotes.

Five cages were set up in total, including one cage combining *nanos-*Cas9 with the drive targeting *oct*, one cage containing *nanos-*Cas9 and the drive targeting *nox*, two replicate cages containing *nanos-*Cas9 and the drive targeting *stl*, and another cage containing *CG4415-*Cas9 and the drive targeting *oct*. Most of these cages were set up with a drive carrier frequency of 18%-24%, except for one of the *stl* cages (cage 1), which started at 59% drive carrier frequency (Figure 4, Data Set S3).

**Figure 4.**
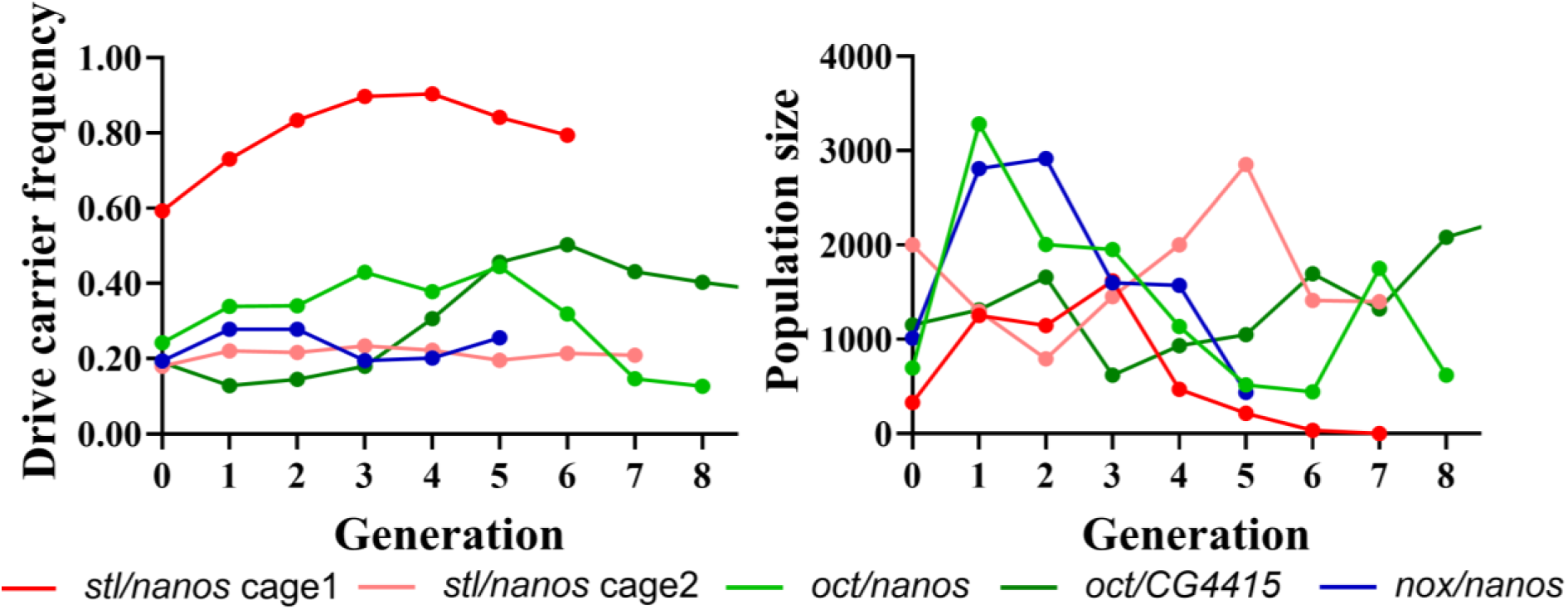
Drive carrier frequency in cage populations. Flies carrying one copy of the drive and two copies of Cas9 were released into Cas9 homozygous populations. All flies in each discrete generation were phenotyped. In the legend, the first gene is the drive target, and the second is used for Cas9 promoter and 5’ UTR regulatory elements. See Data Set S3 for later generations of the *oct*/*CG4415* cage.

In cage 1 for *stl*, the drive carrier frequency increased to 90% by the fourth generation and then was slightly reduced in subsequent generations. During this time, the population size expended at first but then quickly dropped after the third generation, with complete population elimination by the seventh generation. Though the drive carrier frequency was only at a moderately high level, nonfunctional resistance alleles likely also contributed to the fraction of sterile females. However, when released at a lower frequency in cage 2, the drive frequency remained approximately constant at ∼20%, with no apparent effect on the population size (Figure S4). The *nox* drive had a similar performance.

In the cage with *nanos-*Cas9 and drive targeting *oct*, the drive carrier frequency marginally increased but did not reach a high level. The population size of this cage fluctuated, but ultimately was not eliminated by the drive. When combined with the Cas9 element driven by *CG4415* promoter, the *oct*-targeting drive carrier frequency gradually rose up to 50%, after which it slowly declined to approximately 33%, with the population size apparently unaffected (Data Set S3).

Our result indicates that with a higher initial release ratio, the cage population could be completely eliminated. However, this suppression capability was diminished when the release ratio was reduced and the population size increased. Most likely, the high release frequency drive, even though its frequency would perhaps have declined in the long run to a lower equilibrium, was able to remain high for long enough to place a temporary high genetic load on the population, allowing its elimination. Allele effects may have contributed to this after the population size was substantially reduced (when too few flies laid eggs on the bottles, bacterial or fungal growth would often render larvae nonviable). Despite achieving a drive conversion rate of 85% to 93% in drive heterozygotes (Figure 3), the equilibrium frequency of drive carriers in our cage populations was lower than anticipated based on a previous modeling study^25^. This suggests a fitness cost in drive carriers in our cage populations, particularly the *CG4415* cage with its likely low embryo resistance, leading to failure to eliminate the population.

### Maximum-likelihood analysis of fitness in cage populations

To assess the drive performance of our cages, a maximum-likelihood method was applied to quantify fitness components^24,30,37^. This model utilized a simplified design with a single gRNA and assumed no functional resistance, consistent with the expected outcome when using multiplexed gRNAs. This is because even if functional resistance forms at one or more target sites, the remaining sites can still be cleaved and disrupted, significantly reducing the likelihood of functional resistance. Drive conversion and embryo resistance allele formation were set as per our previous measurements (embryo resistance in the *CG4415* cage was set to zero, because it was lower than 5% even with a highly active gRNA element^30^). Fitness costs were assumed to occur in female drive heterozygotes due to cleavage of wild-type alleles where they were needed in certain cells (either somatic or germline cells). The parameters used for modeling can be found in Table S3.

The estimated fitness of drive females in cages targeting *nox* and *oct* with the *nanos* promoter were 0.262 and 0.245, respectively. When combined with *CG4415* promoter, the cage of drive targeting *oct* exhibited a slightly higher fitness of 0.318. For the drive targeting *stl* with the *nanos* promoter, the inferred fitness was 0.205 in the cage with a lower initial release ratio. However, in another successfully suppressed *stl* cage with a higher initial release size, fitness was improved to 0.681, with the upper limit of the 95% confidence interval reaching 1.065. The reason for this difference is unclear, but may be related to the specific conditions of the cages. Alternatively, the improved performance and fitness in the *stl* cages might be due to a relatively smaller population size (Data Set S3), causing reduced competition among larvae.

The failure of the four cages either with *nanos* or *CG4415* promoter could perhaps be attributed to low fertility of drive females within these cage populations. However, an important caveat is that our maximum likelihood method would probably have inferred a low fitness even if the nature of the fitness cost was different^37^.

### Fecundity and fertility assessment

To confirm whether there was a fitness cost in drive females, we conducted single-pair crosses to assess individual fecundity and fertility. A specific cross scheme was designed to mimic the cage performance of drives (Figure S3, biparental Cas9 cross, Data Set S4). In the founder generation, males drive heterozygotes that were homozygous for Cas9 were crossed with Cas9 homozygous females. Their offspring, including drive heterozygotes and non-drive flies (which likely had a single nonfunctional resistance allele resulting from a high germline cut rate in the drive parental germline with *nanos-*Cas9^30^), were collected for single pair crosses. Note that these offspring contained two copies of Cas9. In comparison, we crossed drive flies (without Cas9) with Cas9 homozygotes in the founder generation (Figure S3, maternal/paternal Cas9 cross, Data Set S5 and S6). In these crosses, offspring contained only one copy of Cas9, and a single resistance allele might be formed in the maternal cross but absent in the paternal cross. These drive and non-drive offspring were individually crossed to non-drive flies (Cas9 homozygotes in the biparental Cas9 cross and *w*^1118^ flies in maternal/paternal Cas9 crosses), and their offspring were counted to assess parental fecundity and fertility.

For fecundity assessment, there are often large differences between fly batches (compare cross schemes, which were separate experiments), but differences in flies originating from the same vial were more likely due to drive effects (flies would experience identical rearing conditions in the same vial). Egg numbers laid per day were counted over the course of three days (Figure S4). In the biparental Cas9 cross, *nox* drive females laid more eggs per day (26.92±2.17) compared to the non-drive female controls (18.20±1.57) (*p*=0.0026, t test). No significant difference was found in the fecundity between drive carriers and non-drive control in the drives targeting *oct* and *stl* (Data Set S4). In comparison, no significant differences were found between the drive and non-drive flies in all groups for the maternal Cas9 cross and most groups foor the paternal Cas9 cross (Data Set S5 and S6). It is noted that in the paternal Cas9 cross with the drive targeting *nox*, non-drive females laid more eggs per day (26.67±2.96) than drive females (18.37±1.60) (*p*=0.0150, t test), which is contrast to the results of biparental and maternal Cas9 crosses. Besides possibly *nox*, however, we did not see any substantial fecundity effects that were likely caused by drive effects. The reasons for these differences are unclear and may be due to epigenetic effects from parents or grandparents reared in different environments. Random effects also cannot be ruled out.

To further assess fertility, egg-to-adult survival rates were determined from the same experiments (Figure 5). In the biparental Cas9 cross, the egg-to-adult survival rates of drive females were lower than non-drive females in both *nox* and *oct* groups (both *p<*0.0001, z test), indicating that drive activity likely disrupted these genes in cells that were important for reproduction. In the *stl* group, both drive males (*p*=0.0319, z test) and drive females (*p*=0.0025, z test) showed less egg viability compared to their non-drive controls. Considering that *stl* is essential for female fertility, which should not affect male reproduction, the reduced male egg viability was likely caused by stochastic factors existing in Cas9 population or in the test vials (the marginally significant result would not be considered significant after multiple-testing correction). These results suggest that drive female heterozygotes in the drives targeting *nox*, *oct*, and *stl* in the cage populations suffered moderate fitness costs, reducing the genetic load of the drives and contributing to the failure to suppress the target populations. In the maternal cross, significantly lower egg viability was detected in drive females targeting *nox* (*p<*0.0001, z test), *oct* (*p<*0.0001, z test), *stl* (*p<*0.0001, z test) *tra-v2* (*p=*0.0002, z test) and *ndl* (*p=*0.0009, z test). Most of these results were in line with the biparental cross, while the egg viability loss in *stl* drive males in the biparental cross was not found in the maternal cross.

**Figure 5.**
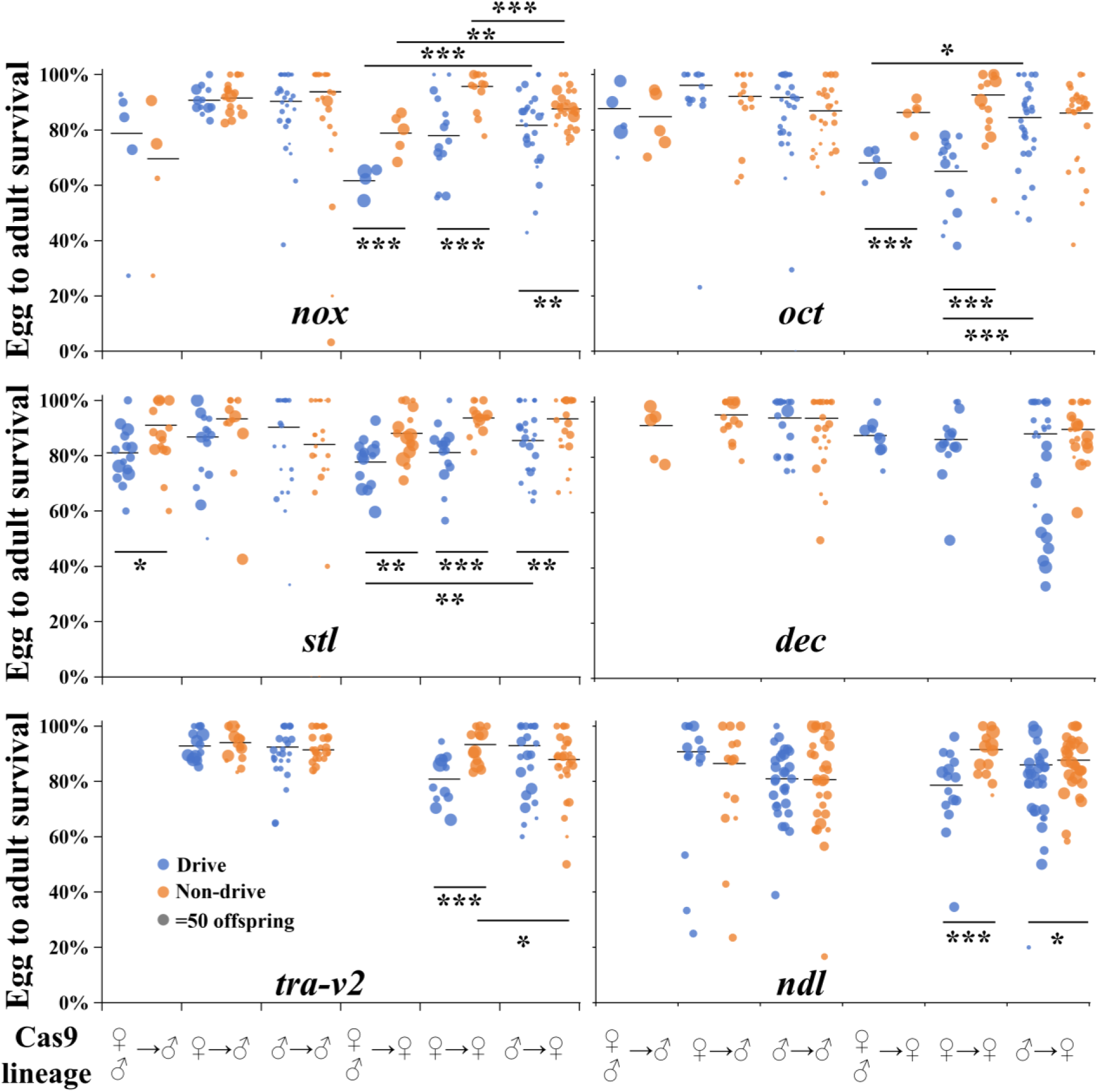
Fertility assessment of drive carriers. Three cross schemes were applied, as shown by Cas9 lineage. For male Cas9, drive heterozygote females (without Cas9) were crossed to males that were homozygous for Cas9. For female Cas9, drive heterozygote males were crossed to Cas9 homozygous females. For biparental Cas9, crosses were made between male drive heterozygotes and wild-type females at the drive site, but both parents were homozygous for Cas9. Drive and non-drive progeny from these crosses were then crossed to non-drive flies, and the offspring were phenotyped. Biparental Cas9 data cross data was not collected for *tra*-v2 and *ndl*, and only partially collected for X-linked *dec*. Drive and non-drive flies are marked with dots in different colors. Significant difference is indicated with \**p<*0.05, \*\**p<*0.01 or \*\*\**p<*0.001 (z test).

These fitness costs could have two potential sources. One is due to the cleavage of target genes in these drive females from newly expressed gRNA and Cas9 (which should be mostly germline-confined for *nanos*-Cas9). However, Cas9 deposition from the maternal founder in both biparental and maternal Cas9 crosses, coupled with newly expressed gRNA, may have led to Cas9 cleavage in late embryo cells, which could affect somatic cells later in development and cause significant fitness costs. Note that while this phenomenon is already incorporated in our embryo resistance studies (in which both Cas9 and gRNA were maternally deposited), it could also occur in split drives with a Cas9 mother and gRNA father, causing fitness costs even though resistance alleles do not form in the zygote and early embryo (gRNA requires time to express).

In comparison, in the paternal Cas9 cross, the egg-to-adult survival rates of drive females in *nox*, *oct*, and *stl* groups were all increased, though the difference between drive and non-drive females still exists in the *nox* and *oct* groups. This may be because only one Cas9 copy exists in these flies. Besides, given that these target genes are also expressed in germ cells at some level, they may also play some roles in those cells (Figure S1B). Therefore, some fitness costs in our drives could still be caused by germline cleavage, which is necessary for drive activity. Although the *nanos* promoter was known to have undetectable somatic activity^30,33^ based on phenotyping assays and sequencing, some somatic activity may still be present, leading to fitness costs.

Note that the maternal and paternal cross setups, with only one copy of drive and Cas9, more closely resembles a canonical complete drive design, making it more reliable in reflecting the fitness of drive females that inherit a drive from their mother or father. Therefore, we only applied these crosses to assess the fecundity and fertility of two additional drives, *tra-v2* and *ndl*, which showed good drive performance in individual vials. The results showed that the egg viability from drive carriers did not differ from non-drive controls in the *tra-v2* group, consistent with the result of *nox* and *oct* groups. When targeting *ndl*, fewer eggs survived in drive females compared to non-drive females, while the egg viability did not vary in male crosses. A similar pattern was also observed in drives targeting *stl* and *dec*. These results indicate that Cas9 and gRNA expression from drive alleles likely affected the fitness of heterozygous drive females targeting *stl*, *dec*, and *ndl*. We also noted lower fitness in non-drive *nox* females from biparental than paternal cross, suggesting potential haploinsufficiency (individuals from biparental cross likely were drive/nonfunctional resistance allele heterozygotes).

## Discussion

Gene drives hold significant promise for population suppression, targeting applications such as disease transmission control and mitigating agricultural losses. The most effective suppression strategies typically involve disrupting sex-specific viability or fertility-essential genes, which gradually eliminate target populations over several generations. In this study, we evaluated nine female fertility genes using a split homing drive system in *Drosophila melanogaster*. While the drives demonstrated varying levels of efficacy, the *stl*- and *oct*-targeting drives achieved the highest drive conversion rates, with the *stl* drive successfully leading to complete population suppression in one of the cage populations. Further analyses revealed that fitness costs, driven in part by maternal Cas9 effects that would be present only in split drive testing, significantly impacted drive efficiency.

The genetic load is an important parameter for understanding the suppression power of a drive. A target population will only be eliminated if the genetic load is high. Factors influencing genetic load include drive conversion efficiency, fitness costs, and nonfunctional resistance allele formation^27,38^. For suppression drives, null mutants resulting from drive or resistance alleles are ultimately removed from the population due to the sterility or lethality conferred by the disrupted target gene, contributing to population sterility, but also slowing the drive by removing drive alleles in females and blocking drive conversion in males. High drive conversion rates are critical to ensuring sufficient genetic load, since this produces more offspring carrying drive alleles in the next generation. Our drives targeting *oct* and *stl* achieved a high drive conversion rate (approximately 80%), though still falling short of the >90% rates observed in comparable *Anopheles gambiae* systems, where suppression was achieved in a few generations despite moderate fitness costs^12^. This discrepancy may explain why some of our cage populations were not eliminated. Optimizing drive conversion efficiency requires strong germline Cas9 activity, perhaps with appropriate timing, to enhance homology-directed repair.

Fitness costs associated with drive carriers pose significant challenges to achieving suppression, especially when nonfunctional resistance allele formation is high (as we saw with *nanos*-Cas9 in many of our drives). We observed reduced egg viability in female drive heterozygotes when Cas9 was maternally provided (even without maternal gRNA), but this fitness loss was mitigated when Cas9 was paternal. Somatic Cas9 expression is another source of fitness cost in drive females because a lack of functional female fertility genes in somatic cells will directly impact female fertility^39^. However, the *nanos* promoter used in this study has been shown to exhibit minimal somatic activity^24,30^, reducing the possibility of somatic cleavage induced fitness costs in female drive heterozygotes. Additionally, target genes used in this study are expressed and probably function in germ cells at various levels, despite our attempts to find suitable targets that were not required in the germline. Thus, loss of these essential genes in reproductive tissues could also impact reproductive egg viability and egg development.

Regulating Cas9/gRNA expression is important for both drive efficiency and fitness. Because strong gRNA is usually only achieved with ubiquitously expressed Pol III promoters, choosing strong germline-specific Cas9 promoters is necessary for avoiding fitness costs from somatic expression in suppression drives. These promoters should have high germline conversion capacity to promote the inheritance of drive alleles and no embryo/somatic activity to avoid fitness costs. Our previous works showed that both *nanos* and *CG4415* have high drive conversion rates^30^, but *nanos* failed to suppress target populations in a homing drive targeting the female fertility gene *yellow-G* due to its fitness cost in drive females^24^. *CG4415* had much lower embryonic activity, which allowed the elimination of cage populations by targeting *yellow-G*^30^. Here, we tested both promoters with drives targeting *oct* and *stl*, showing slightly higher drive efficiency than the drive targeting *yellow-G* in small-scale crosses. However, most of them were unable to suppress cage populations with low release frequency. Computational analysis of cage performance indicated large fitness reductions in drive females, and the follow-up fecundity/fertility tests revealed that maternal Cas9 effects may explain some of these costs in cage experiments with *nanos*-Cas9. Because *CG4415*-Cas9 showed less maternal deposition but more somatic expression in the previous study, its fitness cost in cage populations may be a combination of somatic and germline cleavage effects, rather than a combination of maternal and germline cleavage effects like *nanos*-Cas9.

While maternal deposition poses a persistent challenge in gene drive systems, its impact is particularly pronounced in the context of split drives compared to complete drives. In our cage tests designed to simulate the spread of a complete drive, homozygous Cas9 is integrated into the background population, and gRNA carriers are subsequently introduced. This setup ensures that Cas9 is always present for maternal deposition and cleavage with newly expressed gRNA, contributing to fitness costs in drive daughters. These fitness costs arise even when drive females are derived from a drive father because Cas9 will still be deposited by a non-drive mother. In this case, switching the positions of Cas9 and gRNA (i.e., locating Cas9 in a fertility target gene and gRNA in an un-linked allele) may improve the performance of drive in cage populations because only gRNA will be integrated into the population background. This consideration may also be important in tethered drive^34,40^, with an extra complexity of applying different endonucleases^41,42^ at the two sites if the confined drive is also CRISPR based. Any remaining fitness costs are likely to be more representative of a complete drive. However, because the gRNA line is specific to the target site, multiple testing of several Cas9 promoters with different split drive targets would require a larger number of transgenic constructs. Note that for Cas9 systems with minimal maternal deposition, these considerations do not apply.

The better performing *stl*/*nanos* cage with the higher release size showed higher fitness, which may have been improved by less competition among larvae in a lower-density population. Additionally, higher release ratios hasten the spread of drive, quickly generating more sterile females and crashing the population, perhaps before the drive slowly declined to its natural equilibrium frequency. This indicates that for a less ideal gene drive (with high drive conversion rates but low fitness), increasing release size could help ensure a favorable outcome, at least if the release is widespread. The release size will still usually be smaller than self-limiting strategies^43,44^.

Previous work showed that drive efficiency could be affected by different genomic positions, where Cas9 or gRNA expression pattern is possibly affected by regulatory elements or other features surrounding the insertion position^45–47^. This could also affect gRNA expression, potentially affecting the performance of our drives. Additionally, our two *tra*-targeting drives, differing only in gRNA sets but sharing almost the same insertion site, displayed markedly different efficiencies, emphasizing the critical role of gRNA activity in target cleavage efficiency.

For drives exhibiting low to moderate drive conversion efficiency at fertility target sites, alternative gene drive designs can be employed to enhance suppression efficacy. One promising approach involves a two-target suppression system, as we previously reported^29^. In this design, drive conversion occurs at a distinct genomic site, while separate gRNAs target female fertility genes. This allows for a genetic load that is determined by the sum of the end-joining and homology-directed repair rate at the female fertility site, rather than only homology-directed repair-based drive conversion at the drive site in a standard suppression drive. By decoupling the drive element from the fertility target, this system does not require high drive conversion rates to achieve suppression, only high total mutation rates. However, this system is still vulnerable to embryo resistance and fitness costs and thus still requires suitable target sites and Cas9 promoters, even if some requirements are relaxed.

In conclusion, this study provides insights into the use of female fertility genes for suppression gene drive systems. Maternal Cas9 effects emerged as a key limitation, reducing genetic load and hindering suppression in cage populations. Future efforts should prioritize reducing maternal deposition while retaining low somatic expression and otherwise minimizing fitness costs. Target genes used in this study are functionally conversed and essential for female fertility, holding potential to be used in suppression gene drives in non-model insect species. By addressing these challenges, gene drives can be further refined for effective population control in a broader range of ecological and pest management contexts.

## Supporting information

Supplementary Data Sets

Supplementary Alignment Files

## Acknowledgements

This project was supported by grants from the National Natural Science Foundation of China (32302455 and 32270672) and laboratory startup support from Peking University and the Center for Life Sciences. We thank Sam Champer for assistance with batch effect analysis.

## Supplemental Information

**Figure S1.**
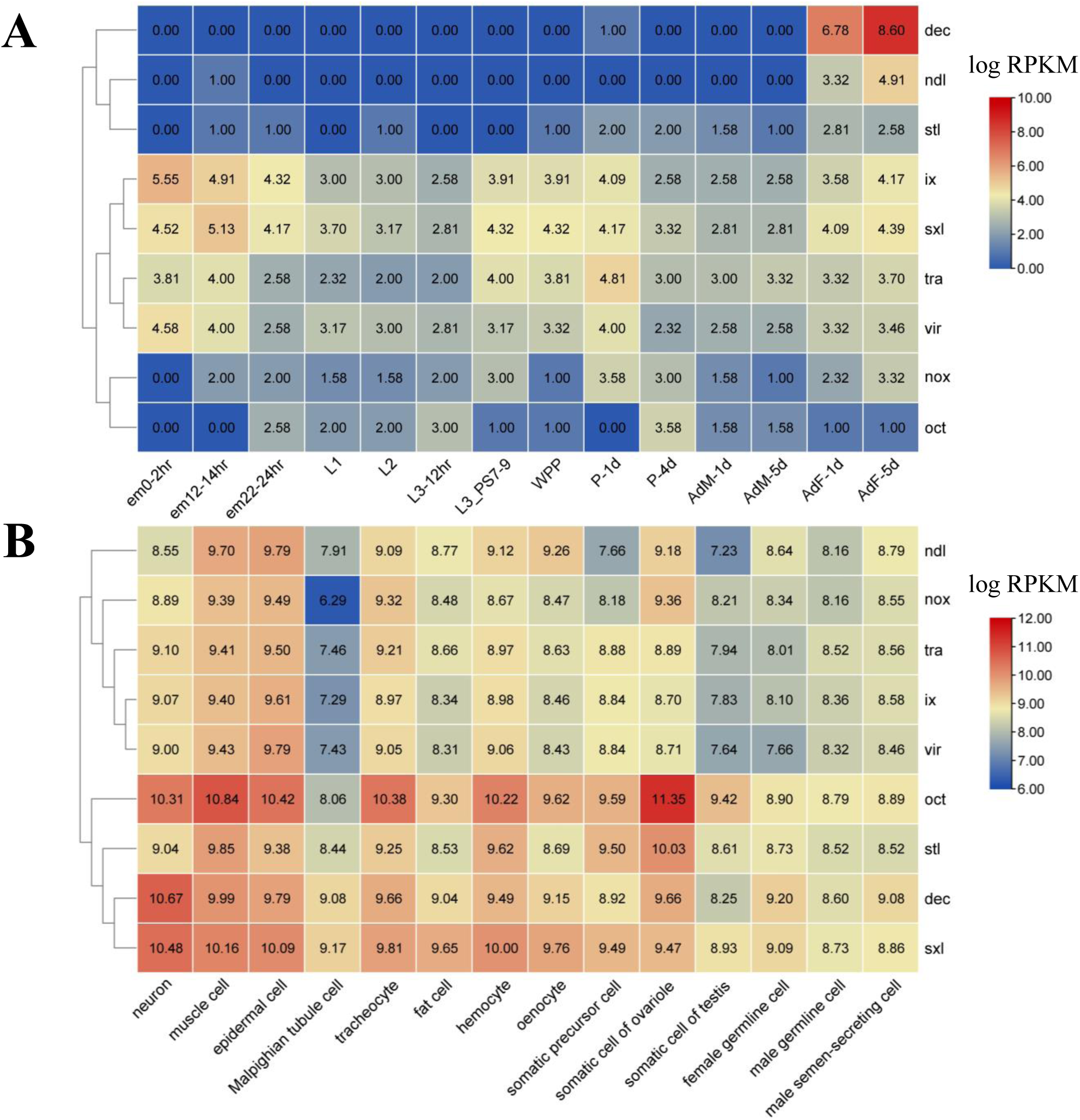
Expression profile of target genes. (**A**) Expression pattern in different developmental stages. em - embryo, L - larva, WPP - white prepupa, P - pupa, AdM - adult male, AdF - adult female. (**B**) Expression pattern in different adult cell types. The values inside the heatmap indicate expression levels of target genes (log2 RPKM). Red color indicates high expression, and blue color represents the low expression. Genes with similar expression profiles are clustered together.

**Figure S2.**
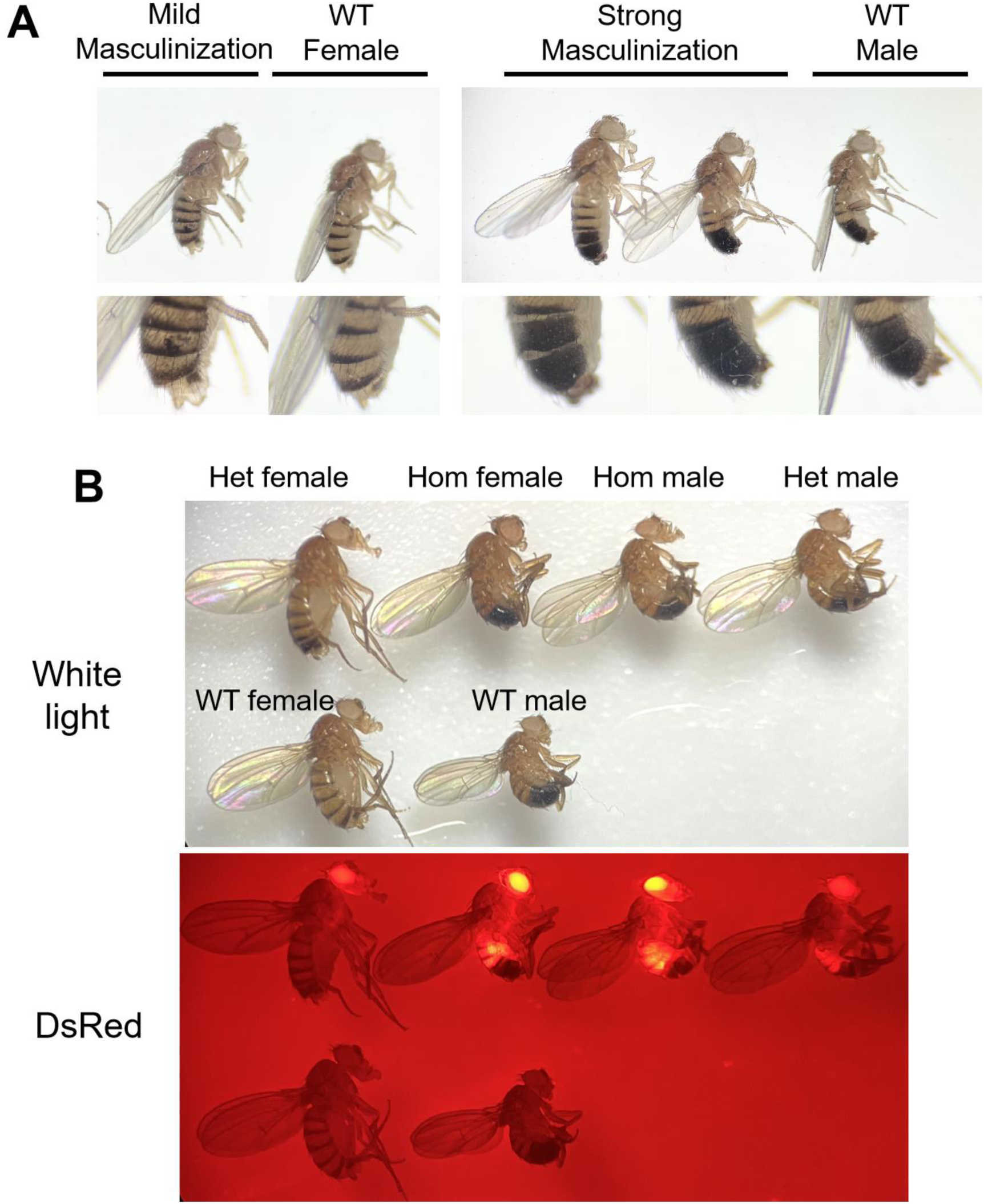
Phenotypes of drive targeting *tra*. (**A**) After crossing flies heterozygous for both *tra*-targeting drive and Cas9 alleles with *w*^1118^, two abnormal phenotypes were observed in their offspring. These were genetically female but exhibited various levels of masculinization. We defined unusual patchy pigmentation in dorsal fragments as mild masculinization, while male-like flies were defined as strong masculinization. (**B**) After intercrossing drive heterozygotes (Het) in the absence of Cas9, homozygous (Hom) females showed male phenotype, while heterozygotes were identical to *w*^1118^ flies (wild-type, WT). Homozygosity is indicated by strong fluorescence and confirmed by genotyping.

**Figure S3.**
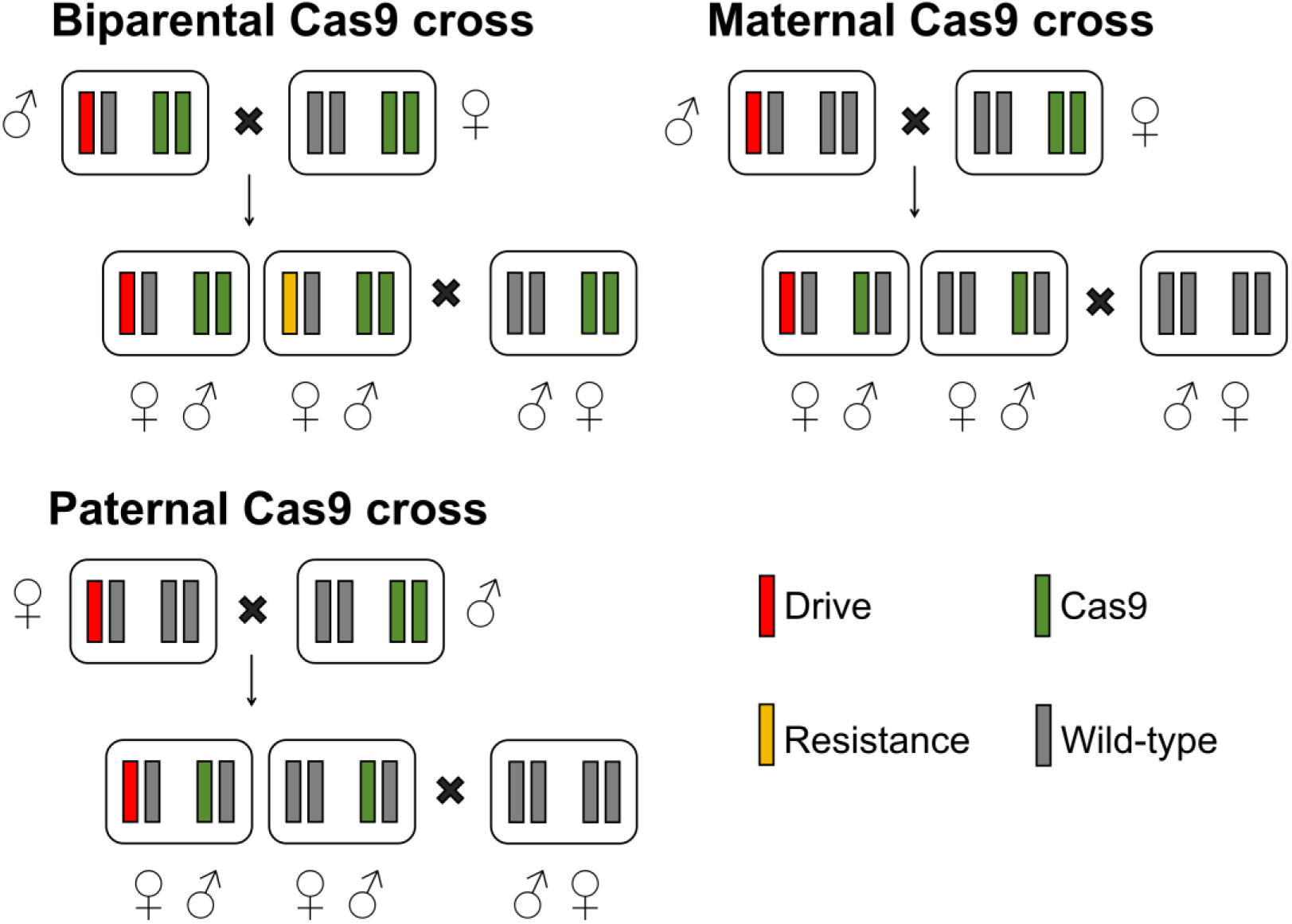
Illustration of cross schemes for fecundity and fertility tests. The biparental Cas9 cross was specifically designed for assessing drive fitness in cage populations. In the initial generation, males heterozygous for drive and homozygous for Cas9 were crossed to Cas9 homozygous females to generate drive and non-drive flies for fecundity and fertility assessment. In this case, the fitness of drive females was likely reduced by maternally deposited Cas9 and zygotically expressed gRNA. In maternal and paternal Cas9 crosses, Cas9 was provided by female and male flies, respectively. Biparental and maternal crosses have identical maternal effects in their female offspring, which are not present in the parental cross. Fitness costs from somatic Cas9 expression would still be present in all crosses, and these are likely higher in the biparental cross.

**Figure S4.**
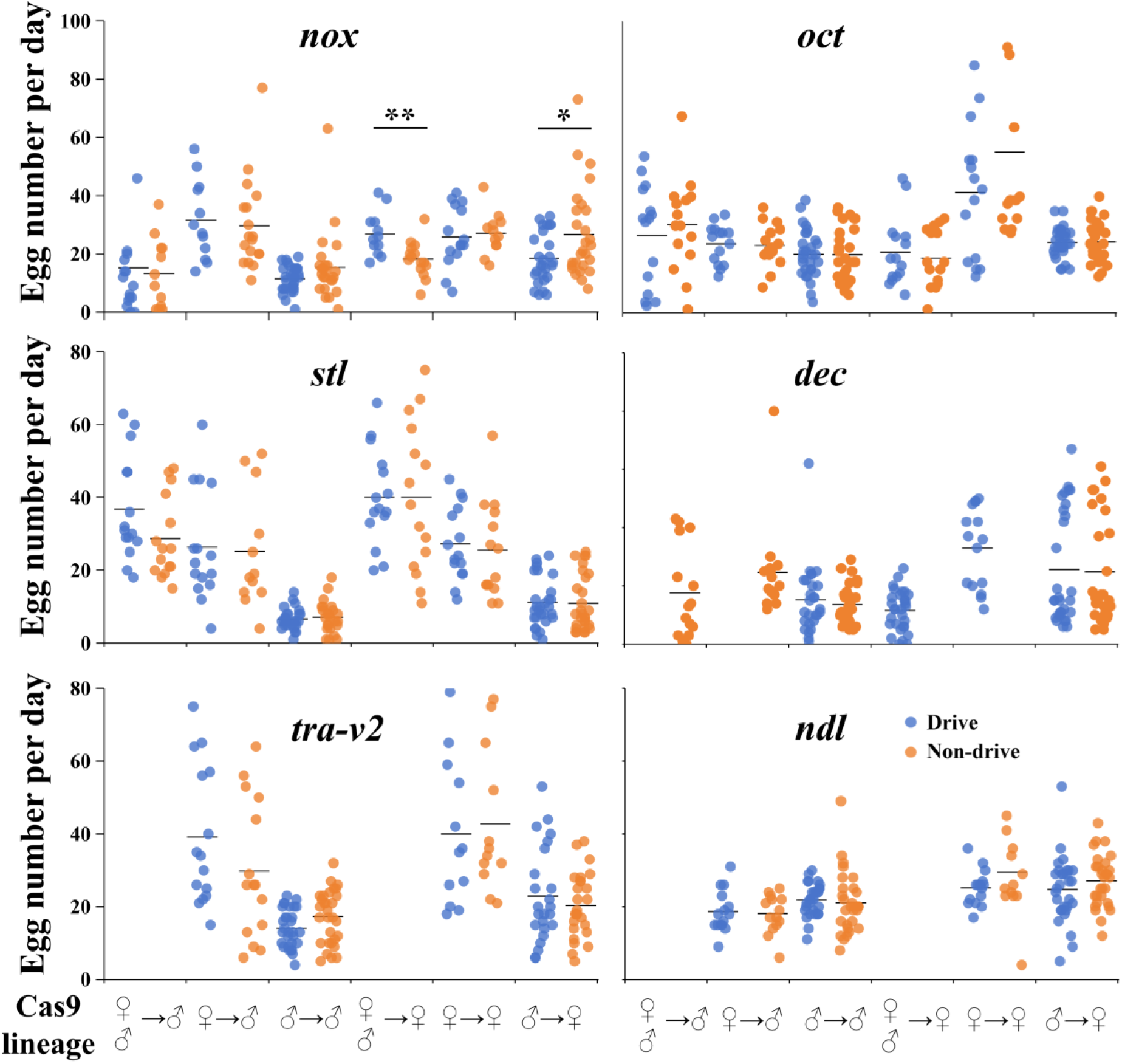
Female fecundity. Thre<u>e</u> cross schemes were applied, as shown by Cas9 lineage. For biparental Cas9, crosses were made between male drive heterozygotes and wild-type females at the drive site, and both parents were homozygous for Cas9. For maternal or paternal Cas9 crosses, either drive heterozygote males or females (without Cas9) were crossed to opposite sex flies that were homozygous for Cas9. For all three crosses, drive and non-drive progeny were then crossed to non-drive flies, and the offspring were phenotyped. Drive and non-drive flies are marked with dots in different colors. Biparental Cas9 cross data was not collected for *tra-v2* and *ndl*, and only partially collected for X-linked *dec*. Drive and non-drive flies are marked with dots in different colors. Significant difference is indicated with \**p*<0.05, \*\**p*<0.01 (*t* test). Raw data is provided in Data Set S4, S5 and S6.

**Table S1.**
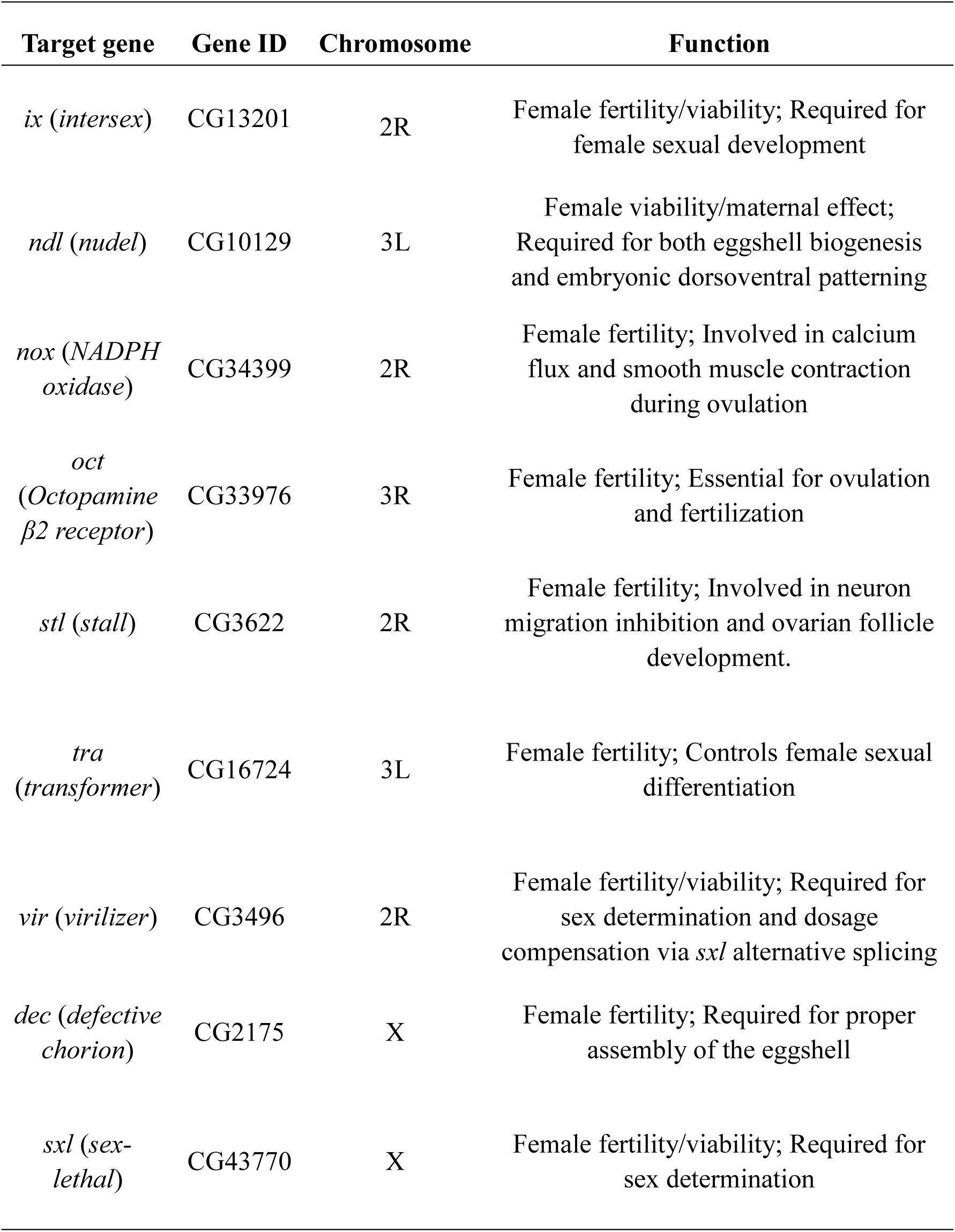
Target gene information.

| **Target gene** | **Gene ID** | **Chromosome** | **Function** |
| --- | --- | --- | --- |
| *ix (intersex)* | CG13201 | 2R | Female fertility/viability; Required for female sexual development |
| *ndl (nudel)* | CG10129 | 3L | Female viability/maternal effect; Required for both eggshell biogenesis and embryonic dorsoventral patterning |
| *nox (NADPH oxidase)* | CG34399 | 2R | Female fertility; Involved in calcium flux and smooth muscle contraction during ovulation |
| *oct (Octopamine β2 receptor)* | CG33976 | 3R | Female fertility; Essential for ovulation and fertilization |
| *stl (stall)* | CG3622 | 2R | Female fertility; Involved in neuron migration inhibition and ovarian follicle development. |
| *tra (transformer)* | CG16724 | 3L | Female fertility; Controls female sexual differentiation |
| *vir (virilizer)* | CG3496 | 2R | Female fertility/viability; Required for sex determination and dosage compensation via *sxl* alternative splicing |
| *dec (defective chorion)* | CG2175 | X | Female fertility; Required for proper assembly of the eggshell |
| *sxl (sex-lethal)* | CG43770 | X | Female fertility/viability; Required for sex determination |

**Table S2.** Homologs of target genes across different insect species.

| Species name | Protein access number of target gene |  |  |  |  |  |  |  |
| --- | --- | --- | --- | --- | --- | --- | --- | --- |
|  | *dec* | *ix* | *nox* | *oct* | *stl* | *sxl* | *tra* | *vir* |
| *Drosophila melanogaster* | NP<br>72720<br>2.1 | NP_61<br>0677.1 | AEO203<br>23.1 | NP_0011<br>63596.2 | NP_0010<br>97423.1 | NP_0012<br>59303.1 | NP<br>524114.1 | NP_5249<br>00.1 |
| *Ceratitis capitata* | XP<br>02071<br>5373.1 | XP_00<br>45375<br>14.1 | XP_0207<br>17669.1 | XP_0207<br>13689.1 | XP_0121<br>54838.1 | XP_0121<br>58509.1 | XP<br>0207149<br>97.1 | XP_0121<br>57537.1 |
| *Spodoptera frugiperda* | - | XP_03<br>54531<br>21.1 | XP_0354<br>54891.2 | XP_0354<br>41849.1 | XP_0505<br>51061.1 | XP_0354<br>48223.1 | - | XP_0505<br>49381.1 |
| *Plutella xylostella* | - | XP_03<br>79625<br>58.1 | XP_0484<br>87167.1 | XP_0379<br>67636.1 | XP_0484<br>83858.1 | XP_0484<br>82129.1 | - | WQH200<br>77.1 |
| *Aedes aegypti* | - | XP_00<br>16543<br>64.1 | XP_0016<br>60738.2 | XP_0216<br>93342.1 | XP_0016<br>60627.2 | XP_0217<br>04539.1 | - | XP_0016<br>60468.2 |
| *Anopheles gambiae* | - | XP_32<br>1918.4 | XP_0615<br>17613.1 | XP_0034<br>36280.1 | XP_0615<br>05633.1 | XP_0615<br>05793.1 | - | XP_0615<br>14589.1 |
| *Culex quinquefasciatus* | - | XP_00<br>18496<br>07.1 | XP_0381<br>11558.1 | XP_0381<br>15039.1 | XP_0381<br>20942.1 | XP_0381<br>20092.1 | - | XP_0381<br>11858.1 |
| *Leptinotarsa decemlineata* | - | XP_02<br>30284<br>83.1 | XP_0740<br>37292.1 | XP_0740<br>32976.1 | XP_0740<br>32509.1 | XP_0230<br>12638.1 | - | XP_0230<br>12377.2 |
| *Rhynchophorus ferrugineus* | - | KAF7<br>27197<br>5.1 | KAF726<br>7617.1 | KAF727<br>4530.1 | KAF728<br>0146.1 | KAF728<br>6598.1 | - | KAF727<br>3791.1 |
| *Periplaneta americana* | - | XP_06<br>96767<br>39.1 | XP_0696<br>81706.1 | XP_0696<br>95476.1 | XP_0696<br>81719.1 | XP_0696<br>77446.1 | KAJ4447<br>572.1 | XP_0696<br>88161.1 |
| *Blattella germanica* | - | PSN42<br>758.1 | PSN4709<br>0.1 | PSN5195<br>7.1 | PSN5391<br>0.1 | PSN3034<br>3.1 | QGB210<br>93.1 | PSN3613<br>9.1 |

“-” indicates no hit.

**Table S3.** Maximum-likelihood analysis of fitness in cage populations.

| **Cage** | **% Effective population size** |  |  | **Female heterozygote fitness** |  |  |
| --- | --- | --- | --- | --- | --- | --- |
| *nox/nanos* | 0.030 | [0.005 | 0.084] | 0.262 | [0.000 | 0.664] |
| *oct/nanos* | 0.025 | [0.007 | 0.057] | 0.245 | [0.000 | 0.664] |
| *oct/CG4415* | 0.046 | [0.021 | 0.085] | 0.318 | [0.166 | 0.508] |
| *stl/nanos-cage1* | 0.160 | [0.032 | 0.421] | 0.681 | [0.426 | 1.065] |
| *stl/nanos-cage2* | 0.261 | [0.072 | 0.638] | 0.205 | [0.098 | 0.327] |

Brackets indicate 95% confidence interval.

## References

(1) Bier, E. Gene Drives Gaining Speed. Nat Rev Genet 2022, 23 (1), 5–22. 10.1038/s41576-021-00386-0.

(2) Nolan, T. Control of Malaria-Transmitting Mosquitoes Using Gene Drives. Philos Trans R Soc Lond B Biol Sci 2021, 376 (1818), 20190803. 10.1098/rstb.2019.0803.

(3) Kefi, M.; Cardoso-Jaime, V.; Saab, S. A.; Dimopoulos, G. Curing Mosquitoes with Genetic Approaches for Malaria Control. Trends in Parasitology 2024, 40 (6), 487–499. 10.1016/j.pt.2024.04.010.

(4) Champer, J.; Buchman, A.; Akbari, O. S. Cheating Evolution: Engineering Gene Drives to Manipulate the Fate of Wild Populations. Nat Rev Genet 2016, 17 (3), 146–159. 10.1038/nrg.2015.34.

(5) Wang, G.-H.; Hoffmann, A.; Champer, J. Gene Drive and Symbiont Technologies for Control of Mosquito-Borne Diseases. Annu Rev Entomol 2024. 10.1146/annurev-ento-012424-011039.

(6) Green, E. I.; Jaouen, E.; Klug, D.; Proveti Olmo, R.; Gautier, A.; Blandin, S.; Marois, E. A Population Modification Gene Drive Targeting Both Saglin and Lipophorin Impairs Plasmodium Transmission in Anopheles Mosquitoes. eLife 2023, 12, e93142. 10.7554/eLife.93142.

(7) Gantz, V. M.; Jasinskiene, N.; Tatarenkova, O.; Fazekas, A.; Macias, V. M.; Bier, E.; James, A. A. Highly Efficient Cas9-Mediated Gene Drive for Population Modification of the Malaria Vector Mosquito *Anopheles Stephensi*. Proc Natl Acad Sci U S A 2015, 112 (49), E6736–E6743. 10.1073/pnas.1521077112.

(8) Kaduskar, B.; Kushwah, R. B. S.; Auradkar, A.; Guichard, A.; Li, M.; Bennett, J. B.; Julio, A. H. F.; Marshall, J. M.; Montell, C.; Bier, E. Reversing Insecticide Resistance with Allelic-Drive in Drosophila Melanogaster. Nat Commun 2022, 13 (1), 291. 10.1038/s41467-021-27654-1.

(9) Yadav, A. K.; Butler, C.; Yamamoto, A.; Patil, A. A.; Lloyd, A. L.; Scott, M. J. CRISPR/Cas9-Based Split Homing Gene Drive Targeting *Doublesex* for Population Suppression of the Global Fruit Pest *Drosophila Suzukii*. Proc Natl Acad Sci U S A 2023, 120 (25), e2301525120. 10.1073/pnas.2301525120.

(10) Meccariello, A.; Hou, S.; Davydova, S.; Fawcett, J. D.; Siddall, A.; Leftwich, P. T.; Krsticevic, F.; Papathanos, P. A.; Windbichler, N. Gene Drive and Genetic Sex Conversion in the Global Agricultural Pest *Ceratitis Capitata*. Nat Commun 2024, 15 (1), 372. 10.1038/s41467-023-44399-1.

(11) Ma, S.; Ni, X.; Chen, S.; Qiao, X.; Xu, X.; Chen, W.; Champer, J.; Huang, J. A Small-Molecule Approach to Restore Female Sterility Phenotype Targeted by a Homing Suppression Gene Drive in the Fruit Pest Drosophila Suzukii. PLoS Genet 2024, 20 (4), e1011226. 10.1371/journal.pgen.1011226.

(12) Kyrou, K.; Hammond, A. M.; Galizi, R.; Kranjc, N.; Burt, A.; Beaghton, A. K.; Nolan, T.; Crisanti, A. A CRISPR-Cas9 Gene Drive Targeting *Doublesex* Causes Complete Population Suppression in Caged *Anopheles Gambiae* Mosquitoes. Nat Biotechnol 2018, 36 (11), 1062–1066. 10.1038/nbt.4245.

(13) Hancock, P. A.; North, A.; Leach, A. W.; Winskill, P.; Ghani, A. C.; Godfray, H. C. J.; Burt, A.; Mumford, J. D. The Potential of Gene Drives in Malaria Vector Species to Control Malaria in African Environments. Nat Commun 2024, 15 (1), 8976. 10.1038/s41467-024-53065-z.

(14) Deredec, A.; Godfray, H. C. J.; Burt, A. Requirements for Effective Malaria Control with Homing Endonuclease Genes. Proc Natl Acad Sci U S A 2011, 108 (43), E874–880. 10.1073/pnas.1110717108.

(15) Lester, P. J.; Bulgarella, M.; Baty, J. W.; Dearden, P. K.; Guhlin, J.; Kean, J. M. The Potential for a CRISPR Gene Drive to Eradicate or Suppress Globally Invasive Social Wasps. Sci Rep 2020, 10 (1), 12398. 10.1038/s41598-020-69259-6.

(16) Gierus, L.; Birand, A.; Bunting, M. D.; Godahewa, G. I.; Piltz, S. G.; Oh, K. P.; Piaggio, A. J.; Threadgill, D. W.; Godwin, J.; Edwards, O.; Cassey, P.; Ross, J. V.; Prowse, T. A. A.; Thomas, P. Q. Leveraging a Natural Murine Meiotic Drive to Suppress Invasive Populations. Proceedings of the National Academy of Sciences 2022, 119 (46), e2213308119. 10.1073/pnas.2213308119.

(17) Fuchs, S.; Garrood, W. T.; Beber, A.; Hammond, A.; Galizi, R.; Gribble, M.; Morselli, G.; Hui, T.-Y. J.; Willis, K.; Kranjc, N.; Burt, A.; Crisanti, A.; Nolan, T. Resistance to a CRISPR-Based Gene Drive at an Evolutionarily Conserved Site Is Revealed by Mimicking Genotype Fixation. PLoS Genet 2021, 17 (10), e1009740. 10.1371/journal.pgen.1009740.

(18) Hammond, A.; Galizi, R.; Kyrou, K.; Simoni, A.; Siniscalchi, C.; Katsanos, D.; Gribble, M.; Baker, D.; Marois, E.; Russell, S.; Burt, A.; Windbichler, N.; Crisanti, A.; Nolan, T. A CRISPR-Cas9 Gene Drive System Targeting Female Reproduction in the Malaria Mosquito Vector *Anopheles Gambiae*. Nat Biotechnol 2016, 34 (1), 78–83. 10.1038/nbt.3439.

(19) Carrami, E. M.; Eckermann, K. N.; Ahmed, H. M. M.; Sánchez C, H. M.; Dippel, S.; Marshall, J. M.; Wimmer, E. A. Consequences of Resistance Evolution in a Cas9-Based Sex Conversion-Suppression Gene Drive for Insect Pest Management. Proc Natl Acad Sci U S A 2018, 115 (24), 6189–6194. 10.1073/pnas.1713825115.

(20) Chen, W.; Guo, J.; Liu, Y.; Champer, J. Population Suppression by Release of Insects Carrying a Dominant Sterile Homing Gene Drive Targeting Doublesex in Drosophila. Nat Commun 2024, 15 (1), 8053. 10.1038/s41467-024-52473-5.

(21) Xu, X.; Chen, J.; Wang, Y.; Liu, Y.; Zhang, Y.; Yang, J.; Yang, X.; He, Z.; Champer, J. Gene Drive-Based Population Suppression in the Malaria Vector *Anopheles Stephensi*. bioRxiv 2024, 2024.05.24.595689. 10.1101/2024.05.24.595689.

(22) Beaghton, A. K.; Hammond, A.; Nolan, T.; Crisanti, A.; Burt, A. Gene Drive for Population Genetic Control: Non-Functional Resistance and Parental Effects. Proc Biol Sci 2019, 286 (1914), 20191586. 10.1098/rspb.2019.1586.

(23) Hammond, A. M.; Kyrou, K.; Bruttini, M.; North, A.; Galizi, R.; Karlsson, X.; Kranjc, N.; Carpi, F. M.; D’Aurizio, R.; Crisanti, A.; Nolan, T. The Creation and Selection of Mutations Resistant to a Gene Drive over Multiple Generations in the Malaria Mosquito. PLoS Genet 2017, 13 (10), e1007039. 10.1371/journal.pgen.1007039.

(24) Yang, E.; Metzloff, M.; Langmüller, A. M.; Xu, X.; Clark, A. G.; Messer, P. W.; Champer, J. A Homing Suppression Gene Drive with Multiplexed gRNAs Maintains High Drive Conversion Efficiency and Avoids Functional Resistance Alleles. G3 (Bethesda) 2022, 12 (6), jkac081. 10.1093/g3journal/jkac081.

(25) Champer, S. E.; Oh, S. Y.; Liu, C.; Wen, Z.; Clark, A. G.; Messer, P. W.; Champer, J. Computational and Experimental Performance of CRISPR Homing Gene Drive Strategies with Multiplexed gRNAs. Science Advances 2020, 6 (10), eaaz0525. 10.1126/sciadv.aaz0525.

(26) Prowse, T. A. A.; Cassey, P.; Ross, J. V.; Pfitzner, C.; Wittmann, T. A.; Thomas, P. Dodging Silver Bullets: Good CRISPR Gene-Drive Design Is Critical for Eradicating Exotic Vertebrates. Proc Biol Sci 2017, 284 (1860), 20170799. 10.1098/rspb.2017.0799.

(27) Khatri, B. S.; Burt, A. A Theory of Resistance to Multiplexed Gene Drive Demonstrates the Significant Role of Weakly Deleterious Natural Genetic Variation. Proc Natl Acad Sci U S A 2022, 119 (32), e2200567119. 10.1073/pnas.2200567119.

(28) Oberhofer, G.; Ivy, T.; Hay, B. A. Behavior of Homing Endonuclease Gene Drives Targeting Genes Required for Viability or Female Fertility with Multiplexed Guide RNAs. Proc Natl Acad Sci U S A 2018, 115 (40), E9343–E9352. 10.1073/pnas.1805278115.

(29) Faber, N. R.; Xu, X.; Chen, J.; Hou, S.; Du, J.; Pannebakker, B. A.; Zwaan, B. J.; van den Heuvel, J.; Champer, J. Improving the Suppressive Power of Homing Gene Drive by Co-Targeting a Distant-Site Female Fertility Gene. Nature Communications 2024, 15 (1), 9249. 10.1038/s41467-024-53631-5.

(30) Du, J.; Chen, W.; Jia, X.; Xu, X.; Yang, E.; Zhou, R.; Zhang, Y.; Metzloff, M.; Messer, P. W.; Champer, J. Germline Cas9 Promoters with Improved Performance for Homing Gene Drive. Nat Commun 2024, 15 (1), 4560. 10.1038/s41467-024-48874-1.

(31) Öztürk-Çolak, A.; Marygold, S. J.; Antonazzo, G.; Attrill, H.; Goutte-Gattat, D.; Jenkins, V. K.; Matthews, B. B.; Millburn, G.; Dos Santos, G.; Tabone, C. J.; FlyBase Consortium. FlyBase: Updates to the Drosophila Genes and Genomes Database. Genetics 2024, 227 (1), iyad211. 10.1093/genetics/iyad211.

(32) Chen, C.; Chen, H.; Zhang, Y.; Thomas, H. R.; Frank, M. H.; He, Y.; Xia, R. TBtools: An Integrative Toolkit Developed for Interactive Analyses of Big Biological Data. Mol Plant 2020, 13 (8), 1194–1202. 10.1016/j.molp.2020.06.009.

(33) Champer, J.; Chung, J.; Lee, Y. L.; Liu, C.; Yang, E.; Wen, Z.; Clark, A. G.; Messer, P. W. Molecular Safeguarding of CRISPR Gene Drive Experiments. Elife 2019, 8, e41439. 10.7554/eLife.41439.

(34) Metzloff, M.; Yang, E.; Dhole, S.; Clark, A. G.; Messer, P. W.; Champer, J. Experimental Demonstration of Tethered Gene Drive Systems for Confined Population Modification or Suppression. BMC Biology 2022, 20 (1), 119. 10.1186/s12915-022-01292-5.

(35) Champer, S. E.; Kim, I. K.; Clark, A. G.; Messer, P. W.; Champer, J. *Anopheles* Homing Suppression Drive Candidates Exhibit Unexpected Performance Differences in Simulations with Spatial Structure. Elife 2022, 11, e79121. 10.7554/eLife.79121.

(36) Hou, S.; Chen, J.; Feng, R.; Xu, X.; Liang, N.; Champer, J. A Homing Rescue Gene Drive with Multiplexed gRNAs Reaches High Frequency in Cage Populations but Generates Functional Resistance. Journal of Genetics and Genomics 2024. 10.1016/j.jgg.2024.04.001.

(37) Liu, J.; Champer, J.; Langmüller, A. M.; Liu, C.; Chung, J.; Reeves, R.; Luthra, A.; Lee, Y. L.; Vaughn, A. H.; Clark, A. G.; Messer, P. W. Maximum Likelihood Estimation of Fitness Components in Experimental Evolution. Genetics 2019, 211 (3), 1005–1017. 10.1534/genetics.118.301893.

(38) Zhang, X.; Sun, W.; Kim, I. K.; Messer, P. W.; Champer, J. Population Dynamics in Spatial Suppression Gene Drive Models and the Effect of Resistance, Density Dependence, and Life History. bioRxiv 2024, 2024.08.14.607913. 10.1101/2024.08.14.607913.

(39) Hammond, A.; Karlsson, X.; Morianou, I.; Kyrou, K.; Beaghton, A.; Gribble, M.; Kranjc, N.; Galizi, R.; Burt, A.; Crisanti, A.; Nolan, T. Regulating the Expression of Gene Drives Is Key to Increasing Their Invasive Potential and the Mitigation of Resistance. PLoS Genet 2021, 17 (1), e1009321. 10.1371/journal.pgen.1009321.

(40) Dhole, S.; Lloyd, A. L.; Gould, F. Tethered Homing Gene Drives: A New Design for Spatially Restricted Population Replacement and Suppression. Evol Appl 2019, 12 (8), 1688–1702. 10.1111/eva.12827.

(41) López Del Amo, V.; Juste, S. S.; Gantz, V. M. A Nickase Cas9 Gene-Drive System Promotes Super-Mendelian Inheritance in Drosophila. Cell Rep 2022, 39 (8), 110843. 10.1016/j.celrep.2022.110843.

(42) Sanz Juste, S.; Okamoto, E. M.; Nguyen, C.; Feng, X.; López Del Amo, V. Next-Generation CRISPR Gene-Drive Systems Using Cas12a Nuclease. Nat Commun 2023, 14 (1), 6388. 10.1038/s41467-023-42183-9.

(43) Zhu, J.; Chen, J.; Liu, Y.; Xu, X.; Champer, J. Population Suppression with Dominant Female-Lethal Alleles Is Boosted by Homing Gene Drive. BMC Biol 2024, 22 (1), 201. 10.1186/s12915-024-02004-x.

(44) Han, Y.; Champer, J. A Comparative Assessment of Self-Limiting Genetic Control Strategies for Population Suppression. bioRxiv 2024, 2024.09.23.614516. 10.1101/2024.09.23.614516.

(45) Anderson, M. A. E.; Gonzalez, E.; Ang, J. X. D.; Shackleford, L.; Nevard, K.; Verkuijl, S. A. N.; Edgington, M. P.; Harvey-Samuel, T.; Alphey, L. Closing the Gap to Effective Gene Drive in *Aedes Aegypti* by Exploiting Germline Regulatory Elements. Nat Commun 2023, 14 (1), 338. 10.1038/s41467-023-36029-7.

(46) Xu, X.; Harvey-Samuel, T.; Siddiqui, H. A.; Ang, J. X. D.; Anderson, M. E.; Reitmayer, C. M.; Lovett, E.; Leftwich, P. T.; You, M.; Alphey, L. Toward a CRISPR-Cas9-Based Gene Drive in the Diamondback Moth *Plutella Xylostella*. The CRISPR Journal 2022, 5 (2), 224–236. 10.1089/crispr.2021.0129.

(47) Champer, J.; Reeves, R.; Oh, S. Y.; Liu, C.; Liu, J.; Clark, A. G.; Messer, P. W. Novel CRISPR/Cas9 Gene Drive Constructs Reveal Insights into Mechanisms of Resistance Allele Formation and Drive Efficiency in Genetically Diverse Populations. PLOS Genetics 2017, 13 (7), e1006796. 10.1371/journal.pgen.1006796.

