## Supplementary Alignment Files for "Assessing target genes for homing suppression gene drive": dec.pdf

```
             1          10                  20            30            40
Drosophila   MRLFSLLPLLALLVVQ...........AAGQSEVTSDDPATDAGSTTNSTTDTKPRIPSQ
Ceratitis    MIWFKNVALLAFLLAISIASIASVQGSASGTAAEPQDEDTIDLTPSADPTSEKNQDNRGD

            50            60               70            80            90
Drosophila   DEILGQMPSINPIRTGNP..QMDAFYMMFPALGSLLKWGSLFPAYSILG..........A
Ceratitis    GQIVGQSLLTPLLGQGLLPAPLNMFSSMFPALNSLLLLGNLFPGASLMGPPSLLGSTQQM

             100           110           120           130
Drosophila   IPDNLQPTAAASKVVLVLADDATAKTRVARQNPPPN...................PLGQL
Ceratitis    GPSATDESPAASKVVLVLAEDRNPHVAMRVANQEAARTIRQNDMLSQMLNQFGQVDFGQM

            140           150           160                              170           180
Drosophila   MNWPALPQDFQLPSMDLGPQVGSFLAQLP..............AMPTVPGLLGAAAPVPA
Ceratitis    LSQSLEQMQLQNPGQLLQQNPGQLLQQIVPQDLSQPWSQLLSQTFQQIPGLDLSQFPLIG

                 190           200           210           220           230
Drosophila   PAPAPAAAPP.PAPAPAADPPAAPVPDAPQPAILGQAALQNAFTFFNP............
Ceratitis    QAPTSSEPTVGAAMESAGGEGAAPAADAPASAPQQAFQFPNFQTAIDAGTQMLGQALQQP

                                         240                   250           260
Drosophila   ..............ANFDASSLLG..........QSVPTFAPPNLDFVAQMQRQFFPG..
Ceratitis    NSDPTGAFQLPNIQSALDAGTQMLGQAFQQANSDSSTSGFQLPNIQSVIDSGNQMLSQSG

                          270           280           290            300           310
Drosophila   .......MTPAQPAAAGTDAQ..ASDISEVRVRPEDP.....YSQEAQMKIKSALEMEQE
Ceratitis    GLFNFRPLQSLDPATSASDAEPAASQISEVRVKPEDLPLRTTFNQQQAQKDEKMDDLKMM

                  320           330                 340           350           360
Drosophila   RQQQAQVKDQEQVPLLWFRMPTTQNQ..........DATEEKTLEDLRVEAKLRAFERQV
Ceratitis    SALQKKILHK.KLPILWFHIPTDSSERNHDFGSKTYSQTNTKSLEEMQLETKLQAFQRQV

                  370           380           390           400           410
Drosophila   IAELRMLQKIELMAKQMRSSAAAQNG....DSPYRISYPLSRTPIHKITRADIEQALRDD
Ceratitis    VTELKLLQDIERKAKELRSAAKGSMAKTFKDNNSSNESALSKIPIHKITRGDIEKALNDD

             420           430                 440           450           460
Drosophila   YVRRLVNKEAQRRARNSGIN.........TQKANALKRQAKSQDQTLSKEDIVQIMAYAY
Ceratitis    YVKKLLHKTAITTRKETSNSPFEQLLVPSSMNIKRQTMQSTMKPRQMTREDIIRLMAYAY

            470           480           490           500           510           520
Drosophila   RMANEQMESEKGKQDKVYAAYRTEQNPMMMQQRQWSEEQAKIQQNQQQIQQNPMMMQQRQ
Ceratitis    RMANANGIRMLDSSENMKTTQDSNENARQSSDNSFTQTYRQWANDARTPSQMADMRPTAK

            530           540           550           560           570           580
Drosophila   WSEEQAKIQQNQQQIQQNPMMMQQRQWSEEQAKIQQNQQQIQQNPMMMQQRQWSEEQAKI
Ceratitis    QWMDEEMQMSHKKKIDQIPTRIPMERQMEEKIERQWAEEQTQPQRFSMKQNFDTQAPHNA
```

```
              590       600       610       620       630       640
Drosophila    QQNQQQIQQNPMMVQQRQWSEEQAKIQQNQQQIQQNPMMMQQRQWSEEQAKIQHDQQMAQ
Ceratitis     QQRQWVNQKSDITALMQQHPLQQMPIMMQQPQMVGQQTPMAMQQWHMVEQTQKDQPQQMQ

              650       660       670       680       690       700
Drosophila    QMAQQGLMMTEQRQRQWSEDQAKIQQAQQMAQQTPMMMPQMQQRQWTEDPQMVQQMQQRQ
Ceratitis     TMMQQPLMVGQQTPMAMQQR....QMVEQTQKDQPQQMQTMMQQPLMVGQQTPMTMQQLQ

              710       720       730       740       750       760
Drosophila    WAEDQTRMQMAQQNPMMQQQRQMAENPQMMQQRQWSEEQTKIEQAQQMAQQNQMMMQQMQ
Ceratitis     MEHEMQQHQPQQIQTMMHQPQMVGLQTPMTMQQGQMVEQTQKDQPQQMQTMMQQPLMVGQ

              770         780       790       800       810         820
Drosophila    QRQWSEDQ.AQIQQQQRQMMQQTPMMMKERQWAEENPQSVQQQGPMMMQQQ..MPSMMQR
Ceratitis     QTPMTMQHRQMVDQFQKHQPQQIQTMMQQPQMVGQQTPMAMQQWQMVEQTQKDQPQQMQT

                 830        840       850              860         870
Drosophila    EVEDED.NKAEDDLVGEAGPQMPENEGTARHK.......VDALGVGGNK..RKKSKSKSA
Ceratitis     MVPDIERQMQEENMIGEAMPQMPENAEKARHKGWPVVDEFDIFGVGGGRDHKKKHKGGPL

                 880                                     890       900
Drosophila    PPTVINYYYAAPQR............................PVVQSYGTSYGGGGYGSN.
Ceratitis     RPTVINYYYNAGGRGPSYGGGTGYVRGGGAGYGSGSTSYTSNLNPSYGTSYGGGAGGSYS

                   910       920       930       940       950       960
Drosophila    ....AYGVPRPVNSYQSQGYRAAVGNDEVDEMLRQHQTMARTINPKQPGEVGGSESQKS.
Ceratitis     GNSAAGSYTSGYPAASSGLYRTAVGDEEIRAMLKEHSQMKMMPEPDRNPVITTDMSTTSP

                                970       980                        990
Drosophila    ...............NSNPPTTLTPAPQEQPQEHRVHKSPS.............SAPSETE
Ceratitis     FQSTTSSIDDINQQDTTSYTTSTETPLKNQKKNRTTAPSTTIEVTPSPATFKLSSNTNDN

               1000       1010      1020      1030       1040      1050
Drosophila    IENAPSSDPQVG.SIFTYGEGLLHPFMGLLPVERPD.DPWNQKPYDPHHPLYTGGGSYDA
Ceratitis     VETSSAMDPEIVGSLFSFSPNILRPFMQLQPVPLYESDPWNHKSYDPHYPLYEGGGSYEP

                 1060      1070      1080      1090      1100        1110
Drosophila    YLRDGRHRRDTHIMGQGTQHGILTPGMLERLLRIKMDFQRRFPHLYKGMLN..HHTNLTR
Ceratitis     YLQNKRMKRNSHAYTN....NILTPSILDNYLKILVLFEKSFPQLYKSLREQQQAFNLTR

                   1120      1130      1140      1150      1160      1170
Drosophila    VEVQPPVLGKISKPKTKTKPKNEDEPVFELGAAERSLFEDETNDSLEKDPEPEPDEEDDR
Ceratitis     VSVKPPVIPKVVT...........YPATELGAAELLPADIQNSTQAN.............

                   1180      1190      1200
Drosophila    DVEEPSESSEPRGFSSKKSRDENDIDYFNFDDDDVDD
Ceratitis     ...............ELKTKRDDLEDYFLFDDEEDE.
```
