## Supplementary Alignment Files for "Assessing target genes for homing suppression gene drive": ix.pdf

```
               1        10                                       20
Drosophila     .MNPNMNMMPMS..........................GP.QMMQ.......VMQSSP..
Ceratitis      .MNPNMNMHNIQQMN..................ISQVGGPNQMMQSGVSPVSMMQASPQQ
Spodoptera     MNHNQMNMHVPM..NTV...................GGTPNVGMQ.......MPVPG...
Plutella       ..MNQMNMHVPM..NAV...................AGAPNVGMQ.......MPVTG...
Aedes          .MMNQMGMHMQQ..PGV.VPGGPGGPVGMAGGPVGGVGVSPVMMQ.......SPQMQ...
Anopheles      .MMNQMGMMMQQ..QGVGVPGGPGGVGGVGMPGPGGVGVAPGMMQ.......SPQMQQAQ
Culex          .MMNPMGMHMQQ..PGV.GPGGPVGMPGGPGGPVG.VGVSPVMMQ.......SPQMQ...
Leptinotarsa   .........MLI..SLQ...................QQQQNSQQQ.......AVQQQ...
Rhynchophorus  .MFSMPDKNIQF..GMA...................KQQEMQQQQ.......AQQAQ...
Periplaneta    ...........M..NLP...................PMQQQGPGV.......MPQVP...
Blattella      ...........M..NLP...................PMQQPGPGV.......MPQVQ...

                                   30        40                  50        60
Drosophila     ..............SGPPGPVQHQQQQPPQPLQQQQ.........QAEKLDNISRVKSLL
Ceratitis      QMQGQMQ.......AQVSNAAQQQQQQQQQQQQQQQ.........QAEKMDNILKVKSLL
Spodoptera     ..............TIMQQQSPQQMQQPVMPQQPQ..........QQDKMDNISRVKTLM
Plutella       ..............PIMQQPSPQQMQQQQMQQQPMP......QQPQQDKMDNISKVKTLM
Aedes          ..............QQQQVAAQQQQQQQQQQQAQAH....QQQSQQTEKVDNISKVKGLV
Anopheles      QQQVQQQQVQQQQVQQQQVQQQQQQVQQQQQQQQQHSQSAQQQAQQTEKVDNISKVKVLV
Culex          ..............QQQ....VQQQQQHQAQQQQQQ....QQQAQQPEKVDNIAKVKALV
Leptinotarsa   ..............MQQQQ.QAQQQQQAQQQQQQQQ......Q....QPLDNISQIKALI
Rhynchophorus  ..............QQLQQ.AQQLQQQAQQQQPQQQ......QSQQIQHLDNISKIKSLV
Periplaneta    ..............QQPGPGTPQQAQQLQ.PQIQQP......Q....EKLDNISKVKSLI
Blattella      ..............QQPGPGTPQQAQQMQQPMPQQQ......Q....EKLDNISKVKSLI

                         70        80        90        100       110
Drosophila     GPLRESMFLTIRSSAFALQQNNLADN.LKRDTGAHHVPRFDKHLEDFYACCDQIEIHLKT
Ceratitis      SPLRESLFLTLRTGAFALQEHNLADN.LKRDS.VSHAGRFEKHLEDFYAYCDQIELHLKT
Spodoptera     GSLRESIPMTLKSAAQILHQNHNVDSNSQKG.IDNPVPRFDKNLEEFFSICDQMELHLRT
Plutella       GSLRESIPMTLKSAAQVLHQNYNVDSVSQKG.IENPVPRFDKNLEEFFSLCDQMELHLRT
Aedes          GPLRDSLSTTVKTAAQLLQQNNLNDAGTKGGDMSTTTPRFDKHLEEFYSICDQIELNLKT
Anopheles      GPLRDALSTTIKTAAQLIQQNTLADAGSKTVDLNN.APRFDKHLEEFYSICDQIELNLKT
Culex          GPLRDALSTTIKTSAQLLQQNNLNDAGTKGGDLNAPTPKFDKHLEEFYSICDQIELNLKT
Leptinotarsa   GPLRESLSTTIKTAAQTLNQNSQVDAGIQKG.VDVQIPRFDKNLEEFYSICDQIELHLKT
Rhynchophorus  VPLRETLALTIKTAAQTLNQNSQIDSGSQKT.PDVQIPRFDKNLEEFYSICDQIELHLKT
Periplaneta    TPLRESLSLTLKTAAQTLHQNSLVDVGSLKG.VDMAVPRFDKNMEEFYSICDQIELHLKT
Blattella      TPLRESLAVTLKNAAATLQQN...................................NLKT

               120       130       140       150       160       170
Drosophila     AMQCLQQQNSSNHYLPGPVTPMRMETFMPDN.AGPISYPTYLNTVRVHIQSAKDIHDTLI
Ceratitis      AIQCMQQLSSAQHYLPSAVTPARIEPYMQENPGGPMTYSTYLNIVRVHVQSAKDIHDTLI
Spodoptera     AITCIQQAQSAAHYLPMTVMPSRLDTGPATTDTS.LSYPQYLNTVRLQITYAKDIHDTLV
Plutella       AITCIQQAQSAAHYLPLTVIPSRLDAAPTTQETNPLSYPQYLATVRAQVSYAKAIHDILE
Aedes          AKLCMQQGASSQQYLPIPVAPTQ....PNPAETNALSYSQYLDVVKIQIGYAKDIHDTLI
Anopheles      TKLCMQQCTSSQQYLPIPVATSQ....PPLPETNALTYNQYLEVVKLQIGYAKDIHDTLI
Culex          AKLCMQQGASSQQYLPIPVAPTQ....PNPAETNALSYSQYLDVVKIQIGYAKDIHDTLI
Leptinotarsa   SLKCQSQQESSNRFLPLQVAPTRSESLG.MNDN.TQSYPQFLAAVNMQVSYTKEIHDTLV
Rhynchophorus  SIKCLMQAESSNRYLSLQVTPTRNETLG.LSES.TLTYPQFLATVGAQVSYTKEIHETLL
Periplaneta    SIECLAQGTASQRYLPLPVALTRTDPLP.SQEGAALTYPQYLATVRAQVGFAKEVHDMLV
Blattella      SIECLSQGAASQRYLPLQVALTRTDPMP.GQE..ILTYPQYLATVRAQVGFAKEVHDMLV

               180
Drosophila     SAAQNISQAD.......
Ceratitis      SAAQNISQAD.......
Spodoptera     AAAQNISPTE.......
Plutella       SAAQNISPPELPDKGQV
Aedes          CAAQNICPSE.......
Anopheles      CAAQNISPSE.......
Culex          CAAQNISPSE.......
Leptinotarsa   AAAQNISPSD.......
Rhynchophorus  AAAQNISPSE.......
Periplaneta    GAAQNVSPGE.......
Blattella      GAAQNVSPGE.......
```
