## Supplementary Alignment Files for "Assessing target genes for homing suppression gene drive": ndl.pdf

```
Drosophila      ............................................................
Ceratitis       ............................................................
Spodoptera      ............................................................
Plutella        ............................................................
Aedes           ............................................................
Anopheles       ............................................................
Culex           ............................................................
Leptinotarsa    ............................................................
Rhynchophorus   ............................................................
Periplaneta     MVTGGEDDNIPLEENSRRISDDVAESREASRRAHTSLSDWCQLRRNRRRMVTVVALVLGA
Blattella       ............................................................

Drosophila      ............................................................
Ceratitis       ............................................................
Spodoptera      ............................................................
Plutella        ............................................................
Aedes           ............................................................
Anopheles       ............................................................
Culex           ............................................................
Leptinotarsa    ............................................................
Rhynchophorus   ............................................................
Periplaneta     ATVAVLVPIIVYNSTHIKHKMELLSRYVTKSETNDGDLEVAMGSRDTDDNYSASSAFPSF
Blattella       ............................................................

Drosophila      ............................................................
Ceratitis       ............................................................
Spodoptera      ............................................................
Plutella        ............................................................
Aedes           ............................................................
Anopheles       ............................................................
Culex           ............................................................
Leptinotarsa    ............................................................
Rhynchophorus   ............................................................
Periplaneta     ATFFEGNNAILTENHFVDDNSEIKYKRTGNKITLDLKTNSLQWNEEANIDRQSREENIDD
Blattella       ............................................................

Drosophila      ............................................................
Ceratitis       ............................................................
Spodoptera      ............................................................
Plutella        ............................................................
Aedes           ............................................................
Anopheles       ............................................................
Culex           ............................................................
Leptinotarsa    ............................................................
Rhynchophorus   ............................................................
Periplaneta     SVTRIKRIRRHIEDVDRNSEDKCLGRKCRKIVQKISDKVQSLEAKVEALKTLMSELNGPD
Blattella       ............................................................

Drosophila      ............................................................
Ceratitis       ............................................................
Spodoptera      ............................................................
Plutella        ............................................................
Aedes           ............................................................
Anopheles       ............................................................
Culex           ............................................................
Leptinotarsa    ............................................................
Rhynchophorus   ............................................................
Periplaneta     ETYEEMPSQQSPEHGDTLVKNSENESSNFMLAKPVNKENKEETGDKMSKGHAVQTKFSEV
Blattella       ............................................................

Drosophila      ............................................................
Ceratitis       ............................................................
Spodoptera      ............................................................
Plutella        ............................................................
Aedes           ............................................................
Anopheles       ............................................................
Culex           ............................................................
Leptinotarsa    ............................................................
Rhynchophorus   ............................................................
Periplaneta     TNGYSNSTDTDSTLHEVTESPRSFKEIVQNESQTTISLEVTQNTDKDSVLQRDTTHRSIP
Blattella       ............................................................
```

```
                    1       10        20        30        40        50
Drosophila    ......MNYNMDEMEATRLLRHPRRWWSIGFGKRIVAISILVIIVLLFSLVYHGLVVEKI
Ceratitis     .....MAKIKLPTDDKKVFNAGCQRKCLKTTLVRVLAISGVTLMALMIILVYHAITLHEA
Spodoptera    ...............................MTSENMKERWDQLGLPPMELAE.......
Plutella      ...............................MNEKVQLEKT..LGLPPMELAE.......
Aedes         ............................................................
Anopheles     ...........MADHDQTVLDEELATINSTRMQQLVGAGRTNWLCLMMTAIGGVIVMAAI
Culex         .................MDKPQKSEKVDLLRCRKKIILKRLSLLGLVLVTVGALFWIAGL
Leptinotarsa  ......MWNDLVFREMVSPKITCFRSDGDTTPSPQPFEEPQDLVTIDLNLSKKEASFTKE
Rhynchophorus ............................................................
Periplaneta   SNLTTVANSKTSVDDRTISSASVGWLLTDQNEDKRVQAGTETGTGTDRLYEEKNAFYDDS
Blattella     .........MFCFRNVQNVEETARDILMNELISPKTAAFKSVQIDATNIFIKRCLDDILE

                  60        70        80        90       100       110
Drosophila    DQVQQIAALNARHQVLFNQPFEEDQSALIVSPQTLHFKLLDEDMNKDMEDSKNRRRKHMR
Ceratitis     GKHHPVAPTGPQYQTFVHSDNE...............KLTDERHVTGNEFDQNLIVQQLR
Spodoptera    ............................................................
Plutella      ............................................................
Aedes         ............................................................
Anopheles     AFGFYLTIP...................................................
Culex         LS.VYLTNT...................................................
Leptinotarsa  KTKCKN......................................................
Rhynchophorus ............................................................
Periplaneta   IASGLEADNLLWTTDNGKSEGGLSEGKIETTTEKPAYNSEYGINETLEKVYEGTVTEEPV
Blattella     SSSKNKLLSVN.................................................

                 120       130       140         150       160         170
Drosophila    QMLVKFRLNKKHRMRRDLHGLDLLDPVRMEA..NMQHLYTKLRSKRAREALS..QLEHEF
Ceratitis     ......SIDVRQRHGRSLP..DPLDPVFLEHGPNFDSLYAKVRNKRTRIELG..KLEKEY
Spodoptera    .................................SSGNSRDNNG...GNRCLF..PSLQKL
Plutella      .................................GTTYIKRESGSNIAQKCFH..PWLKRA
Aedes         ............................................................
Anopheles     ..................ASALSQQPSNLIELEPIEVQSPEELETIRETYFK..AEAHDI
Culex         ..................FSAP..KPVSLEGNSFIFTYRSEIHSTSSNSISD..TVLEPI
Leptinotarsa  ............RVYKILLYMFLVNIIVLVIFGILEYINEETYLFKTKPPTPHTAVKTEN
Rhynchophorus ............................................................
Periplaneta   TVNILIAQNLETQEKNEENALSTQTTSIFANYVNIDNRYEDITRKTSEELYDKDQSLDIV
Blattella     .............RTLEPQNGVLRRLTPANTTELVPETTTVHQPNETEQPVG..EAVPII

                       180       190       200       210            220
Drosophila    VRCKKHTP..QDCMSAFLRMYKMAKEVTEKMEKMKAIMREQQPKLE....SSSMESHEQK
Ceratitis     LRCKKESVNSSSCMVAFLKLYDLAKEISEKVDKMKEIFDDNEETLQRSNSSESTESHEVN
Spodoptera    ILLLLLTFFAVGTYILLLRISENTD....STKIIIVTGNNDLPNVTLRNMELYSIHLDKE
Plutella      LQTVLLTACAVGTFVVLLRLLQSTDNNNKSTEIIVVTGLRDSPNITVQ...QHNFEIILQ
Aedes         ............................................................
Anopheles     LMGSLELLELLNDLDPLLQRHRSRREAFGAEPVPNRISRGDDRSGEITYGVRKRVQREFD
Culex         RNLLSSVFNLLGIQAEHNTRNKRDIEQDNRISLVDHNQLDKEEDQSNEHGFVAVSPEVVT
Leptinotarsa  IANELIIDDSKDPLEYLVLKCLETDTVENCCSFFSNAETNDDFLELCKDEQESSPNNFID
Rhynchophorus ............................................................
Periplaneta   TRNNGIHSTDRVILTSSHQFQEEENDSTTTSELNTHSFTSKDDGLVVTSIEIYESASTPK
Blattella     EGSVIITRVKENDRDEKNANNNTVTTQRENVYHKNISVTQDRYSVNVTSAMSIVESSLKP

                     230       240       250                 260       270
Drosophila    G...TFSPADLIQVTTAEATTVAVHATEKPARTKIK..........PSRISWIIDGHDHD
Ceratitis     RRNETDTTKDGSNDTTTAATTEARLITVTEAASVTNRNASANGEEEPIHYNWIIDGQDED
Spodoptera    AKNETSEESSLDPLIIERKKRDTLEKTAENMEEPIL........................
Plutella      LRNKTSNRS........RRKRSITSK..QDLLATFK........................
Aedes         ............................................................
Anopheles     LGTVPIDYRGILVGSESPRAPRKRRHTDISLDELER........................
Culex         QSKRTLRILYAPLENEQTQHNDIIRSLKRRKRRSLR........................
Leptinotarsa  IRRRRGIETKISRRQVNHINGDVIQEVYPDYYYDYY........................
Rhynchophorus ............................................................
Periplaneta   TSQDWISYSDKQRSFPTASSYELDYKNTKITAGTIEDKYETSIRTTKREITKSERQEFGN
Blattella     ITEKPVDKINLFLNNTTNKNLDEQNSLLDSSPVISN........................

                     280                                     290       300
Drosophila    ESPVYTDGAPKKE..............................TTKAPWNTTQLVEITTT
Ceratitis     ....ITDQAPLKENVTEVSDFVSTTTMAAIKKVTNITENLPLRTSRGPTSTSTEATPTES
Spodoptera    ...........................................YSDELLQKSRELMMEYD
Plutella      ...........................................LHEEVVKKARNIVMNHD
Aedes         ............................................................
Anopheles     ...........................................AKRSAQKDYQRLEKEYN
Culex         ...........................................DKRYAS.AIKELENEYS
Leptinotarsa  ...........................................YNNGHYKSENEVTNYGE
Rhynchophorus ............................................................
Periplaneta   VGTENSVGTTLNEITVIGDHELHNRETERFIDTTMLTEGEDNELNYIETLRIEDIVEQGP
Blattella     .........................................TFKETVETNGKSNVMNDQS
```

```
                   310       320       330            340       350
Drosophila    KIDATATERTTVESTTEKISWILDHFDKPQEILRT.....TEGPGQRIIRNVTTTSASSE
Ceratitis     GKNHNDTSFTHTENESHKISWILDGYENDNETMTVDKPIINETTAKPTHMNVATENSANE
Spodoptera    AHCKNENKNEICKDLVTKLRELQNDDEMKI..............................
Plutella      KPCNSDTQDSICRDLVHKLDLVTK....................................
Aedes         ............................................................
Anopheles     RCRKEAPNGKLCDQIYEKLQRLSEEVNARFLEMANLLQGFSEMG............PAKE
Culex         RCKKEIPKERDCEKIYEKLKKLSEGINKNFIAMSNLIHDFQDKQSADLAKELSDRFPKKD
Leptinotarsa  TYYDHQQENSRFSENQGDYKFSKENFHTPAETQFLPPHYINTYESMIPDIQENIQAYDNP
Rhynchophorus ............................................................
Periplaneta   NIKNTGNELNYTETTNTEDTALQEGTENEGNKQNYMETTDSEETEIEKESENEDKLSKYT
Blattella     STDNSQNKTVVQEVEEEKKPSITLHQTESTVHFKTNAYSINNQPQWGLKDTPSSTFGFRP

               360       370              380       390       400
Drosophila    PIVDTENTNSDHVPTTENGLVFN........ITTDGPVETTKSTAQRKLSFDWILDGEEN
Ceratitis     TTTPSTIKVTVHEKPEKISWIIDGHHDDDKEKAEEDEEEFVANTTKSNLSKEIVSIGTEK
Spodoptera    ............................................................
Plutella      ............................................................
Aedes         ............................................................
Anopheles     KTTPQYEEMYYSSTERST............SYR..PEMHAPKNDP...............
Culex         HKKYKEDQDSDENGERKT............MKKDKPDVAQPSNWPTVHAMPSNMQLVTEI
Leptinotarsa  SALININSDNTNKYETNFSVHYR.....................................
Rhynchophorus ............................................................
Periplaneta   ETTNFVETGIKKGPENVDNESDYIKTTKTDHAEIEKGYEIVDNSSEYTETTKPEDKEIKK
Blattella     PILQTDEPWHPIVPNYRT..........................................

            410       420       430       440       450       460
Drosophila    VEPEVKSTNTTTTTAATTTTGATSETIIVTTELPKITFDWIIDGREVVEPQETTTEVTGT
Ceratitis     VLWILDENDENDENVETTAKETPKSTTQIKDKSDRIS..WIIDGHDDESVTSSTTRLTTS
Spodoptera    ...........KEDQPKMNLDVNNNVQRVATEGKAKNSVDFNDLTKRDTLMNLEHFMDNV
Plutella      ...........KRSAGHTNEGLKN.HEDVLTSARESDTIIEDVNHEGDHIMDANEGEDHH
Aedes         ............................................................
Anopheles     ............KITTVEQLVNQMNITQAIPKEKLRTSAEDEMVTTTPMLPVFDPSLHKP
Culex         MGTTTSLPSSQQKGNSTLQVNNATHRADVQTLDDLIQSIKSTFMNDSKQSEHFNPAYHNP
Leptinotarsa  ..QIPEFIAAAEKSTTNPNLADQYTQTTTENAPLRVTLRLEYPFDDNISDFEATTATTEI
Rhynchophorus ............................................................
Periplaneta   LSEIVNNESETTNSEGTENQNGSESEDKESENTETTISEETDIQKGSENEDNNQENIETT
Blattella     ...........QTTKPVHSYEYDDNIGTGLAEVMVVPPSALENSNSEEFNKKYSSRLGQP

            470       480                   490       500       510
Drosophila    TERLRKMPFDWIIDGEEVVEP............QENVTTTTIATTVAVSTTEINERIHNS
Ceratitis     STRNIFEELEEVTEGVEKEEEKLRKLKYDAMIEQEQVESKTPRTTVAEQPTTTAPLKHAT
Spodoptera    PMPMNARNHDPLPS..............................................
Plutella      DHGFQARSQYPGPAY.............................................
Aedes         ............................................................
Anopheles     IENSSIRSLDDLFDHMRLSGED.......ASTTEWTTERRELFEFEIDSTSTSPVTVKPS
Culex         VTVDCVKNLDQLLETLEEFKTGNVTNKNCTKTNHSSTPAPGDFRFAKPSHSTNSPYKAED
Leptinotarsa  IHEEVTTEIFSISSEAFTQSTRSNTEETETIFDGNIDVATYQISTISSAYNDGQSIEEKS
Rhynchophorus ............................................................
Periplaneta   NSEETEIQKESQNEDSEPESTETTNYENTRIQIETESENNGLKYTEATNSEITVAEEGIE
Blattella     AAEGSDKKNQVMSVGTIGDFGVN.....................................

               520       530       540       550       560       570
Drosophila    TAYPTKPKPVKFDWIIDGGESSGEVSTSSTSQPKLTTREAISNPESPRSSHPLDNPTSIE
Ceratitis     SSLVMNQTVHLTNTAGITITYGGDCMTTTTTKPEQLEQEKARASDTQDRSHPLDHPSSAE
Spodoptera    ........................AHMVPHSHSVTPQHLSDTCLLARLIKQNYPRAKEIY
Plutella      ........................PNPGPSYPIVAPAPIADTCLLARLLKHNPQQLHGVY
Aedes         ............................................................
Anopheles     TQP..................TSKRRTTTKSPLQAARNLHDVVHQLQLAQMLQP.HPFHE
Culex         HRPHVDECIQTLDDLMDTALFKKSRKCSPKAPSSPQRNIPDTTIDTLHSQFGQPAHPIHE
Leptinotarsa  ESDETITHLEPMSLDSYDAHITNLTRIASETVNSKPPETFEGSIANYNGNSELTSERSNE
Rhynchophorus ............................MRNLSCGSGLEPSSKFRHHLTHAGLYQNVP..
Periplaneta   YEANELANTAATNSIGTKVYVGNDEDNEISDTEIAHSENTGTQEKTKDEDNGLDYTETAT
Blattella     .....................DTVSRVSLHILGTGSASNETSVLHTTSGNGIIVRKPPNT

               580          590          600         610       620
Drosophila    NMLESFEQHEEQK.......PILRVLNANESS..SETVTDGYERQLWLKKFED..QARPN
Ceratitis     NMLEVSEKYHKFK.......PFKAVSVSPKLKNITSTTTSPYNAEMWEKLFRQSSAVQHQ
Spodoptera    EPSSDYLHTQHHE.......HPVMSKAR....FFSPSVHTPYIQERDMDMQRRK...IHP
Plutella      DVPPSYIPHSPVP.......VPYQQIPPPYPEYGPPAYHVPYHGRAVSDVQEHEDHNVHP
Aedes         ............................................................
Anopheles     DLVMS..PPENIN.......EFVMSRSRNREEMHHDTTPSPMRAVVHQQHQHLQHQTQPQ
Culex         ELGDHSTEPQDIH.......DFIMARSRNRYQYNHD........YVDHRFPTTIEAPDSR
Leptinotarsa  NYLNFSTDGVSIT.......TKSSLLDSTSTRYEQDNIASDFKKTQHTQGRQRFTAEEND
Rhynchophorus ........RGTLR.......PIAMDYTLPN...........CLGRSSTSTAS........
Periplaneta   TVDTEFQVQGATANEDNREITFSETVENYNVKNTTDYFKLTENKNDKTKKINSRDTSTVS
Blattella     IPLSYISTQQPPR.......TIANPFTLPELPILHQFRDVDAILKSPTPTASESQTESYH
```

```
              630       640       650       660       670       680
Drosophila    QNELIDTFGTALDAKALDKMGPKINPLNGHTWNAADAQILSLCERVALRMRNKVATMSDG
Ceratitis     QDELIDTFG.PTDIQSMLEFGAKLNTANTNTQGLLDSQFLSLCDQIVARR.....LLNKG
Spodoptera    QD..........................................................
Plutella      QD..........................................................
Aedes         ............................................................
Anopheles     SPHQPLVVG.........................PSAPFLNLCDQLRS...........A
Culex         TTKQMATLG.........................ASGPFLELCNQFRNQPNQQGPLKTVS
Leptinotarsa  TPNKVKSVQ...................................................
Rhynchophorus ............................................................
Periplaneta   KPFQFQVYEYQSTVDDGNIPVTTMDIEKYSENELMDKSGTDISMGYFTSSGASSNFLSFA
Blattella     ADFKWIQNSTNQDLDAIPIITVKNISTYNYQSESIRNDYKQGTSTNNSIKEISKFSPNMG

              690       700                           710       720       730
Drosophila    ETKEKGETFTASPSVQFTSRAPG...............GFPVSGETMKASAQFMFNPNFG
Ceratitis     MNTEE.EPFTPAPNPQFTSRGPG...............GFPVTGETMKATAQFIYNPNFG
Spodoptera    ............................................................
Plutella      ............................................................
Aedes         ............................................................
Anopheles     AGNGHLKPTHPGATFQHSP...................GIPITGEATKASSQIIVNSAFG
Culex         SNHHNFQAQQFGPGGFGQP...................NIPITGETLKASSHVSFNSAYG
Leptinotarsa  ........................................QSSPPELSNNQQTLYNPCYY
Rhynchophorus ..............................................IIHPHHSLWQER..
Periplaneta   TTMSDILDERHISDVHPAGSKDPMAPLEDTALNAEHILPTTTRNNKGSNTSSVVNRPLH
Blattella     IGLALPADDDDDTLFKDTPSKPG...............IVLDSPSIGSHQNQDLWNEMKN

                     740               750       760        770       780
Drosophila    MPSIPVCFYMTPANFRMP....MWSNTPTFMGMQGAHFGGS.SNPGAGIFFVPQQFGPSG
Ceratitis     MPTIPICFYITPANFRMANQSPMWSNN..FAGMSGGFSNPMGLNTPNGVFFVPQNFGPTG
Spodoptera    ............................................................
Plutella      ............................................................
Aedes         ............................................................
Anopheles     AGYVPNTVCFYQNAPPVGAQPTAYGFRPAGSGAPYVYPNAPGYPPYQPHQQQPYHQQHHH
Culex         G.FVPNHFCFYQMPQQYGAVRPGYGG..................................
Leptinotarsa  PYANPNPYSQLYRVPDISQLMRIVPADTPKSPQFVLN....PPMYPYTQPSLVPPVQFS
Rhynchophorus .........HHLRTPMSCIG........................................
Periplaneta   FPPLTSFNAFSTDNFAHTQFGNTAVDEVMKSHLQVEAEPAVLVPPYWVPYPMCVYKVPPS
Blattella     EDNIDNLKKSNSRNPNPGLGSVSPIEESHWVELVNVTTSAVRPQKFKTHEEIPSRQPQRP

              790          800       810         820       830       840
Drosophila    NFFGGSGGSGAGGQ...GANIFSKNASPQKPTNGQQQ..VYCSYMQNQSGQGAGQSQTSS
Ceratitis     NFFGPSGSTGTGGSQTSMGNVFGKNAAARK...QQQQ..FFCTYMQNNGNNSPSGGSGRG
Spodoptera    .........................................VDIFMSMLTPKAATPRES.
Plutella      .........................................EIILRSHTTTPRITLTN..
Aedes         ............................................MMLQSSQELFVG....
Anopheles     HHPGAGAMINMPVS...IQNLHQAGARTSELPADQNGPILLCSMMQNVEQPTRHHAGNET
Culex         VQTNPGGIINVPVN...IHNIPGQRSAGEIPPGSQ....LICNIVPAKATTESYQSRAIS
Leptinotarsa  TSQWPAQVQQAQYMPNFWPQQAQQGQSGQMKVSGPGGQYYVCNPISPVTNGVVSMSGVEI
Rhynchophorus ..............VDRKPPSSRGRICSIVLAIASFVVLVIVLAIAGLALYMGISNTEGP
Periplaneta   RLLFTSGAGPLESEGSLTAVSQNQDQYQWYFPTGPGEQFLYPGVIKTHYPPQLGSSPQQY
Blattella     NWGDQAYRSRPNYRIQEIESSTPRTQMAENTSPTPLITLIPVRSNSGVGRPLRPRPQLPS

                        850       860               870       880       890
Drosophila    QQQQGGQS..AFSNANFKMRHANQTNT......ANQQGQIIYASYAGLPQQP.IQERSRC
Ceratitis     GLGGSGINGMGFRTDSFKMRTSGQ..........NLTNDIIYASYTDIPDHLGHADHFQC
Spodoptera    ..................................................NQTKASREAM
Plutella      ..................................................ESLSSTREAD
Aedes         .................................................STRARG.HHRH
Anopheles     HPEDAEEELDENLDTMFAAG............................RSLAARAKGRHH
Culex         QPTPNDDGIEPRITSGNVSAGTEPT....................PHQDSLRAHGNRRRH
Leptinotarsa  RRQASNLQDMLQNVTINAVAEELS..........................RNKTREASII
Rhynchophorus ANAIISFGCSAKVIKGDR............................FVGNLEEKAQKYKR
Periplaneta   VYCPPVIPSLRLQSQQFASMTGSTGFQRDSTNQDFVRGTDEIDDPLVQKLLGVQPKAAEG
Blattella     HRIQIGINSTSEYFFPRSKPDIKDNN................DMKILNIQSEHRSDSYKP

                  900       910       920       930       940       950
Drosophila    PEPDQFSCFGQQECIPAARWCDNVVDCSDGSDESACTCADRVDEERLCDGYEDCPMGEDE
Ceratitis     SLPEQVACYGGKECIQESAWCDGFVDCADGSDEAACRCIDRLASKRICDGYADCPMGEDE
Spodoptera    CPRGTVSCLSGDTCIDEKQWCDGNVDCLDVSDEASCACKDRVDKSRLCDGYFDCPFGEDE
Plutella      CPIGSVSCEDGGSCVAQKQWCDGLVDCADASDEARCTCKARVDKSRLCDGYYDCPFGEDE
Aedes         CPRGKVPCADGIQCVLSSHLCDSRVDCFDGSDESHCSCLSRLAGKRRCDGYVDCPLAEDE
Anopheles     CRRGSVPCFSSHQCVKRSSWCDSKTDCMDGSDESACSCVSRLPKRKLCDGYADCPLGMDE
Culex         CVRGWIPCANGQQCIRNTEWCDSKTDCLDGSDESHCSCLSRISEKRLCDGYADCPLGEDE
Leptinotarsa  CSFGQKACRDGSKCLPKHLVCDNVAHCDDGSDEVFCTCKERVGKLRNCDGYFDCPEGEDE
Rhynchophorus QIELLYQRSALGPALLS...................CAVDRFGKDAATIFFKLSFN....
Periplaneta   CRPGTKACWDGLECVSSSEWCDGIVHCNDTSDERDCRCQDRLDKNRMCDGYFDCPQGEDE
Blattella     VIHPFVRPDIPSSTLKTPSNVNKVHTSFSQLHRNTSGYIATFGSERTRPHLYITSAPQTT
```

```
                      960       970       980       990     1000        1010
Drosophila      LGCFGCESLAYSCYENPQDFAKRNRSTISMCYSRLERCDGFMNCLNG.RDEEQCSMLVTD
Ceratitis       LGCFGCNDLIYSCYDGPEDYAAYNRSTVSMCYSATERCDGVMNCLNG.RDELECNIIVDN
Spodoptera      MGCYGCSETMFSCEDLDISAQST.......CFSKEQRCNNVVDCPNH.KDELECNMLAPS
Plutella        MGCFGCSEDTFSCEDLDLNSRSS.......CFSREQRCDNTADCPNR.KDELECNLLAPS
Aedes           MGCFGCDKFSFSCFNTFFEYQASHHSET.RCFTLIEKCDGFNNCMNR.KDEQDCTMLVRD
Anopheles       MGCFGCEKFSFSCFHTEAEYRSAHRSGS.MCYSLLEKCDGFDNCLNR.KDEQDCTMLVRD
Culex           MGCFGCDKFSFSCFSNAAEHDAAHKSGA.MCFTAIEKCDGFDNCLNR.KDEHDCTMLVKD
Leptinotarsa    LGCFGCDEDEFSCDDWSKLRPST.......CIPIRQRCDNIRQCEITGKDEEDCSALADN
Rhynchophorus   .RRKLVENLTNIEKAIRDIFLSDAFSKKPIFRNIRFDSKSILVKQLQ.EGASVYSAPSAV
Periplaneta     LGCFGCNDQSFSCENWDRSHTATT......CMSVEHRCDGTRNCPNG.KDEAECLLLEDK
Blattella       WKTSLLPLTTPTKRITTMISSPTTSSIKPIPITSTSKPKTTIVTSTT.TSISTAKPNIAT

                     1020      1030      1040      1050                1060
Drosophila      VADHMSHGASASEGYLYHNYRGDWHPVCNNGEKWAALACQMDENSRM........DHSAS
Ceratitis       VLDHMSHAASSSSGFLFHNYRGEWFPVCNNAQKWASEACAN....EA........EHSAT
Spodoptera      LLKKPLFAISNTEGFLHRNFKGDWYAVCKNPYMWAHDACRRETGVIIRPPFISTVPIDPM
Plutella        LHDKPMFAVSNTEGFLQRNFRGTWFPVCNNPYMWAHDACRRETGLIIRPPYIQMVPIDPR
Aedes           LRSPLAFAVGHSVGVLHRNYKGKWYPVCHNPLNLAREACEAELG...........PADRD
Anopheles       LGHYLAYSVPHSTGVLHRNYRGKWYPVCHNPTQWAREACETELG...........PLDRE
Culex           LGSPLAFAVGHSHGILHRNHKGKWYPVCHNPTVLARDACESEIG...........PIDHD
Leptinotarsa    VGNQPQNKISNAVGFLHRNFRGKWYPTCFGTELWALEICKTESGPSSINPRSHMMLTMNN
Rhynchophorus   TSPPPSFKSNVKSGKMGVVLKSPKNMTVSPSTTSSKPIIKD.............EPEFNE
Periplaneta     LFHHQLFPVSYTKGFLHRFWQGKWYPMCNTLEEWALEVCKSDVGFFENKPDIEKRYVTSE
Blattella       STTKPKIATASTVKPIAATIMSTKIPKITKPTTSEMPTTKSNKNIST......AKPETTV

                    1070      1080       1090      1100          1110
Drosophila      LNVSFQTLTLPG.PFIEPSLHAGVHFAQACHGRNSHDSLVD....HVAYVKCP.PMQCGL
Ceratitis       PNVTFREITLSG.TFIEPTLVGQARFPQSCQQRDAHDKLVD....QAAYVDCMSTTQCGL
Spodoptera      VKIDYLNTGPGGLIQTSDTCMNSSAVYVTCPDLICGTRVQS....TSQLLKQSNVMENRL
Plutella        LKVTYLNTGPGG.VQMSDSCVNSSAVYVTCPDLLCGTRVLT....TPQFLRENAAFENHL
Aedes           PVILQHHGDLPG.PFIQPSPRSHHVFQPEFT.....DTCNG....LINYVKCP.APKCGS
Anopheles       PLLSHGHGSLPG.PYISQRANS.AVPQPEFS.....EACNG....VYVHVKCP.AVRCGT
Culex           PQITQRQGHLPG.PFIQPSLQSPHVFHPEFL.....ETCNG....FFTYVVCP.QPKCGT
Leptinotarsa    YQGLFINILPNNEVSLVKTCVQDRAAFVECPPMYCGLRMTMKNPYRPQEVDISVENIIGD
Rhynchophorus   EDLPVIQGSFKISKTEADITEKKTEAPVQKVKIEKTTSAPT....KKTTFKESSVT....
Periplaneta     YRGLFLMVDSHEKVELVKSCIGNEAIYVTCPRMFCGTRILPGVKDEEELSTEDSDLNLGR
Blattella       ISTTKPRTTITVSSTSKHKTSTSTIKPKITATLVTRHSTST....PKTTVSPTKTTTVEI

                           1120      1130                           1140
Drosophila      PSK...........SSMLEHSKRVRRAVSDSK.....................EIVGDGR
Ceratitis       VKKKVKTPFRRKRSGSLSKLHKSIRQQQEQRK.....................RENFNGR
Spodoptera      FGRNKRNFPARHAYPPLFYGNRFKRDVDQDSTKTNYGMG...INRFGAFDTMRNKRTESR
Plutella        FGRNKR.FLLRNPRYSIVYFNRSKRHVNSTFSRNSVAEENV.PYIDGGIRGWRDKRAETR
Aedes           SKQ...........NEMENLRIKIRGKRN.........................ATELVQ
Anopheles       SRM...........HEQHAARINVRTRREANES....................EIVESVR
Culex           TKL...........NEFPSARIKIRNKRANPD.....................EVVESTR
Leptinotarsa    LERDARLPNEGDDKHVETNMDEDAVSRFEEEKEKN..................LTLGTPR
Rhynchophorus   .......................................................SALYK
Periplaneta     IKRLENIVQESLNVNADEDSSKLLRFHLMIPDSNNSTSETLGHEDGDAAHKQRPIRDEGR
Blattella       LRKSNTTVLLMKEPDMTVTTLKSEVSTIKPKTSSFSALT...............VSSLVR

                  1150      1160      1170      1180      1190      1200
Drosophila      IVGGSYTSALQWPFVVAIYRNGKFHCGGTIYSDRWIISAAHCVINYGKYFYEVRAGLLRR
Ceratitis       IVGGSYSSPLQWPFVVAIYRDGKFHCGGTIYSEKWILSAAHCVINYHKYYYEIHAGMLRR
Spodoptera      VVGGRPSQPAAWPWMVAVYRNGMFHCGGVVINHSWVISAAHCVSKFWKYYYEVQVGMLRR
Plutella        VVGGRPSQPAAWPWMIALYRNGMFHCGGVVISQKWVVSAAHCVHKFWEHYYEIQVGMLRR
Aedes           IVGGTKAEPAAYPFIVGIFRDGKFHCGGSIFNEHWIVTAAHCCDNFPRHHYELRAGLLRR
Anopheles       IVGGSHADPEAYPFIVGIFRDGKYHCGGSIYNEHWIISAAHCCDNFDQHYFEVRSGMLRK
Culex           IVGGSYSEPSAFPFIVAIFRDGRFHCGGSIFNEHWIISAAHCCDNYHKHYYELRAGMLRK
Leptinotarsa    VVGGKPSQPAAWPWLVSIYKNGIFHCGGVLINELWVLTAAHCVDKYSLYYFEIEAGTLRR
Rhynchophorus   IAGDKTKVGVTATTTTTSTKTPATTKQAATTTSSTTTTSTTTTSRSTTVPIKEFKGSPE.
Periplaneta     VVGGTASKPGAWPWVVALYRDGQFHCGGVLLQESWVMTAAHCVDGFSQHYFEVQAGMLRR
Blattella       MAQSPEKLTSEATTELSTKQTTSAVKNSSSIPTTSTTMSSNVELSAATNFVPPNISLPEN

                  1210      1220      1230      1240      1250      1260
Drosophila      SSYSPATQIQPVSHVVVHQAYERRSMRNDLSLLRLLNPLQFNRWVKPICLPDKGRTTVGD
Ceratitis       TSFAGSTQIRTVSHIIVHQAYERRSMRNDLSLLAMAEPLKFNRWVKPICLPDVGRTTAGA
Spodoptera      FSFSPQEQSHRVTHVIVNQFYSQTDMKNDMSLLRVEPAIQFSRWVRPICLP..GPDTAGP
Plutella        FSFSPQESNHRVTHVIVNQHYDRVDMKNDLSLLRVTPSIQFSRWVRPICLP..GPDTAGP
Aedes           RSFSPQVQVSTVTHVFIHRGYSAQKMINDISLMHSDRPFQYNRWVRPICLPDRHMTTNDR
Anopheles       RSFAPQVQITRVTHMIVHHAYSSSLMANDIALMRVEHPFHYNRWVRPICMPERHRTTDDR
Culex           HSYAAHVQVSLVTHVIVHQGYSAEKMINDISLMRSDRPFGFNRWVRPICLPDRHRTTNDR
Leptinotarsa    FSHAPMEQHRWASQIVPHESYDKSKLRHDIALMKLSSPVRFNRYVRPICLP..SEATAGR
Rhynchophorus   .IYNDEPWIPILPN................IPVYPSKTVPINDYNQPS............
Periplaneta     FSFSPAEQTQPVEHVILFERYDRSDMRNDLALVQLREPLRLNRWVRTVCLP.........
Blattella       LITNAIPSVPVTEQSTKISEIVVTPSK...ILTEPTRMTTTAQYVSPSSNRNSTLPLKTK
```

```
               1270      1280      1290      1300      1310
Drosophila    DWIWGPVEHTLCTVVGWGAIREKGPSSDPMRQVIVPIRKKCTDPED.........QASED
Ceratitis     DWIWGPEENTLCTVVGWGAVREKGPGSDQLREVIVPIRKQCTELED.........KESED
Spodoptera    DWLWGPAPGTICTAVGWGATTEHGPDPDHMREVEVPIWENCKHRED.........QTGKE
Plutella      DWKWGPPSGTSCVAVGWGATVEHGPDPDHMREVEVPIWGQCKHRED.........RAGKE
Aedes         DWIWGPKPGTMCTAVGWGALREHGGSPDHLMQVTVPILPFCKHKND.........RDGLA
Anopheles     DWIWGPKAGTVCTAIGWGALRERGGAPDHLMQVSVPILGYCKHKSD.........RDSLQ
Culex         DWKWGPKPGTLCTAIGWGAVREHGGAPDHLKQVTVPILSYCKHKSD.........REGLE
Leptinotarsa  DFVNGPAVGTICTTVGWGATFEHGSDPDHMREVEVPILKHCTRPED.........ITDDE
Rhynchophorus ........PPTYTSFTNPGLSYHFQEAEQLGTTGIKIHP.....................
Periplaneta   PEVWGPRSGTVCTAVGWGATFEHGPDPDHMQEVEVPILPKCKHAAD.........NKGFE
Blattella     NVVLTTIFPETRLAISNKMENIKNSSMELLGNSSLYTKGNYKFNQTTGTQNDTKVEENKQ

              1320      1330          1340                 1350      1360
Drosophila    ICAGDPDGGRDACQG....DSGGPLFCRSVSNPDE..........FYLAGVVSHGNGCAR
Ceratitis     VCAGDADGGRDACQG....DSGGPLFCRSVNNTEE..........WYLAGVVSHGNGCAR
Spodoptera    ICAGFVEGGKDACQG....DSGGPLLCRNPLNSQQ..........WYMAGIVSHGDGCGR
Plutella      ICAGPVEGGKDACQG....DSGGPLLCRNPLNAQQ..........WYMAGIVSHGDGCAR
Aedes         ICAAEMSGGHDACQG....DSGGPFACISVSSPHE..........WYLAGVVSHGEGCAR
Anopheles     ICAAEEDGGHDACQG....DSGGPFVCQSKSNPFE..........WYLAGVVSHGEGCAR
Culex         ICAAEEDGGQDACQG....DSGGPFVCVSITNPHE..........WYLAGVVSHGEGCAR
Leptinotarsa  LCAGLSEGGKDACQG....DSGGPLMCRNPNNPSQ..........WYLAGLVSHGEGCAR
Rhynchophorus IPVNK.............IPFTKPFVDLTTTETSK.........................
Periplaneta   ICAGVPEGGRDACQG....DSGGPLLCRHKG..HQ..........WYVAGIVSHGEGCAR
Blattella     IIVEKEDKPHGQEDNRTVIDHTNPFQDSLANSSSKQFQKILLKDPLHVSSVSENQNKFSL

                   1370      1380      1390      1400      1410      1420
Drosophila    PQEFGVYTRVTLYLDWLEMATTPRLLPKLQPLQLCPGFICVWGGKRCIAKRQRCDRNVDC
Ceratitis     KGEFGAYTRVALYLDWIEMASKYR..TKLQPRLTCPGYTCVWGGNRCIPMKRRCDRIVDC
Spodoptera    KDEPGVYTRVSLFVKWIKYYISSNTLPTIQPIQECPGFKCDTGISKCLPLKRRCDKIVDC
Plutella      KGEPGVYTRVSLFVKWIKFHIFSKTLPGIQPKQECPGFKCISGIAKCLPKKRMCDKIIDC
Aedes         PNEPGVYTRVALFNDWIQRKTREVL.TSSSTRQDCPGFQCSVGVSFCIPRQKRCNGKVDC
Anopheles     AHEPGVYTRVALFIDWIAEKVNAPL.PARTARADCPGMRCIWGGGICLPPGKKCNGYVNC
Culex         ANEPGVYTRVSLYIEWIHQKVHEPP.PHMTARLTCPGFHCTWGSPVCLPVEKRCNGVVNC
Leptinotarsa  PNEPGVYTKVARYVGWIAENMGDFKPHLVIPLQHCPGYTCKFTN.RCLPKKRHCDKIVDC
Rhynchophorus .....................QKQTPTEITTDFVNIKYQAENEYKKDEEYG...IKNISS
Periplaneta   PNEPGVYTRVMLFLDWIKEKVGTTQLPDKTPRPSCPGLQCSVGNARCLSKMKVCDRVVDC
Blattella     KNQSVQILNNTTDNYQILITTQKATATTPNQEYRNSTVVTAESYNQRIKIDGDVVANNTT

                   1430                                       1440
Drosophila    LGGEDEVGCT.................................YNFLPDMVGGVRQNIST
Ceratitis     LGGEDEVGCI.................................YNFIPDLVG.SSKNISS
Spodoptera    LNGEDETRCD.................................FTKASF.TDSLLMETKN
Plutella      LDGEDEMNCN.................................FDSPLHGADKNMFISEN
Aedes         LGGEDELSCS.................................LDQLLAESIQETTIATT
Anopheles     LGGEDESGCG.................................MDQMLRSMSQRGEEESE
Culex         LGGEDELHCP.................................LDQMTGIFLNRTTTESA
Leptinotarsa  LFGDDETNCK..................................HTFQNIFKHARGKTHH
Rhynchophorus IFYDLASSLD.................................TENITIVEREDYEPDLK
Periplaneta   LRAEDEMNCPSLSSRSMGSSHLNSRAEIENEDMFGRNYLSNDILASENQSGGAIGFDNHA
Blattella     YLKENSSEAN.................................NRNISVDSGISSEKVHS

            1450      1460
Drosophila    TTESDYHPVKESEEKSKMRE........................................
Ceratitis     TTESDYHP....EEVVKTTH........................................
Spodoptera    TLEEL..MERKEAETEAPLMSTASNQENMMSTRMPMDKETPKPQLD..............
Plutella      TVEKRNNFETKEAETESPTVKDTTQG....SKDIIEDTDT....................
Aedes         PKNSTATSSTTAAATTEAVTSKIDFLAENSEASDPATTVTEASSTETAT...........
Anopheles     EEGGSATQRASDVDTTEAET........................................
Culex         ANATTSVPTTSKSSTTKPAK.........ARAEEP.........................
Leptinotarsa  TTLIAKKTNGQQPGLVSPED........................................
Rhynchophorus NIAGQGHVEVVEVDMEELLG........................................
Periplaneta   ALEDNTYLQRETERTSESLQRRKIIRDTTNSGSVDADGRKVGGILQSLTGPAAYIFSNNE
Blattella     KIEEMSITEASLASNINIED........................................

Drosophila    ............................................................
Ceratitis     ............................................................
Spodoptera    ............................................................
Plutella      ............................................................
Aedes         ............................................................
Anopheles     ............................................................
Culex         ............................................................
Leptinotarsa  ............................................................
Rhynchophorus ............................................................
Periplaneta   AQVPPLPYIRSKMERLTRKDNVVWDNSTGNTTYHNGTKNETFTDGEMKGIIVSNSETNNG
Blattella     ............................................................
```

```
                                                                1470
                                                                   .
Drosophila      .....................................................VIPIDDE
Ceratitis       .....................................................CKAADT.
Spodoptera      ..............................NENPLSSTENILGMSTNGNRGNTVSTMMAF
Plutella        ...................................SETNNIR..........TVSHQATM
Aedes           ..............IETMSATLETVSAKDLSDDIAQISTAATTVEPNTTLDMTVSSSTQS
Anopheles       ....................................................TVLFTSEE
Culex           .................................VAQLS..............AIASSDII
Leptinotarsa    ......................................................TPILGM
Rhynchophorus   ....................................................QTTKIPLV
Periplaneta     NITSDSDIWNKTLLEEEEAAQIETFKNISYNEGQDAVTETKKTTIDNGTQNNSVSDSDSN
Blattella       ....................................................KSISTHFK

                 1480
                    .
Drosophila      DLKAEQ....................................................DE
Ceratitis       ARNAGV....................................................EE
Spodoptera      DEETPKPQLDDDLSSAELHLHKHGW...........................ESKKIIKQ
Plutella        RDNS......................................................DS
Aedes           VSTSPE....................................................KV
Anopheles       TTTLSV....................................................PE
Culex           VSPVDD....................................................RT
Leptinotarsa    NLKSEE.....................................................N
Rhynchophorus   TLLPVR.....................................................S
Periplaneta     NKTVNDSETSNKTISVNYKMCNSETGNNTLCDKEMENKTLNLILFTEKHDEGIGTVEKII
Blattella       VMSPITRTSEHTTVPAIYPSVIKVK...........................ESQRQPKM

                  1490      1500          1510      1520      1530
                     .         .             .         .         .
Drosophila      EDLLKSTTSLGQTETTQGPM....DLSFAEQITSTTSDDLSITDETTSTDFTVSDSATSP
Ceratitis       ENEVEVDDEKASTERRVSDD....EESITNASTATITQNITKIESSTN...DATEYTTDF
Spodoptera      EDIISKENADDQSKTVDETP........DSSQTMFENNDRSSEGVVSSKDNAVTESVVKM
Plutella        AGTVSRGDIETISISVTKDP........LESRSMMLDSQKKNE...ELRESDIPENFDDF
Aedes           EESTMDESTSSTTETSFTHPTTMNSETTTNNATTLISETTLPSLSTKEDNATSVDYSATI
Anopheles       EASVEAQESMIMTNTAQTEP........ITTTTSTVATTTPQLELSTAEESTTEDS..ST
Culex           PNSEVNEQTT.TYKSTLTN........TTSRSVTTVQQTTLP..TTTGKSATSVNH..TT
Leptinotarsa    NSQEHTTSEPQNSTSLWEVP........EKKQIDVTDEATKEKNVSKDDKQVSKKRIGSL
Rhynchophorus   NSGIGRPFNRKRFGDSNNLSVE.....NRSFPSHAVKQTLDTFEKPKNKSKTVPEPVNIS
Periplaneta     SDNETQKNTASSSEKGNDGPLKNDIENKTETDREMHNDTVSVLNETRALPENVEITTVSV
Blattella       KSYIGAPEFEQKGKDFLSLEDS.....IKDKGNHLKTDSLTSVINKSNKTIEVALTLTTQ

             1540                                        1550      1560
                .                                           .         .
Drosophila      STLLPTTTN................................PSTWLPSTNIETSTFSFTT
Ceratitis       STTESSDID................................ETNATPQATFADLTTSETI
Spodoptera      TTVSRGDVELITM.........................GFDPASLESGSSVLETTVNYIP
Plutella        TTITTQ...................................SPEKNSITTISDTTEDFS.
Aedes           STDQTTHTPCTTSPAADSLDDSSSAHSSPANDSEPSTESSVHATTEFTTTVIELTSVEST
Anopheles       SASTSEDAP...............................TEASTSPESTTLEVTTDLST
Culex           VSDTSIHT...............................STIRTREATEAVIGVAKNLST
Leptinotarsa    TNRLHSDIK...................................................
Rhynchophorus   SADNY.......................................................
Periplaneta     DGTQAQEINKNTIDMNETRSRIVGNNKTINEKVSDSEKQSTAGSVKEINDITENTFEDEA
Blattella       DPNNFNYSITLPSTVFKKLNESVKSTKESELVGHDKQEEIEEEIVTSEKIDSETYEDYHI

Drosophila      T
Ceratitis       S
Spodoptera      P
Plutella        .
Aedes           T
Anopheles       E
Culex           T
Leptinotarsa    .
Rhynchophorus   .
Periplaneta     V
Blattella       S
```
