## Supplementary Alignment Files for "Assessing target genes for homing suppression gene drive": nox.pdf

```
                1       10        20        30        40        50
Drosophila    MNADQESN...NHRGSVSSSRSLEIPATPSRSPKKVSFSDELPQSQTPAQQPINQ.....
Ceratitis     MDHNQKNNTSENPRGSVSSIKSLEVPSTPTRSPKKVSFSDDLPFSETTDSIAVVQNLSSS
Spodoptera    ...........................MASNGKTPEILTNFET.VDLDKNGD........
Plutella      ...........................MAENGTSQPNDPEKETNRTIDNQDE........
Aedes         MENPSTSSEIPKRHSLTVSSESESETVKLSDSRSVSPQLPHKAPKKVSFSDELPQESSTF
Anopheles     MEG.SVPNSPSRPTSPVAAGESESEPTGVSGSRSPSPQPGHKQPKKVSFSDELPQSGATV
Culex         MNPVPSTSSGAKRLSLSVTNENESE..SAPNSRSASPQLPHKAPKKVSFSDELPQESAGM
Leptinotarsa  MSMAQPLVGTSVPPIPEESAESTPIVPRRKSSTKP...LDISSVKTTSNPENISDG....
Rhynchophorus MASLTGYENTEETASVLGYNKGVDGTDRRKESVMQNGQVEINGNVIKLNKNQLGDV....
Periplaneta   ............................................................
Blattella     ............................................................

                                             60        70        80
Drosophila    .........................QSAPTVVSHVLQQAAQYLERLHGARDSAEEKTEEV
Ceratitis     NNNDSSRDNNVASSLSNNKSPTIVTEEELTAVNHVLLQAAQYLEQMHGTRERQSPTVEAT
Spodoptera    .........................EDVGKNGQLVVSAAN...GQFDGN...........
Plutella      .........................KEETKNGGFLLSVDSSDGGNTDSSNVN........
Aedes         AGTSA.....PDSTVTQKSESVKLAHGDVNPFVFVLQKNDSYLKNLHQIDPS.PS.....
Anopheles     QQPEK..........CAPNESESGRPQEENPFIFVLRQNDSYLQDMHSSAQS........
Culex         SQSSAT.EGGPSAPLTPKSESVKLSHFDVNPFVFMQQKNDSYLNNMHQMDEARAE.....
Leptinotarsa  ...................ETVISCSETPEPNPAERRRSSGTLAKLVKQKKV........
Rhynchophorus ...................ENNVIK.ENPKRNSLESSTQTLDLIRDQQFDQK........
Periplaneta   .............................................MTEPAEA........
Blattella     ............................................................

                                                90
Drosophila    QP...............................ERADIRDS...................
Ceratitis     EPTHSPPQSHHHSDGDISTTSVHINLKQPPQKVRRAEKRDYGGGNGDGDSGAALSNGRVQ
Spodoptera    ............................................................
Plutella      ............................................................
Aedes         ............................................................
Anopheles     ............................................................
Culex         ............................................................
Leptinotarsa  ............................................................
Rhynchophorus ............................................................
Periplaneta   ............................................................
Blattella     ............................................................

                   100       110       120                    130
Drosophila    .....SDTSEAAVLDLDALDASADEPAANAG.............PIVASSPSIL......
Ceratitis     EINICDEFDKVQVHKLPNMNTSANDIATTATTTTTKAAPELKIAAAVSTSPAVLKLDDAD
Spodoptera    ............................................................
Plutella      ............................................................
Aedes         ..........IAKPLTDSDVFPNQRRASTPGPNESETVVD..........LEPKPSASIL
Anopheles     ............PSGGGGGVFPNARRSSLHS.NESETGSSTGSGTSTVTATDTGGDNSGV
Culex         ..........PAPPLSTTDVFPNQRRASTPSTSSADESVTPPIGGDLGVKVTPQP.ASIL
Leptinotarsa  .................DGTVGFIN.........................NGFS...ADV
Rhynchophorus .................SGAVQSRS.........................IGVGTHIDIV
Periplaneta   .................GGPVPQQG.........................VKME......
Blattella     ............................................................

                    140       150       160
Drosophila    ..IGSVNRQEAIGSANGNGNANPNQRQSN...............................
Ceratitis     DEFNSIEQSKTLLSQQQQQQQQQQQQQQTKSQPIVSINTTQPATTTAATRKTFTTSLSNL
Spodoptera    ...................CLGVPRTGHR...............................
Plutella      ..GIKMADLKQNGDANPMMLLGVPRTGHR...............................
Aedes         KPSDPVPKSVSPFDPISAGLVNDLKTQRDLVDAAL..VPAG...................
Anopheles     VQQDKSTTELVPFDPIAAGLVNSLGGGSDEPAPPP.........................
Culex         KPSDPAT.....FDPISAGLVNDLRAQNDLVNAALGATPKGGSPT...............
Leptinotarsa  SDAATLSDVPSKASVVDERLAQRRKSLVP...............................
Rhynchophorus EEVPSCSKSASCALPVDEGTIIRRRSLVP...............................
Periplaneta   ....EIPKDNVTDVEVKRPLPHTPNTSFK...............................
Blattella     ............................................................

                                                     170       180       190
Drosophila    ..............................LYMERYNLNLDKNCSSMELEARREKQRWLL
Ceratitis     QLEPVVQAEPTQPTTSAQMQEISEEQSHVARYIDRYNLNLDKNFSAMEVEARREKLRWLL
Spodoptera    ...........................................PSIMEVECAKERTRLAL
Plutella      ...........................................LSIMEMESARERQRVSL
Aedes         .................KKKNLADSDKIEKSVLGSGSEMPPPVVSAMELEVKRDKRRWLL
Anopheles     ......................ASTEPRKPS............VSAMELEVRRDKRRWLL
Culex         .................KKQNLADSDKIEK.....NGTVP...VSAMELEVRRDKRRWLL
Leptinotarsa  ...........................................MSSMEEENLRERLYLAQ
Rhynchophorus ...........................................MSSMEEENLRERLHLAH
Periplaneta   ...........................................MSSMEEESIREKLRTSL
Blattella     ............................................................
```

```
                      200       210       220       230       240
Drosophila     ISECSALFDEGEG.KHTREAFRKLFLDEEFQQKLFQLFDLERNGYLLQDRWIEHLKGRLT
Ceratitis      ISECSALLAEG...KHTREAFRKLVLQREFQQKFFHLFDLEGNGYLVQDRWIEHLKGRLT
Spodoptera     LKQCSAILKQG.DQKYTKESLHRTFQDEDLLERIFRLFDADGNDNLDQNDWLEFLKERLS
Plutella       LKQCSAILRQGEDQKYTKESLHKTFQDEDLLERIFRLFDVERRDYLVQEDWIEFLKARLT
Aedes          ISELSAICGEEK...HTVDGFKRIFREQTFLQNLFRLFDMDGENCLAQDKWIEHLKGRLT
Anopheles      ISELSAILGEDK...HTIDGFKRIFREQTFLQNLFRLFDMDNENFLVQDKWIEHLKGRLT
Culex          ISELSAVFGEDK...HTVDGFKRVFREQTFLQNLFRLFDMDNENCIEQDKWIEHLKGRLT
Leptinotarsa   LRLCTDIVNKEA..KYTRDTFRRLFQSKELLEQLFKLFDQDRDHCLHQEDWIEFLKQRLT
Rhynchophorus  LRLCSEIVNKEQ..KYTRDGFRKLLQSKDLSELLFKLFDQDRSQTLPQEQWVEFLKQRMT
Periplaneta    LKQCSDILKGDGATRYTRDAFRRLFQDKELLQRVFRLFDQDRDSLLVQEEWVEFLKERLT
Blattella      ............................................................
```

```
               250       260       270       280       290       300
Drosophila     DDRQMDFAEQIESVAYVICGENKRVSFKNFRDIWHTRGILDKLYRLIELDGSNLVSTNQV
Ceratitis      DDRQMDFAEQLESVAYVICGENSKITPKNFTEIWQARGILDKLYRLIDADGTNLISTNQI
Spodoptera     EDKQIDFVEQVESVAYVLCGNGV.ITLDTFQLILKNKVVTNKLFRVVDTDGDGYATADEI
Plutella       EEKQLAFVEQVEAIAYVLCGDGE.LALPTFQTIFKSRVVTDKLFRVLDP.GDNHTTAELI
Aedes          DEKQMDFAEQLESVAYVLCGEGT.VSYEKFCQIWHAKGILDKLYRLIDVDSTNLLSTNQI
Anopheles      DEKQIDFAEQLESVAYVLCGDGT.VTYDKFCQIWHAKGILDKLYRLIDVDCTNLISTNQI
Culex          EEKQMDFAEQLESVAYVLCGEGT.VTYEKFCQIWHAKGILDKLYRLIDADSTNLLSTNQI
Leptinotarsa   NEKQIDFAEQLESVAFCLCGEKP.IFLDQFYQIFHAKGILDKLFRLIDEENSGVISSDAI
Rhynchophorus  NEKQIDFVEQLESVAFCLCGDEP.IKQHQFYQIINAKGISDKLFRLIDEDNTGAAISENI
Periplaneta    DEKQIDFAEQVESVAYVLCGDDA.ITLEKFYQIFQAKGIVDKLFRLLDQDLDGIVTPDQA
Blattella      ............................................................
```

```
               310       320       330       340       350       360
Drosophila     MEFISHLTNSRPRTGFDKSSLARLEQLFRTTVGNEQEIRREEFQKIVTSKNPFFTERVFQ
Ceratitis      MEFISHLTNSRPRTGFDKSSLERLEQLFRSTVGNEQEIRREEFQKIVTSKNPFFTERVFQ
Spodoptera     MEFLSTVTSSRPRVGFDKHSVDRLEKLFRATVGDEKEITREQFQKIVVSKNPFFTERVFQ
Plutella       MDFLSTVTSSRPRVGFDKTSVDRLEQLFRQTVGDQREITKEQFQNIVVSKNPFFTERVFQ
Aedes          MEFISNLTNSRPRTGFDKMSLERLEQLFIKTVGNEKEIRREEFKKIVTSKNPFFTERVFQ
Anopheles      MEFISNLTNSRPRTGFDKSSLERLEQLFIKTVGNEKEIRREEFKKIVTSKNPFFTERVFQ
Culex          MEFISNLTNARPRTGFDKSSLERLEQLFIKTVGNEKEIRREEFKKIVTSKNPFFTERVFQ
Leptinotarsa   MEFLSTITNTRPRTGIDKGSLDRLEQLFRETVGNGKEIKKEEFKNIVTSKNPFFTDRVFQ
Rhynchophorus  MEFLATITNTRPRTGIDQNNLARLEQLFRQTVGKGTEIRREDFQKIVSSKNPFFTERVFQ
Periplaneta    MEFLANLSNARPRTGIDKANLEWLEQLFRQTVGNEREIRRDDFKKIVISKNPFFTERVFH
Blattella      MEFLASLSNARPRCGIDKGNLEWLEQLFRQTVGNEKEIRRDDFKKIVISKNPFFTERVFQ
```

```
               370       380       390       400       410       420
Drosophila     IFDKDNSGSISLQEFIDAIHQFSGQSADDKIRFLFKVYDIDGDGLIQHKELHDVIRHCIK
Ceratitis      IFDKDNSGSISLQEFIDAIHQFSGQSADDKIRFLFKVYDIDGDGLIQHKELHDVIRACME
Spodoptera     IFDEDDSGAISLQEFIAAVHQFAGQTPEDKIRFLFKVYDLDGDGLIQHRELQHVMRACME
Plutella       IFDEDDSGSISLQEFIAAAHRFAGQAPDQKIRFLFKVYDLDGDGLIQHRELQHVMRACME
Aedes          IFDKDNSGSISLQEFIDAIHQFAGQSPEDKIKFLFKVYDLDGDGLIQHRELQHVMRACME
Anopheles      IFDKDNSGSISLQEFIDAIHQFAGQSPEDKIKFLFKVYDLDGDGLIQHRELQHVMRACME
Culex          IFDKDNSGSISLQEFIDAIHQFAGQSPEDKIKFLFKVYDLDGDGLIQHRELQHVMRACME
Leptinotarsa   IFDKDNSGSISLKEFLDAMHQFAGQTAEDKIRFLFKVYDLDGDGLIQHKELQHVMRACMV
Rhynchophorus  IFDKDNSGSISLQEFLDAMHQFAGQSPEDKIRFLFKVYDIDGDGLIQHKELQHVMRACME
Periplaneta    IFDKDNSGSISLQEFIDAMHQFAGQSPDDKIRFLFKVYDLDGDGLIQHRELQHVMRACME
Blattella      IFDKDNSGSISLQEFIDAMHQFAGQSPDDKIRFLFKVYDIDGDGLIQHRELQHVMRACME
```

```
               430       440       450       460       470       480
Drosophila     ENGMEFSEDQIEDLTSAMFEDADPHNSGEITYEALKNQLHKHGGLLENLSITIDRWLVPI
Ceratitis      ENGMEFSEEQIEDLTTAMFEDADPYNRGEITYEALKNQISKHGGLLENLSITIDRWLVPI
Spodoptera     ENGMQFSDDQLLELTTAMFDDADSERRGAITYDALKKQLESHDGLLENLSISIDRWLVPP
Plutella       ENGMQFSEEQLLELTSAMFEDADTERRGAITCEALKVQLEKHGPLLENLSISIDRWLIPP
Aedes          ENGMRFSEDQIEDLTMAMFEDADKFNRGAITYEALKSQLEKHGGLLENLSISIDRWLVPL
Anopheles      ENGMRFSEDQIEDLTMAMFEDADKYNRGAITYEALKNQLEKHGGLLENLSISIDRWLVPL
Culex          ENGMRFSEDQIEDLTMAMFEDADKFNRGAITYEALKSQLEKHGGLLENLSISIDRWLVPL
Leptinotarsa   ENGMQFSEEQIDNLTMAMFEDADTNNRGAITYEALKSQLEKHGGLLENLSISIDRWLVPP
Rhynchophorus  ENGMQFSEEQIEDLTVAMFEDADTCNRGAITYEALKNQLEKHGGLLENLSISIDRWLVPP
Periplaneta    ENGMKFSEDQIDDLTMALFEDADPENRGTITYEALKNQLEKHGGLLENLSISIDRWLVPP
Blattella      ENGMKFSEDQIDDLTMALFEDADADNRGAITYEALKNQLEKHGGLLENLSISIDRWLVPP
```

```
               490       500       510       520       530       540
Drosophila     A.EDRQAGGAAKSGFWNSLPHQFSLAYMKNNQVFVTYLFFYITVNLCLFISRAIQYRASN
Ceratitis      T.QEKPKPKPGCPTCRITWPHQLTMAYMKNNQAFVSYLYIYIAINLCLFISRAIQYRASN
Spodoptera     ..KQDPQPVSFKQRLSRLKPYQLSRPYFKNNYVFLTYLGLFSLVNVGLFVARCVEYRKAN
Plutella       ..KPVPEKKSFYQKMVKLKPYFLSKQYFKNNYVFVAYLAVFTLVNIGLFVSRAIEYRHSN
Aedes          ...PQEDTTRKKHKKKPL.PHQLTAPYIKNNYVYLSFLSVFILINVGLFISRAIQYRKSN
Anopheles      ...PQEDTK.KKRKKKPL.PHQLTAPYIKNNYVYLSFLTVFTLINVGLFVSRAIQYRNSN
Culex          ...PQENVVRKKQKKKPL.PHQLTAPYIKNNYVYLSFLAVFTLINVGLFISRAIQYRKSN
Leptinotarsa   ..KPQSQPTSLLGKLLSYRPYQLTLPYIRNNHIYLIFLSGFLLVNAALFISRAIEYAGST
Rhynchophorus  PLKPNS.STSLFRRIISYRPYYFTIPYIRNNLVYLAFLLGFIMVNAALFVSRAFQYRESN
Periplaneta    ..KPKPPPRSLLGKMSTLRPYQLSRPYVKNNYVYLIFLTLFAIVNIGLFVSRSIQYRASN
Blattella      ..KPKPPPRSLLGKLSALRPYQLSRPYMKNNYVYLIFLDVFVIINLGLFISRSIEYRAHH
```

```
              550       560       570       580       590       600
Drosophila    GFVIIARACGQCLNFNCAWVLVLMLRHSLTYLRGRGLSSYLPLDHHVYLHKLTGITISVL
Ceratitis     GFVIVARACGQCLNFNCAWILVLMLRHSLTYLRSQGLSSYLPLDNHIYLHKLTGIVVSIL
Spodoptera    GFVILARACGQCLNFNCSWVLVLMLRHCLTELRVRGFSSYLPLDHHIYLHKLTGVLIAIY
Plutella      GFVIVARACGQCLNFTCTWVLVLMLRQCITWARARGAGAVLPLDYHIYLHKITGVMIAVY
Aedes         GFVILARACGQCLNFNCAFILVLMLRQCITFLRTRGFTAFLPLDQHIYLHKLTGVLVAAF
Anopheles     GFVIMARACGQCLNFNCAFILVLMLRQCITFLRTRGFTAFLPLDQHIYLHKLTGVLVAIF
Culex         GFVILARACGQCLNFNCAFILVLMLRQCITFLRTRGFTAVLPLDQHIYLHKLTGVAVAAY
Leptinotarsa  WYTIFARACGQCLNFNCTFIVVLMLRQCITFLRTRGYSSILPLDQHIYFHKLTGMFIFGY
Rhynchophorus WYTIFARACGQCLNFDCMFILVLMLRRCITFLRTRGYGSVLPLDQHIYFHKLTGWLIFGF
Periplaneta   GFVIVARACGQCLNFNCMFILVLMLRQCITFLRTHGFSVILPLDQHIYFHKLTGVLIVIY
Blattella     GFVIVARACGQCLNFNCMFILVLMLRQCITFLRTHGFSMFLPLDQHIYFHKLTGVFIFGL
```

```
              610       620       630       640       650       660
Drosophila    SLIHTIMHLFNFSIIVINDPNINAGHYTIGEWLLTDRPGLFGLIPGCANPTGVALLAILV
Ceratitis     SLLHTIAHLFNFSIIVVNDPKINAGHYTIGEWLLTDRPGLFGLIPGCANPTGLALLIIIL
Spodoptera    SLVHTLMHLCNFSIIVVNDPVLNKANYTLSEWLVTEKPGLFGLVPGCANPTGVALAVILL
Plutella      SLVHTVMHLCNFSIMVMPDPVINANNYTLTEWLLTERPGLFGLVGGCANPTGIALAVCLT
Aedes         SLLHTIMHLFNFTTIVVNDPVVNAKNFTTAEWLFTSRPGLFGLVGGCANPTGFALFFILL
Anopheles     SLVHTIMHLFNFTTIVVYDPVLNANNYTTAEWLFTARPGLFGLIGGCANPTGVVLVLILL
Culex         SLLHTIMHLFNFTTIVVNDPVINANNYTTAEWLFTARPGLFGLVGGCANPTGIALFVILL
Leptinotarsa  SVLHTVMHVCNFSLIVVNDPIINEANYTASEWLFTTKPGLFGLIGGVANPTGVALLVILI
Rhynchophorus SFWHSAMHVFNFSLVIVNDPKINSQNFTTAEWLFTSRPGLLGLIPGWANPTGIALFFILL
Periplaneta   SLVHTLMHLLNFSVIVVNDPVINKANFTLAEWLLTDRPGLFGLVGGVANPTGVALIIILI
Blattella     SVLHTIMHLSNFSLVVVNDPSMG..NYTLTEWLLTDRPGKFGLVPGVANPTGIALIIILV
```

```
              670       680       690       700       710       720
Drosophila    VMFVCSQPFVRRKGSFEVFYWTHLLYVPFWILCLFHGPNFWKWFLLPGLVYIVERALRFI
Ceratitis     IMFICSQPFVRRKGSFEVFYWSHLLYIPFWILLLFHGPNFWKWFLVPGLVYIVERILRFV
Spodoptera    VMFICSQPFIRRGGSFEIFYWSHLLYVPFWLLLLFHAPNFWKWFVLPGTVYLSERIMRLA
Plutella      LIVFCSRPAVRRGGSFEVFYWTHLLYVPFWLLLIFHGPNFWKWFVLPGAIYLAERIARLI
Aedes         VMFICSQPFVRRGGSFEVFYWTHLLYVPFWILVLFHGPNFWKWFIVPGFIYVVERTVRLV
Anopheles     IMFICSQPFVRRGGSFEVFYWTHLLYVPFWILVLFHGPNFWKWFIVPGLIYLVERTIRLV
Culex         VMFICSQPFVRRGGSFEVFYWTHLLYIPFWILVIFHGPNFWKWFIVPGTIFLVERTVRLV
Leptinotarsa  TIVISSQAFVRRGGCFEIFYYTHLMYVPFWILLILHGPNFWKWFFIPGIAYGLERIYRSA
Rhynchophorus VMFICSQAFVRRGGCFEIFYWTHLMYIPFWILLIIHGPRFWMWFIIPAIAYTAERISRVD
Periplaneta   IMFICSQPFVRRGGCFEVFYWSHLLYIPFWVLVIFHGPNFWKWFIAPGIIFLLERIFRFV
Blattella     IMFICSQPFVRRGGSFEIFYWTHLLYIPFWILVILHGPNFWKWFIGPGILYLIERLVRFV
```

```
              730       740       750       760       770       780
Drosophila    WMRGEHGKTYISSGLLLPSKVVHLVIKRPHHFNFRPGDYVFVNIPAIANYEWHPFTISSA
Ceratitis     WMKSEHGKTYISSGLLLPSKVIHLVIKRPFNFNFRPGDYVFVNIPAIANYEWHPFTISSA
Spodoptera    WMRSERGKTYISSGILLPSRVTHLVVKRPPLFHFHAGDYVFVNIPAIATYEWHPFTISSA
Plutella      WVRSERGATYISSGILLPSRVTHLVVKRPPLFDFHAGDYVFVNIPAIASYEWHPFTISSA
Aedes         WMRTEHGKTYISSGLLLPSKVTHLVIKRPFHFCFRPGDYVFVNIPAIAKYEWHPFTLSSA
Anopheles     WMRTEHGKTYISSGLLLPSKVTHLVIKRPLHFCFRPGDYVFVNIPAIAQYEWHPFTLSSA
Culex         WMRTEHGKTYISSGLLLPSKVTHLVIKRPLHFCFRPGDYVFVNIPAIAQYEWHPFTLSSA
Leptinotarsa  FIVTERGKTYISSGLLLPSKVTHLVIKRPLQFDFHPGDYIFVRIPAIAKYEWHPFTISSA
Rhynchophorus TMSS.KGKTYISSGLLLPSMVTHLVIKRPSQFDFHPGDYIFVNIPAIAKYEWHPFTISSA
Periplaneta   SMQSERGKTYIRCGTLLPSKVTQLVIKRPHHFDFHPGDYVFVNIPAIAKYEWHPFTISSA
Blattella     SMRSERGKTYISSGLLLPSRVTHLVIKRPHHFDYHAGDYVFVNVPAIAKYEWHPFTISSA
```

```
              790       800       810       820       830
Drosophila    PEQEDYMWLHIRTVGEWTNRLYRYFEREQQKLQQSGSSQE...IPQHMHAIP.......T
Ceratitis     PEQEDYMWLHIRTVGEWTNRLYQYFEKEQQKLHKGEIIQQRHAIPTDMVRITEKNESRTS
Spodoptera    PEQEEYIWLHIRGVGEWTNRLYSYFEREQARLHGEELPAIENGHQ...............
Plutella      PEQEEYIWLHIRGVGEWTNRLYEYFEREQTRLHLGNVAPHTDTHE...............
Aedes         PEQEDYIWLHIRGVGEWTNRLHSFFEREQERLHSGEIPAVIAGCS...............
Anopheles     PEQEDYIWLHIRGVGEWTNRLHNFFEREQERLHNGEIPALVAGSSGSASVAA........
Culex         PEQEDYIWLHIRGVGEWTNRLHGFFEKEQERLHNGEIPAVIAGCS...............
Leptinotarsa  PEQDDYMWLHIRGVGQWTNRLYEYFEREQEKLHNGEIPALPPSPS...............
Rhynchophorus PEQEDYMWLHIRGVGQWTNRLYEYFEREQVKLHNGEINRQACSSK...............
Periplaneta   PEKEDDICLHIRGVGEWTNRLYGYFEQQQEKLHN.VDGQLTG..................
Blattella     PEQEDYMWLHIRGVGEWTNRMYNYFEREHERLHS.VEDQLSGNMI...............
```

```
              840       850       860       870        880
Drosophila    PSFMLLNEARNPAIAGERSATPQTDFLAKNLGVQAVP..PVRPPRQNRKP..........
Ceratitis     PTPQIDFLAQNLARLQNKTNGNEPSLLLANKSANQMQNVPLKPARQTQVPS.........
Spodoptera    ......KLARSISSQSKKSRKSSVASTKKRS.............................
Plutella      ......KLSRTASNQSAR....SIKSHKKRS.............................
Aedes         ......TEPSSSSS..TQTVQEKPKEVPLKFSAGVPKSASVESPLAKTT...........
Anopheles     .....VAASAKATGGTVDVIQEKLKSTPVAT...IHKSSSYESALNKAKPSGVGGAGGPG
Culex         ......TSPCSSSSPTTEVVQEKPKELPLKFNTGIIK..TVESPLAKTP...........
Leptinotarsa  ......KKSPSNGGINNNQ..SPLKKLQTTITRTLS.NKDMT..................
Rhynchophorus ......ACDNTTEKTNNN....ALTQMKETITRTLSREKSVK...................
Periplaneta   ......NRPPAINGTPERTEPKGIRKIQASIQRKFADRDSSSSVRGKTID..........
Blattella     ......NRPMVINGTPE...PKGIKRLQASIQRKFTDRDSATSTRSKAVD..........
```

```
                     890          900          910
Drosophila     ...................APGAPIDPPATGVN.RIRS.......IKKTLQRTFSRKEAV
Ceratitis      ............KLALESISARDEIDTSSAKPNDRIRS.......IKKTLQRTFSRKDVP
Spodoptera     ......................VDFSPKCFTN..............DAY.VPEEGNGDVK
Plutella       ......................VDFSPVVYTN..............EAFSIEEECNGNGN
Aedes          .....ENTFDKPLVSKRMPPGTSKLAMESYTAPPPSAP.....PKFDRQLSENKAIKKIQ
Anopheles      GPRAGDATTPAGIMKQRHPPGTSKLAMEGYSAPSPVATSTAQSAKFERQMSDNRAFKKIQ
Culex          .....DNTFDKPLINKRLPPGTSKLAMESYTAPQPHSP.....PKFDRQLSENKAIKKIQ
Leptinotarsa   ....................KKPGIQLVAFSH..............EAFQNDERSKSPHG
Rhynchophorus  ....................KKSGVQLVGFSN..............DHF..DETSTP...
Periplaneta    .............FIKEGASPRTGYLNESSSS..............DSDFNLNSSMPHHS
Blattella      .............FMKHADSPRTGYLNESTSS..............DSDFNLNSSMPHHS
```

```
             920          930
Drosophila     DPKK..GIPNGAFIAD..............................GEREDS........
Ceratitis      TKNGSGGHTNDGFVAD..............................QVDSYQKPLDKRVL
Spodoptera     QND............................................APSPTKISYHPSL
Plutella       GVT............................................SPPHITPEKQMKL
Aedes          ATLQRTFSRKDQFRG.......AGVSNEAFTGD..DGLEE.......PKRKVTPEHNLHK
Anopheles      ATLQRTFSRRDQLIPRSGGAGGVGGIANEGFSGDGADGGADGSELKDAKRKVTPEQSLHK
Culex          ATLQRTFSRKDALRSN......GGVSNEAFTGD..DGLEEG......PKRKVTPEHNLAH
Leptinotarsa   ENDTNMASCSNT...E............................QTTKPRKLTPEKRLHR
Rhynchophorus  ......SDCNN....................................RSKRIIPDKRLQR
Periplaneta    DTDLSGPHPGNNDCTS............................LTGPPRKLTPEKKLHM
Blattella      DSDISGPHAGNADITS............................LTGPPRKLTPDKKLHM
```

```
               940          950          960          970          980          990
Drosophila     ..NLKQRPLEKSISLPDISVKSKKRSRLKALRALG..RSESESAFDEKRVRRARNNSVGL
Ceratitis      KLNSRKPQLEKSLSLPDIAVKTKKKERLKALRALG..RSESERSFDESRVRRARLQSVGL
Spodoptera     LLAPPRT.LEKSRSMPDVQKNLKKRQRLMALRDF..MRSESESKLDEGSMLSAR.KTFGL
Plutella       LAVPAKMPLEKSLSMPDMQTRLKKKERLNALRDYNMLRSESEASFDECKMRRAHLRALGM
Aedes          LLFQQKVPLEKSLSMPDMQNKVKKRERMMVLREY..MRSESERSFDEVQIRKARLQSLGL
Anopheles      LIFQQKVPLEKSLSMPDMQNKFKKRERMMVLREY..MRSESERSFDEVQIRKARLQSLGL
Culex          KLLLQKVPLEKSLSMPDMQNKVKKRERMMVLREY..MRSESERSFDEVQIRKARLQSLGL
Leptinotarsa   LLFTNKAPLEKSLSMPDMQTKAKKRERLLVLREY..MRSESEKSFNECQIRRARLQSLGL
Rhynchophorus  LLFSKRAPLEKSLSLPDVQNNVKKRERILALREY..MRSESEKSFDQCQMRKARMKSLGL
Periplaneta    LLMTKKSPLSKSLSMPDMETRVKKRERLMALREY..MRSESEKSFDECQVRRARMQSLGL
Blattella      LLMTRN........................LREY..MRSESEKSFDECQIRRARLQSLGL
```

```
                1000         1010         1020         1030         1040         1050
Drosophila     AYLSPQNKSLAQSFRYMRTKPTIIAFKTPSMEEREHQVAAGEANGASPASRAEQGQLSSR
Ceratitis      AYLSPQNKSLAQSFRYMRNKPTIIAFKTPSLEEAEPDFTSIARENGDTLRELEEG....K
Spodoptera     ASKVPQNKSLVHSFRYMRTKPTIIAFKTPSFEERRSNDSILTLARR..............
Plutella       AYKSPQNRQLAHSFRYMRTKPTIIAFKTPSMDNCER..DNMLTIARR.............
Aedes          AYLSPQNKSLAQSFRYMRNKPTIIAFKTPSLENCEPRDSTNSMVVSP.............
Anopheles      AYLSPQNKSLAQSFRYMRNKPTIIAFKTPSLENCEPRDSTNSIVVSP.............
Culex          AYLSPQNKSLAQSFRYMRNKPTIIAFKTPSLENCEPRDSTNSMVVSP.............
Leptinotarsa   AYLSPQNKSLAQSFRYMRNKPTIIAFKTPSLENCEQRDSSSSFGSVGRQ...........
Rhynchophorus  AYLSPQNKSLAQSFRYMRNKPTIIAFKTPSLENCEKKDSSTSFIST..............
Periplaneta    AYLSPQNKSLAQSFRYMRTKPTIIAFKTPSLENCD...ATVALLASS.............
Blattella      AYLSPQNKSLAQSFRYMRNKPTIIAFKTPSLENCE...ASMALLPS..............
```

```
                1060         1070         1080         1090         1100         1110
Drosophila     MDSADKLQLARLSLSAEGASKPLEDQTQTGSPSRKSILRRPTFLRSLSASINNRTGGGGG
Ceratitis      GNGVDKTISTVLDVPEESVSKPLEDSSSRTKVGR...FRRPTFLRSLSTSIKNR..GSHG
Spodoptera     ..RLSKSLSPDRDIECQPEKIVECENDGSEAT...............DDFEANYPIGKPL
Plutella       ..RLSKTISPDKDVESGRKSSLKAQEESTESTS..............DEAFVNYPVGKPL
Aedes          ......GVFTQKEAEEGK...IVHRAGP.........HKPGATDEDKLSRPVNYPVGKPL
Anopheles      ......GVFTQKDAEEGRTAGALPTSGPASSVVAVGSGAASAASSNPASRPVNYPVGKPL
Culex          ......GVFTAKEAEEGK...ILPRTVSPH.......HKPGATDEDKLQRPVNYPVGKPL
Leptinotarsa   MSSLSFTSSPERQAEEGRLIPLSREGQLTPNF...........LQLHQLQPMNYPVGKPL
Rhynchophorus  LGNTSDYYSPERRVEEGQVFQLQDMTAANP.................KLKPFNYPVGKPL
Periplaneta    .RNSGASQNPERAAEEGRAPEIVKIGARSDDVS......LEVVAGKSPISSVNYPMGKPL
Blattella      .....GGQSPERSAEEGRSPAIVKVGARSDDVS......LELVAAKSPITSMNYPVGKPL
```

```
                1120                      1130         1140         1150         1160
Drosophila     GSTGSSTTNSG.........GKVTLDAGVMEIFIDGPYGAPSSHIFGAQHAVLIGTGIGV
Ceratitis      SLPPQGNHNAG.........SKVTLDAGVLEIFIDGPYGAPSSHIFGAQHAVLIGTGIGV
Spodoptera     EIYLDG..............................PYGAASSHIFLAQHAALVAAGIGV
Plutella       EIYLDG..............................PYGAPSSHIFRAQHAALIAAGIGV
Aedes          EIYIDG..............................PYGAPSSHIFQAQHAILIATGIGV
Anopheles      EIYIDG..............................PYGAPSSHIFQAQHAILIATGIGV
Culex          EIYIDG..............................PYGAPSSHIFQAQHAILIATGIGV
Leptinotarsa   EVKRSARRENR.....VEDVIIRISDSAGTEIYLDGPYGAPSSHIFRAQHAVLIATGIGV
Rhynchophorus  ELYNSKCLDNTSRIDMVSDIKVKYSDNGGTEIFLDGPYGAPSSHIFRAQHAVLIATGIGV
Periplaneta    EPINEE............NVSESSAEREGMEIFIDGPYGAPSSHIFRAQHAVLIATGIGV
Blattella      EILED.......................................................
```

```
              1170      1180      1190      1200      1210      1220
Drosophila    TPFASILQSIMHRYWKARHSCPRCQFEWASEIPKSVMNLRKVDFFWINRDQRSFEWFVNL
Ceratitis     TPFASILQSIMHRYWKARHTCPRCHFEWASDIPKTVMNLRKVDFFWINRDQRSFEWFVNL
Spodoptera    TPFASILQAVMYRYWSARQDCPKCGHTWASDMPRHNMNLKKVDFFWINREQRSFEWFVSL
Plutella      TPFASILQCIMHRYWKMRNVCPKCNHAWASDDFRQSMRLKKVDFFWINREQRSFEWFVSL
Aedes         TPFASILQSIMHRYWKARHCCPRCNFEWSSEIPPTIMNLRKVDFFWINRDQRSFEWFVNL
Anopheles     TPFASILQSIMHRYWKARHCCPRCSYEWSSEIPPTIMNLRKVDFFWINRDQRSFEWFVNL
Culex         TPFASILQSIMHRYWKARHCCPRCNFEWSSEIPPTIMNLRKVDFFWINRDQQSFEWFVNL
Leptinotarsa  TPFASILQSIMHRYWKARHTCPKCTYSWASDIPATVMNLRKVDFFWINRDQRSFEWFVNL
Rhynchophorus TPFASILQSIMHKYWKARHTCPKCKYAWASDIPASIMNLRKVDFFWINRNQRSFEWFVNL
Periplaneta   TPFASILQSIMLRYWKARHLCPKCKYSWASDIPATVMNLRKVDFFWINRDQRSFEWFVNL
Blattella     .PINEIIR...........ICD......................................

              1230      1240      1250      1260      1270      1280
Drosophila    LSQLEIEQAELGGAMERFLDMHMYITSSLQRTDMKAVGLQLALDLLHEKGKRDLITGLKT
Ceratitis     LSQLEIEQAELGGPMERFLDMHMYITSALQRTDMKAVGLQLALDLLHEKGKRDLITGLKT
Spodoptera    LSQLEIEQAELGGAMERFLEMHMYITSALQKSDMKAVGLQLALDLLHEKEKRDLITGLKT
Plutella      LSQLEIEQAEA..TAERFLEMHMYITSALQRTDMKAVGLQLALDLLHEKSKRDLITGLKT
Aedes         LSQLEIEQAELGGAMERFLEMHMYITSALQKTDMKAVGLQLALDLLHQKEKRDLITGLKT
Anopheles     LSQLEIEQAELGSAMERFLEMHMYITSALQKTDMKAVGLQLALDLLHEKEKRDLITGLKT
Culex         LSQLEIEQAELGGAMERFLEMHMYITSALQKTDMKAVGLQLALDLLHQKEKRDLITGLKT
Leptinotarsa  LSQLEIEQAELGGAMERFLEMHMYITSALQKTDMKAVGLQLALDLLHEKEKRDLITGLKT
Rhynchophorus LSQLEIEQAELGGAMERFLEMHMYITSALQKTDMKAVGLQLALDLLHEKEKRDLITGLKT
Periplaneta   LSELEIEQAELGGAMERFLEMHMYITSALQKTDMKAVGLQLALDLLHKKEKRDLITGLKT
Blattella     ......SGSELKGDDD.........VSE................................

              1290      1300      1310      1320      1330      1340
Drosophila    RTNAGRPNWDKVFKQLQAQQKGKVTVFYCGPPQLAKTLRYKCDQYGFAFRKECF
Ceratitis     RTNAGRPNWDKVFKQLQAQKKGKVTVFYCGPPQLGKTLRYKCDQYGFAFRKECF
Spodoptera    RTNAGRPNWDKVFKQLQEQNKGKVTVFYCGPPQLAKILRVKCDQFGFGFRKEVF
Plutella      RTNAGRPNWDKVFQQLQQQHKGKVTVFYCGPPQLARILRVKCDQFGFHFSKEVF
Aedes         RTNAGRPNWDKVFKQIQDQKKGKVTVFYCGPPQLAKTLRYKCDQFGFQFRKEVF
Anopheles     RTNAGRPNWDKVFKQIQDQKKGKVTVFYCGPPQLAKTLRYKCDQFGFQFRKEVF
Culex         RTNAGRPNWDKVFKQIQDQKKGKVTVFYCGPPQLAKTLRYKCDQFGFQFRKEVF
Leptinotarsa  RTNAGRPNWDRVFKTISDQKKGKVTVFYCGPPQLARILRVKCGQFGFNFRKEVF
Rhynchophorus RTNAGRPNWDKVFKQISDQKKGKVTLFYCGPPQLGRILKVKCDQFGFIFRKESF
Periplaneta   RTNAGRPNWDKVFKQLQDQKKGKVTVFYCGPPQLGRILRVKCDQFNFIFRKEVF
Blattella     ......................................................
```
