## Supplementary Alignment Files for "Assessing target genes for homing suppression gene drive": oct.pdf

```
                  1                     10                    20        30
Drosophila    ....MLLCDGLG..............PEPPR...........QRHRNRTSAARIRKRPKC
Ceratitis     MSTNILATDMLPVYIMQHNITNAIRLPETATTTKTTIDIVSAKLNRTTSASAKRRKRRNV
Spodoptera    ............................................................
Plutella      ............................................................
Aedes         .....MMNPSND...............DLPKYMYSAAAAAVSSSFSLASSSGSAHSQEGH
Anopheles     .....MMNPSND...............DTTNYMVS.............SSSSSSHLQSAA
Culex         .....MMNPSND...............DLPKYMYSAAASSFS...SLSSSSGSSYGPQAQ
Leptinotarsa  ............................................................
Rhynchophorus ............................................................
Periplaneta   ............................................................
Blattella     .....MLTTSVD...............GIT.......................SSQRQGV

                     40             50                60
Drosophila    CCGDGGSGNQAE.....QPGGIVSNPIS.......YGQSLTT..................
Ceratitis     CVEPDEQRTTCDRAVAATNATTITTPIATANKSAYYADSAATSNIKDFLHQCTIPPPPPP
Spodoptera    ............................................................
Plutella      ............................................................
Aedes         SSFGGGSNSTGG................................................
Anopheles     ALLASTFGLADG................................................
Culex         T...GSGISHGG................................................
Leptinotarsa  ............................................................
Rhynchophorus ............................................................
Periplaneta   ............................................................
Blattella     CVKGKSSDDQER................................................

Drosophila    ............................................................
Ceratitis     SPAAAAATATATLSVMSRTATTTPPPPRLMVATSATDTTCRLLHQHHHHHQQQQQQQHPS
Spodoptera    ............................................................
Plutella      ............................................................
Aedes         ............................................................
Anopheles     ............................................................
Culex         ............................................................
Leptinotarsa  ............................................................
Rhynchophorus ............................................................
Periplaneta   ............................................................
Blattella     ............................................................

                                        70                80        90
Drosophila    ...................LARVTAAALTT.......AAMLHTTNALAATGSSSASNSST
Ceratitis     YRRRATIATTETTTHPPPSLSRARASSLGFNLSTAFAAALLHTSVAVAAAASTHAQSHHG
Spodoptera    .....................................................MDRENLT
Plutella      .....................................................MDLP...
Aedes         ...................TSLFFNGSG..........AALASAAAKAAGSSTIFTVGAS
Anopheles     ...................TVLYNHSVR..........DRGDAGALAAASAASVAGTGGL
Culex         ...................LGLFFNGSGSSAGSSSSSSAALASAAAKAATSATILGSG..
Leptinotarsa  .....................................................MGN....
Rhynchophorus .....................................................MDSG...
Periplaneta   .....................................................MASNPDI
Blattella     ...................QRRMFSAFG................TLSCKIYYGMASNSDI

                  100                110       120       130       140
Drosophila    GG.IALP.........LGTATPATHELNAT.QPFGGSGLNFNESGAGLSDHHHHQQHNPD
Ceratitis     GGGIAAKHLTPQSQAVLDNGVLSQLDSNATSTLLGASGITLPPSNATLLDDGGAGEDGVA
Spodoptera    HH................................AALNATVNATGNSTVAEAEDD.....
Plutella      .....................................NGTANATG......GHDV.....
Aedes         AV............PGRAGDPGGPGDPTGNHVIFQGYSTSSPTSLD.MTTVGSDAVMDDC
Anopheles     ES............ATAGGDTLSVFDASGSSIMLQGFSSTAPIGGTIETVGANESIVDG.
Culex         ..............PGRGGGQDDS.AAGGSHMIFQGYSSTGAASLDPLDLATTSSVLDEG
Leptinotarsa  .................................DTNGSLSHNNTSPLPET..........
Rhynchophorus ...............................KFQSNISMSKNVSAAAYDDSTS.......
Periplaneta   KI............ADPSSGQNIRYNSTSSTSSFVHTTTPASVGDALLDNLTSTVTLIVE
Blattella     SI............EDPSNGQQHHYNSTPPGSLFVQTSTPSSVADAIVDNITSTIATTIS

                           150       160       170       180       190
Drosophila    .........EDWLDNIVWVFKAFVMLLIIIAAICGNLLVIISVMRVRKLRVITNYFVVSL
Ceratitis     GVDGAALNGSEWFDVLFLVFKASLMLFIIIAAIFGNLLVIISVMRVRKLRIITNYFVVSL
Spodoptera    ..........DLTQSIFFKLRMSVLLLIVIMAVLGNMLVIVSVMRHRKLRVITNYFVVSL
Plutella      ..........DLSYAVMFQLRTAILLLIVIMAVLGNLLVIVSVMRHRKLRVITNYFVVSL
Aedes         C..GAAAASGEWLDVLLLVLKASIMMFIIVAAIFGNLLVIISVKRHRKLRVITNYFVVSL
Anopheles     ...GIDAGGGEWIDVVLLVLKASIMMFIIVAAIFGNLLVIISVKRHRKLRVITNYFVVSL
Culex         SGIGAGADSAEVLDGVLLVLKASIMMFIIVAAIFGNLLVIISVKRHRKLRVITNYFVVSL
Leptinotarsa  ..........EWTDDFVIFVKATIMGSIIIASIFGNLLVIVSVMRHRKLRIITNYYVISL
Rhynchophorus .........EEIWNNFAHFLKATIMGSIIAASILGNLLVIVSVMRHRRLRVITNYYVVSL
Periplaneta   EE.......QEWGDILTLILKAGAMGFIILAAIFGNLLVIVSVMRHRKLRIITNYFVVSL
Blattella     D........KEWTNVLSLILKGSAMGFIIIAAIFGNLLVIVSVMRHRKLRIITNYFVVSL
```

```
                 200       210       220       230       240       250
Drosophila       AMADIMVAIMAMTFNFSVQVTGRWNFSPFLCDLWNSLDVYFSTASILHLCCISVDRYYAI
Ceratitis        AMADIMVAMMAMTFNFSVQVTGRWNFGIFVCDLWNSLDVYFSTASILHLCCISVDRYYAI
Spodoptera       AFADILVAMVVMPFNFSVQFYNSWLFGPTICDLWNSSDVYFTSTSILHLCCISVDRYYAI
Plutella         AFADILVAVAVMPFNFSVQFYHEWIFGQVICDLWNSSDVYFTSTSILHLCCISVDRYYAI
Aedes            AMADIMVAMMAMTFNFSVQITGSWKFGSFMCDVWNSLDVYFSTASILHLCCISVDRYYAI
Anopheles        AMADIMVAMMAMTFNFSVQITGSWKFGSFMCDVWNSLDVYFSTASILHLCCISVDRYYAI
Culex            AMADIMVAMMAMTFNFSVQITGSWKFGSFMCDVWNSLDVYFSTASILHLCCISVDRYYAI
Leptinotarsa     ALADMLVAMFAMTFNASVQIYDEWKFGYFMCDVWNSLDVYFSTSSILHLCCISVDRYYAI
Rhynchophorus    ALADMLVAMFAMSFNFSVEVFGRWLFSYTLCDVWNSLDVYFSTVSILHLCDISIDRYYAI
Periplaneta      ALADMLVAMVAMTFNASVQLTGRWMFGYFMCDVWNSLDVYFSTASILHLCCISVDRYYAI
Blattella        AFADMLVAMVAMTFNASVQLTGRWVFGYFMCDVWNSLDVYFSTASILHLMCISVDRYYAI

                 260       270       280       290       300       310
Drosophila       VKPLKYPISMTKRVVGIMLLNTWISPALLSFLPIFIGWYTTPQHQQFVIQNPTQCSFVVN
Ceratitis        VKPLKYPINMTKRVVFIMLLNTWVSPALLSFLPIFIGWYTTEEHKNYVKRHPDRCEFIVN
Spodoptera       VKPLKYPIKMTKKMAFVMLAATWLSPITISYVPIFMGWYTTSDHLKVQRAD..ECEFIVN
Plutella         VKPLKYPIKMTKKMAFVMLAATWLSPVTISYAPIFMGWYTTEEHLEFRKYNPRTCDFKVN
Aedes            VKPLKYPISMTKRVVAIMLLNTWISPALLSFVPIFAGWYTTDKHKEDVLNNPDSCLFIVN
Anopheles        VKPLKYPISMTKRVVAIMLLNTWISPALLSFVPIFAGWYTTEKHKQEVLDNPDSCLFIVN
Culex            VKPLKYPISMTKRVVAIMLLNTWISPALLSFVPIFAGWYTTDKHKEDVANNPDSCLFIVN
Leptinotarsa     VKPLKYTLTMTKKVVALMILNTWLSPAVLSFLPIFIGWYTTTENQEFRNNHPDICEFIVN
Rhynchophorus    CKPLKYSLHMTKKVVALMLINSWVSPAFLSFLPIFMGWYTTDAHQEFRKQHPDECIFVVN
Periplaneta      VKPLKYPINMTKRVVAFMLLNTWVSPAIIAFVPIFLGWYTTDENQLYRIKHPDECDFKVN
Blattella        VKPLKYPINMTKRVVAFMLVNTWVFPAIIAFVPIFLGWYTTDEHQAHRIDHPDECNFVVN

                 320       330       340       350       360
Drosophila       KYYAVISSSISFWIPCTIMIFTYLAIFREANRQEKQLMMRHGNAMLMHRPSMQP......
Ceratitis        KPYCVISSSISFWIPCTIMIFTYLAIFREANRQEKQMMARQGNAMLMHRHSEDAR.....
Spodoptera       KPYAVISSSISFWIPCTIMLFTYYAIFREANRQEKNLHARAGNAMLMHRH..........
Plutella         KPYAVISSSISFWIPCTIMIFTYLAIFKEANRQEKALHARAGNAMLMHRH..........
Aedes            KPYAVISSSISFFIPCTIMVFTYFQIFREANRQEKQLAMRQGTAMLMHQHNAGGGGVGGT
Anopheles        KPYAVISSSISFFIPCTIMVFTYFQIFREANRQEKQLALRQGTAMLMHQHNTGGG.VGGG
Culex            KPYAVISSSISFFIPCTIMVFTYFQIFREANRQEKALAVRQGNAMLMHQHNTGGG..GGE
Leptinotarsa     KPYAVISSSISFWIPCTIMIFMYLSIFREANKQEKEMYNRHGAAKLLHQN..........
Rhynchophorus    KAYAVISSSISFWIPGSIMIYMYLQIFKEANRQEKEMYNRHGAALLLHAN..........
Periplaneta      KVYVIVSSSISFWIPCTIMVFTYLAIFKEANRQEKQLHSRIGNAMLMNHHRNSRE.....
Blattella        KAYVIVSSSISFWIPCTIMVFTYLAIFREANRQEKQLHSRIGNAMLMNHHRNSRD.....

                       370       380       390       400       410
Drosophila       ..........SGEALSGSG.SSKTLTLHEVEQ...EHTPTKDKHLIKMKREHKAARTLGI
Ceratitis        ..AVSGVSNNTGEALSGSG.SSKTLTLHEVEQ...DHTPTKDKHLIKMKREHKAARTLGI
Spodoptera       ..........SREVGDKNG......ALHINAT.....TPTKDRNILKMKREHKAARTLGI
Plutella         ..........SREVNDKNG......ALHIN.......TPTKDKNILRMKREHKAARTLGI
Aedes            GSGSAGGGRTNGEALSGSG.SSRTLTMHEVDA...EQTPTKDRHLIKMKREHKAARTLGI
Anopheles        G....SGGGTNGEALSGSG.SSKTLTLHEVDA...EQTPTKDRHLIKMKREHKAARTLGI
Culex            G...GGGGGTNGEALSGSG.SSKTLTMHEVDA...EQTPTKDRHLIKMKREHKAARTLGI
Leptinotarsa     ........NTNGDMLSNSGGSSKTLTQHEINQ.DLHHTPSKERNISKMKREHKAARTLGI
Rhynchophorus    ........NTNGDMLSNSGGSSKTLTLHEINQ.DLHHTPTRERSISRMKREHKAARTLGI
Periplaneta      ......YN.SNGDVLSSSGGSSKTLTMHEVHHGDLHSTPTKDRNIIKMKREHKAARTLGI
Blattella        ......YNNSNGDVLSSSGGSSKTLTLHEVQHSDLHSTPTKDRNIIKMKREHKAARTLGI

                 420       430       440       450       460       470
Drosophila       IMGTFILCWLPFFLWYTLSMTCEECQ..VPDIVVSILFWIGYFNSTLNPLIYAYFNRDFR
Ceratitis        IMGTFILCWLPFFLWYTITSLCDLT...APDIVVAILFWIGYFNSTLNPLIYAYFNREFR
Spodoptera       IMGAFILCWLPFFLFYVSTSLC.DSCR.CPEVVTVIMFWTGYFNSALNPIIYAYFNRDFR
Plutella         IMGAFILCWLPFFLFYLSTSLCGDGCD.PPETVTVIMFWTGYFNSALNPIIYAYFNRDFR
Aedes            IMGTFILCWLPFFLWYIITSLCDEC..PNPDIVVVLVFWVGYFNSTLNPLIYAYFNRDFR
Anopheles        IMGTFILCWLPFFLWYIITSLCDRC..PNPDIVVALVFWIGYFNSTLNPLIYAYFNRDFR
Culex            IMGTFILCWLPFFLWYIITSLCDEC..PNPDIVVVLVFWVGYFNSTLNPLIYAYFNRDFR
Leptinotarsa     IMGTFILCWLPFFTWYDTVYLCDSCREHCPKEVVAVVFWIGYFNSTLNPLIYAYFNRDFR
Rhynchophorus    IMGIFILCWLPFFLWYDSVSLCTTC..ECPHLVVAILFWIGYFNSTLNPLIYAYFNKDFR
Periplaneta      IMGTFILCWLPFFLWYVSTTLCG.EACPCPDIVVAVVFWIGYFNSTLNPLIYAYFNRDFR
Blattella        IMGTFILCWLPFFLWYVSTTICGEELCPCPDIVVALVFWIGYFNSTLNPLIYAYFNRDFR

                   480       490       500       510       520       530
Drosophila       EAFRNTILCLFCNWWKDRHLPLD.IDIRRSSLRYDQRAKSVYSESYLNSTTPSHRRQSQM
Ceratitis        EAFRNTIQCLFCNWWRDRGMPLD.IDIRRSSLRYDTRAKSVYSENYLSASQP.PRRTSQL
Spodoptera       NAFKNTLACAFCSFCKRSASDLD......ALERLDRRGSAQLRVPVPS......RRTSDL
Plutella         NAFKNTLACAFCSFCRRSATDLD......ALDRLDRRGSGQLRVPMAS......RRASDL
Aedes            EAFRNTLDCMFCAWWRRETSPLD.INVRRSSLRYDCRARSVYSENYLRSTTQNDRLNSEI
Anopheles        EAFRNTLDCMFCAWWRRETSPLD.INVRRASLRYDCRAKSVYSESYLRPSSQTDRFHSEI
Culex            EAFRNTLDCMFCAWWRRETSPLD.INVRRSSLRYDCRAKSVYSESYLR.PSQNDRFNSEV
Leptinotarsa     EAFKNTLQCAFCSLCKRSPSNLDFIDNRRPSIRYEDRTRSVYSETYLK..HVDRR.CSEY
Rhynchophorus    EAFKNTLQCAFCSLCRRPPSDLNYVDNRRPSIRYEDRTRSVYSETYLK..HVDRR.CSDF
Periplaneta      EAFKNTLQCAFCSLCRRPPSDLEALDVRRHSLRYDERTRSIYSETYLK..HIDRRRSSEF
Blattella        EAFKNTLQCAFCSLCRRPPSDLDALDVRRHSLRYDERTRSIYSETYLK..HIDRRRSSEF
```

```
                     540       550                     560           570
Drosophila     VDNLXY.RDEALLTPIAQQQQR......LAAGGS..........RLGG....QLAAAAKD
Ceratitis      VDSLXYSRDDSLLSVTRTTPQRSTSMQLLPTQQSETTSTLRTATGVGGSGMHQLAPLPLP
Spodoptera     ASL.........................................................
Plutella       TSL.........................................................
Aedes          GESL........................................................
Anopheles      GESL........................................................
Culex          GESL........................................................
Leptinotarsa   GSSL........................................................
Rhynchophorus  GSTL........................................................
Periplaneta    GSSL........................................................
Blattella      GSSL........................................................

                     580       590                    600       610
Drosophila     GRESKDAKDTGKDAGKGKSNVDCLAG...............DDVQEIQIEIPMEYINKWN
Ceratitis      SYVGKRLLDIKPSAGQELNGAELKGGRGGGGDGRHANVGDDDEPQQIQIEIPLAYVKKWN
Spodoptera     ............................................................
Plutella       ............................................................
Aedes          ............................................................
Anopheles      ............................................................
Culex          ............................................................
Leptinotarsa   ............................................................
Rhynchophorus  ............................................................
Periplaneta    ............................................................
Blattella      ............................................................

                620       630
Drosophila     KNNNAAASTASSHV..............................................
Ceratitis      KSNKNTATTATATGTTTTTGCTTAISTTAATTTATGAALVNSTATVTTAATTTTSKAASP
Spodoptera     ............................................................
Plutella       ............................................................
Aedes          ............................................................
Anopheles      ............................................................
Culex          ............................................................
Leptinotarsa   ............................................................
Rhynchophorus  ............................................................
Periplaneta    ............................................................
Blattella      ............................................................

Drosophila     .........................
Ceratitis      TATTTITTTVTTPKGTEHTHAHTHV
Spodoptera     .........................
Plutella       .........................
Aedes          .........................
Anopheles      .........................
Culex          .........................
Leptinotarsa   .........................
Rhynchophorus  .........................
Periplaneta    .........................
Blattella      .........................
```
