## Supplementary Alignment Files for "Assessing target genes for homing suppression gene drive": stl.pdf

```
              1                                         10        20
Drosophila    MTVAAIG..................................SNRHRRLALIVLTTLWL..
Ceratitis     MMCSPSGFVFVLIAILIWNCLHLGCAVHDSSSNSGSNNNSNSNNNNNNKSSVSNRFWFSN
Spodoptera    ........................................MVMYWRKVILAAALSSILD.
Plutella      ..........................................MHWKEVFVVASLTVLLD.
Aedes         .......................................MLLRYRWLPQRLVRRRQFCP.
Anopheles     ............................................MPRPQLQPFFKAVG..
Culex         ............................................KCSPSLRFWRRR....
Leptinotarsa  ............................................MRGVWWRFLFTAY...
Rhynchophorus ...........................................MHEKLLLALLACAVVR.
Periplaneta   ..........................................MVSRLLWICFSLGAQLCS
Blattella     ........................................MIVEARALGRRFELQLRPSP

                                                            30        40
Drosophila    ..........................................LIGASSSSSSTSSTNPKS
Ceratitis     KNTANHNRLLPEPGGRLAVARNVNDRKSGNISNSNGNDRSGDKIGSTHNSLSVSNTKTKS
Spodoptera    ............................................................
Plutella      ............................................................
Aedes         .......................................TWDGFSVKMLCIVVFLVAIRA
Anopheles     ..............................................SVMLQIIALLALIL
Culex         ............................................................
Leptinotarsa  ............................................................
Rhynchophorus ............................................................
Periplaneta   A...........................................................
Blattella     AL..........................................................

                              50       60         70        80
Drosophila    VN..........IELRRKRSLHYNGIELDKNTLVGHLREHERALIFGSRS...PEEAPEF
Ceratitis     SNNDAVRGRRLRIDITRKHSVHLNGVEINEDSLANHLAEHEKELIFG..K...AKDGRPL
Spodoptera    .................VSSEQ.FGNDDHSKQLLEHLTESEKIFLFG.....ANPSEAQF
Plutella      .................ATSEQLFGDDEHSNRVLEHLTDAEKLFLFG.....AIDKTPQF
Aedes         VDHRDVGVNCFHLDVQKVNHIVLDNEKVDKSNLHRYLREHEHSFLFGKE......STNDF
Anopheles     IAARNPTVRCLHMEVERVNHLVLDNAKIDERNLHNFLEEHEQLLLFGDR......TTDDF
Culex         ...............RRQSLARNTSGEIDEKNLHQFLKEHEKLLLFGEND...AASAEDF
Leptinotarsa  .................FLIELAVSQQINKHQLRELLTLPEWTSLVGNSHSVASEGGDDS
Rhynchophorus ........................PVVALKYALHDAMTGPDLAYYFHTDD...ENSVPDY
Periplaneta   .................YTSTLHQEVTETN..LRDLLTTAEWTSLLG.....ASSDAPEF
Blattella     ....LDPQFVLLSRWRNHTEMLPSHLAEQNCFFRSDATHRAAVALCGGMRGIITSPENEY

              90     100       110       120       130           140
Drosophila    KLVHLAQQRPEEQQRLEERKRPGEAEQHGGNPHQRRQRRSLPSED....GNGPLPPEEEE
Ceratitis     KLVHLQHLTDANDDNGVSNAAVG.STAHKSSHRVRRSDTSATADDQSDGGSGGAQPQVAS
Spodoptera    EVAHLKQ..........................PKRRKRSLLYD............DQDL
Plutella      EIAHLKH..........................PRRRKRSIYNRQ..........PNQDL
Aedes         SIVSVR............................RRKRTTRST........EKLIEFNIS
Anopheles     KIVNVR............................QRKRESRSTPG......SGFLQFNLS
Culex         TIVSVR............................RRKRTTRST........ERLISFNVS
Leptinotarsa  EDNDQEL..........................SSASRVIIEVP................
Rhynchophorus DIVFLPED.........................WPQQERFLDNEP.......GNELNYSF
Periplaneta   AVVLPVR..........................RRRRRR.WLLP................
Blattella     YVVNPLP..........................RRFRRRSTDVP................

                150               160       170       180       190
Drosophila    QEAGEDVEAR.........LQLQQSPQLIDDAFIFIRRTENGTQFVEHSPQLLKRLERCF
Ceratitis     QSEEEVVSVQIKDHEQSVHLQLRQTPKLIDDAFVFIRRYANTSQFIEDSHKLAERYERCF
Spodoptera    VGLKKHLDIK...........LSPRNSFLDPQFLLMRRWSNNTELVLDQ.HADEEVDSCY
Plutella      ELIKKILDLK...........LSPRNTFLDPQFLVVRRWANNTEPVVD..HSDDEVDSCY
Aedes         DSTDESFLIK...........LRKSNILIDDSFALIESYDNSTQLLDDSYQLVQKFADCF
Anopheles     DGREDNFALR...........LQKSDILIDESFAFIESYDNSTQLLDDSYDRVLKYRDCF
Culex         NSAVADFSVE...........LRKSNILIDDSFAFIESYDNSTQLLDDSYQLVQQYADCF
Leptinotarsa  .ELGHSFDLD...........VKSSPDILTPDFVVLETFANRSQPITV...QKE.DRNCY
Rhynchophorus SAFERQIDLK...........LKRNENIVAPNFKTYIHEHDGNVRYLN.....EIPANCH
Periplaneta   .DETG.AIPV...........SLDAR.....QLGRRFDLRLVPSPALL...DPSFVLLSR
Blattella     .HVIVRWTPP...........AFAARNAETSHISHNYQTNELSEATLS...QSSQIPTDY

               200          210               220        230          240
Drosophila    YRSPA....AALDLCEPG........NVRGVFQQNGSD.LVIHPLPARFGTGT...HVLY
Ceratitis     YSNEN....SALDLCDDE........NVRGVFHHNSTN.YIIHPLPARFGNST...HVIL
Spodoptera    YRGDN....AALY..........VCDDVRGLVRS.NDTYYFIHPLPDRFHDDHAKAHVIV
Plutella      YRSQN....AALY..........VCNDVRGVIKADNDSHYFIHPLPERFHNDDGKHHVIV
Aedes         YRNDY....AAFDLCEG.........EARGLIKLNNSD.LILHPLPERFGANN...HVLI
Anopheles     YRNER....AAFDLCEG.........SVRGLIRTNESE.IVIHPLPERFGTGA...HVIL
Culex         YRNDH....AAFDLCEG.........GIRGLLKSNQSD.LIVHPLPERFGSSS...HVLI
Leptinotarsa  FRGTT....AALS..........LCKGVRGIVHLNDTDFFLIHPLPSRFHNETANPHILI
Rhynchophorus YLHREGDLVAALSICNPK........SVQGLIFFDNST.FEIHPLTPNLQRLAKREARTG
Periplaneta   WRNRT....DMLS.................PRLAEPHCFYRGDDGPHRAAVALCGGDMRG
Blattella     TQGNT....AMASSINDEVNRTLLEQSRNTIRLTADTKSFNNGNVPERNVVHDVQNSSSS
```

```
                      250          260      270                 280
Drosophila    QARVDKS.GGGYS...EASSRRTASPLDSQLQFEPDAEEFNEP..........RRRLR..
Ceratitis     KSNYDLPPKKVLD...TAVMPLAATALNTSSYFEPEEEEFNELDDLNIESTLSEDRLRPA
Spodoptera    KKQMNENKDDDDS.....CIERDNIID.DPQPANTWKKHRRKREVPSNVN....NNLEG.
Plutella      KRSPTDRNNNDQSNQENTCVESTDLDKFNNNPSQIWTKHRRKREARQSTIESERNSLEK.
Aedes         NKNRTTNLTTAHV...NQT........DKNVIFEP...DMSH..................
Anopheles     NRNKT....IPSD...ESG........ETKVMFEP...DMNEP.................
Culex         YRNRTNDSAAPST...NQT........DENVIFEP...DMAA..................
Leptinotarsa  QKKHHAVHTGNCP..HYDTIQLKQN.TTKDITYAKVNNKIHETHLEWSSDEEEDIDKDIN
Rhynchophorus VPHIIKRAKFASEYLLDDLFPTAGNIWNKNPLYNDNIDDFFN..................
Periplaneta   ......IIASPDD..EYYVINP......LPRRFRRRSADVPHVLVRWTP......PAQR.
Blattella     NKKDRLIHYSIDK..ELVTYETNSINAYVSPSVTNQSESVSETSKNSESGRTIDIPAAS.

                                            290       300
Drosophila    ...........................AQSQLQSPLRLRFQPRHHHP.............
Ceratitis     RNSGHEANEDGEPEEIHNYTYHHKGSGDNSRKYLELNLRYKPRYHHNSAQKPQLHEQQLQ
Spodoptera    ...........................FKVNLAHRNKTITDFIQKLS.............
Plutella      ...........................LKLDLAHRNRTISTFINKLD.............
Aedes         ...............................YEPTY..FSMPSKDRS.............
Anopheles     .............................TRRRPTVPFYGQSQQSRL.............
Culex         ...............................YEPT...FNEPRDRRN.............
Leptinotarsa  ...........................ISSRSKRDVLEDYLDDFSYS.............
Rhynchophorus ............................EDLIQNTVQDNILHRKVQS.............
Periplaneta   ...........................LDQALRPR........AAKN.............
Blattella     ...........................SKTNERTSEGSHTCETCSET.............

                    310
Drosophila    ...HQHHQQR..................RRR............................Y
Ceratitis     HLKYQHHQQHNQHLQQQINKQKSNKVPQRRRKQKEHHHKHQKQQEIHDGATFTHEHSDVF
Spodoptera    ..NGAPRQRRA.................................................
Plutella      ..SIQNRQRRA.................................................
Aedes         ....KRHIG...................................................
Anopheles     ....KRHISSN.................................................
Culex         ....KRHIS...................................................
Leptinotarsa  ..NLGRKKRE..................................................
Rhynchophorus ............................................................
Periplaneta   ..PAARRRRSP.................................................
Blattella     ..ILKRSKRTT.................................................

                320                                       330       340
Drosophila    IGDAPRSR...............................WSDIPEELFIETAIFVDSDLY
Ceratitis     VHNKKKNKETSAESENNAEHDDITKQRHRRRRRRYIASGNSKIPQDLFIETAIFVDRDLF
Spodoptera    .......................................ILP...AVFVETAVFVDRDLY
Plutella      .......................................VLP...AIFVETAVFVDRDLY
Aedes         .......................................NIKVPDTLHVETAIFIDKDLY
Anopheles     ......................................HNAKVPDVLHIETAIFIDKDLY
Culex         .......................................NIKVPDTLHVETAIFIDKDLY
Leptinotarsa  .......................................SFPDGAPAFVETAVFVDKDLF
Rhynchophorus ..............................................MTAEIALFFDEAAY
Periplaneta   .......................................ALPPGAPVHVETAVFIDEDLF
Blattella     .......................................MLLPGAPIHVETAVFVDKDLY

                 350       360       370       380       390       400
Drosophila    AHMQKNFPTNTESKVVSFLLAMINGVQLLYHHPTLGRRINFVLKRLEIWKSWDPPGLVRS
Ceratitis     KHMSKNFPDNTESQMIRFVLAMINAVQLLYHDRSLGRRINFVLKRLEILYN.DPPSLRRS
Spodoptera    KHMTINFPKDTERELVRFVLAMINAVQLIYHDPSLGRPVNFILKRLEILHE.DPANLKRP
Plutella      KHMTVNFPKDTERELIRFVLAMINAVQLLYHDSSLGRPVNFILKRLEILHE.DPANLKRP
Aedes         QHMSKNFPKNTEAHLIRFVLAMVNGVQLLYNHPSLGHPINFILKRLEILHN.DPKDLRRS
Anopheles     RHMSKNYPKNTEAHLIRFVLAMINGVQLLYNHPSLGHPINFILKRLEILHN.DPKDLRRS
Culex         RHMSKNFPKNTEAHLIRFVLAMINGVQLLYNHPSLGRPINFILKRLEILHN.DPKDLRRS
Leptinotarsa  EHMKTNYPVDTERELIRFVLAMINAVQLLYHDPSLGRPVNFVLKRLEILKE.EVAGLLRP
Rhynchophorus KIFGP.FYGYDDKKLQTMLLAYINGVQALYHHPSLGTGLELVLVRLDIMKTQPREMPHYN
Periplaneta   QHMALNFPQDTERELVRVVLAMINAVQLLYHDPSLGRQVNFVLKRLEILHV.DPTSLQRP
Blattella     RHMALNFPTDTERELVR........VQLLYHDPSLGRQVNFVLKRLEILHV.DPANLQRP

                 410       420       430       440        450       460
Drosophila    RDVENYLNSFCKWQEKLNPFSDADPLHYDHALVLTGLDLVTYDKG.KANSQVVGMATVKG
Ceratitis     DDIDTYLSNFCEWQQRLNPPLDTDTMHYDHAVILTGLDLYVVDKGGKKISQVVGLAPVAG
Spodoptera    HDIDRFLSNFCTWQRLENPPGDNDPLHWDHALILTGLDLYVVNKNGKVSSQVVGLAPVAG
Plutella      HDIDRFLSNFCTWQRLENPPGDTDPLHWDHALILTGLDLYVVSKNGKVSSQVVGLAPVAG
Aedes         SDIDVYLNSFCGWQRKLNPISDADPVHFDHAVILTGLDLYVVSKTGKVSNQVVGLAPVAG
Anopheles     SDIDIYLNSFCGWQRKMNPISDADPVHFDHAVILTGLDLFVVSKNGKVSNQVVGLAPVAG
Culex         SDIDVYLNSFCGWQRKLNPISDADPVHFDHAVILTGLDLYVVSKTGKVSNQVVGLAPVAG
Leptinotarsa  PDIDRFLSNFCNWQKTKNPPGDSQPLHWDHALILTGLDLYVRGKHGKISNQVVGLAPVAG
Rhynchophorus GERSQLLDSFCEYQASINPRDDSNPDHWDMGLYVSGLDFFAYE.NGKKSGVTMGLATVGG
Periplaneta   HDIDRFLSSFCTWQRSENPPRDSDPLHWDHALILTGLDLYVVNKNGKVSSQVVGLAPVAG
Blattella     HDIDRFLS..............................................LAPVAG
```

```
                      470         480       490       500
Drosophila     MCTSIYSCTINEA............KHFESVFVVAHEIGHNLGMRHDAK..EISCDPTMH
Ceratitis      MCTNTSSCTINEG............KHFESVFVVAHEIGHNLGMRHDTVS.ERNCDPTRY
Spodoptera     MCTVTSSCTVNEG............RHFESVYVVAHEIGHNLGMRHDGPLADNGCDPSAY
Plutella       MCTVTSSCTVNEG............RHFESVYVVAHEIGHNLGMRHDGPLADNGCDPSAY
Aedes          MCTMTSSCTINEG............KHFESVFVVSHEIGHNLGMRHDTS..ENNCDPSQY
Anopheles      MCTITSSCTINEG............KHFESVFVVSHEIGHNLGMRHDTS..ENNCDPSLY
Culex          MCTMTSSCTINEG............KHFESVFVVSHEIGHNLGMRHDTS..ENNCDPSLY
Leptinotarsa   MCTTTSSCTVNEG............RHFESVYVVAHEIGHSLGMRHDGPLAENDCDPSSY
Rhynchophorus  VCLDKYACIIAEFGTTNVFGKPYPSAGFTSVYILAHEIGHNLGMHHDSS..GNTCPKDGY
Periplaneta    MCTATSSCTVNEG............RHFESVYVVAHELGHNLGMRHDGPTAGNECDSSSY
Blattella      MCTATSSCTVNEG............RHFESV..........LGMRHDGPLAENECDPGSY

               510       520       530       540       550       560
Drosophila     IMSPKLGS.GKVTWSKCSRTYLEDFLMDPQAECLFDRDSFAGPWDHTAGGRLPGERFNAN
Ceratitis      IMSPTLGS.GKTTWSDCSRDYLEKFLRKGLGSCLFDHGQFANNLDHSAEGILPGERFDAD
Spodoptera     IMSPTLGS.GKITWSQCSKNYLQKFLDTVQSRCLLDHGNSAGQLDHSAEGILPGERFDAD
Plutella       IMSPTLGS.GKITWSQCSRTYLQKFLDTIQSRCLLDHGNSAGQLDHSAEGILPGERFDAD
Aedes          IMSPTLGS.GKITWSTCSRNYLSSFLKTSQAVCLFDRGHFGPSLDHAAQGMLPGERFDAD
Anopheles      IMSPTLGS.GKITWSSCSRNYLNTFLKTSQATCLFDRGHYGSSLDHTGEGRLPGERFDAD
Culex          IMSPTLGS.GKITWSACSRNYLNTFLKTSQSVCLFDRGHVAPNLDHNVEGKLPGERFDAD
Leptinotarsa   IMSPTLGS.GKNTWSSCSKRYLQKFLDTPQSRCLLDHGSSAGKLDHSAEGAFPGERFDAN
Rhynchophorus  IMSPSRGTNGETQWSSCSADVMAKLG...WAKCLKDHAKPIRSRDHSRFLDVPGRLYTAK
Periplaneta    IMSPTLGS.GKITWSSCSRRYLQLFLETPQSQCLLDHSISGGQLDHNAEGTLPGERFNAD
Blattella      IMSPTLGS.GKITWSSCSRRYLQLFLETSQSQCLLDHSSSGGQLDHSAEGTLPGERFNAD

                570       580       590       600       610       620
Drosophila     QQCMLRFG.KNFMQASTQSKMEICRDLHCRQDGLP..WTSHPALEGTECGPNMWCRGGTC
Ceratitis      QQCMLKYG.HHSMRAKNQKQADICTDLHCQRERYT..WTSHPALEGTECGANMWCRSGTC
Spodoptera     QQCMLKYG.RGSRHSGAQSLEDICRDLHCQRERYT..WTSHPALEGTRCGEDMYCRGGEC
Plutella       QQCMLKYG.RGSRHSASQSLEDICRDLHCQRERHT..WTSHPALEGTKCGHDMYCRGGEC
Aedes          QQCMLKYG.KDSIRSQAQHVADICRDLHCQRDRYT..WTSHPALEGTACGTLMWCRSGAC
Anopheles      QQCLLKYG.KDSTRSKSQDLADICRDLHCQRDRYT..WTSHPALEGTNCGKLMWCRSGVC
Culex          QQCMLKYG.KDSIRSQSQNVADICRDLHCQRDRYT..WTSHPALEGTACGTLMWCRSGVC
Leptinotarsa   QQCMLKYG.RGSRHSSQQPLNDVCRDLHCERERYT..WTSHPALEGTTCGNNMWCRGGRC
Rhynchophorus  KQCEILLRDRDAVAAPNQQQSEICYNLQCKTPHRSGYYHAGPALDGTPCGSGKYCYGGHC
Periplaneta    QQCMLKYG.RGSIHATQQPLDDVCRDLHCQRDRYT..WTSHPALEGTSCGNTKWCRSGRC
Blattella      QQCMLKYG.RGSIHATQQPLDDVCRDLHCQRDRYT..WTSHPALEGTFCGNNKWCRSGRC

                   630       640       650       660       670       680
Drosophila     EARSSKQGSYSALKSWPPEHIPEATHEKIIRHEPNRIPPLLHDYNPMDNELPGSPSNWGE
Ceratitis      VIRPTYVEPYHTPK..LIASATKVGYDKANFLESSKMGHLLHEYN......YNKPVSWSE
Spodoptera     VEREGS...........................TVVRGAWGRW............SGWS.
Plutella       VERYGT...........................STPLGSWGRW............GGWS.
Aedes          VSKIPGLTIQSSGK....HITAFKTIDKKAFIEGLKFASLKQDTAR....TLNTIPTWDA
Anopheles      VSKIPGLTIQSTGK....HITAFKTIDKKTFLEGLKFASLRQDISR....TLNTIPTWDS
Culex          VSKIPGLTIQSSGK....HITAFKTIDKKTFIQGLKFASLKQDTAR....TLNSVTTWEA
Leptinotarsa   VVKGL............................ALTAAYASQQTAA...RLGQINGGWSQ
Rhynchophorus  STRQLP..........................KPVKVVP................GGWSN
Periplaneta    VARGLS...........................ALEAGYSPNQ.........RVDGGWSE
Blattella      VGRGLS...........................ALQAGYSPNQ.........RVDGGWS.

                   690       700       710       720          730      740
Drosophila     WSEPSACESGCLYGQSRRLLEGSTGLRTLNRSCLNFPSR.....CIGRDRRFVTCNTPQC
Ceratitis      WSDASDCASGCLYGPSRRLLEGSTGLRTYSRRCIDHRRR.....CLGRDHKYEACIAKDC
Spodoptera     .....ECASACLVDDT..HAPAAAGVAVATRRCNRARHEKN..HCVGHDKKYQSCHAQQC
Plutella       .....ECASSCLLDER..HSAAASGIAVATRRCNRFRHDSGSSYCSGYDKKYQTCNAQQC
Aedes          WGDPTECESGCLYGESGRLKEGSVGLRQYSRICLDKRNS.....CIGSNKKYETCIAKQC
Anopheles      WSEPTECVSGCLYGESGRLKEGSTGLRQFTRKCLDKRKN.....CSGFSRKYETCIAKQC
Culex          WGNPTECESGCLYGESGRLKEGSVGLRQYNRACRDKRNS.....CPGSNKKYETCIAKQC
Leptinotarsa   WKHFSECASGCLFGEEGRLRSGSTGIMVSERMCNNPRPQGAGTDCDGPSTRYSTCTAQQC
Rhynchophorus  WKEG.TCSSGCIKDSK........GIQMSTRQCNNPVPVNTDQGCEGSNRQFSFCSDNKL
Periplaneta    WTSFSDCASGCLYSEEGRLHAGSTGIMVSTRRCNNPRPENEGLPCRGSDRRFRTCFAEQC
Blattella      ...............................................SDRRFRTCFAEQC

                     750       760          770         780       790
Drosophila     HKVP.VQTINDFATQVCMQARKSDPELT..GEGQQ..LPS.TLDASCKIFCRTRTNGTKS
Ceratitis      YSLKIVMTIDEFAYDKCSRAQKSDAELT..GEGVQ..IIG.DIEESCKVYCRTKSNGTKT
Spodoptera     SNVP.RMTVREFADQICLRARDVDPDLI..GTGLQR.IDSDDVSSACAVWCDTRGGGYKS
Plutella       NNVP.RLTVREFADQICRRAREVDPDLI..GTGLQR.MDSDDVSSSCAVWCDTRGGGYKS
Aedes          YNIA.RTTILDFANQICDRAKAFDSDIR..GIGIQ..KVAEDPEEACKIFCETKSGDPKT
Anopheles      YNIG.RTTILEFSNQICGRAKEFDADII..GYGLQ..KVAEDPEDSCKVFCETKAGVPKT
Culex          YNIA.RTTILEFANQICGRAKEFDSDIL..GMGLQ..KVADDAEDACKVFCETKSGTPKT
Leptinotarsa   SNVP.RLTIREFADQICNRVKDVDKDLT..GLGMQK.ISSD.AEEACLVWCQKRNGSTKS
Rhynchophorus  CKSK.KEGPVDYASQKCKEFSTLLSELDPKGSGLQAPHEEARLWTSCAIFCKRKDSGVFY
Periplaneta    GNVQ.RTTIRDFADQICLRAREVDTELL..GTGLQR.ISSD.AEEACTVWCYKRNGGSKS
Blattella      GNVQ.RTTIRDFADQICMRAREVDTELL..GTGLQR.ISSD.EMVVP.....KVEDG...
```

```
                     800       810
Drosophila      RRWTFPDGTTCKAKQHNPEDIT......................................
Ceratitis       RRWTFPDGTTCRTTLYKSEDIS......................................
Spodoptera      RGWALPDGTACSTAAPMFCISG......................................
Plutella        RGWTLPDGARCSRHAPQYCIAGSCQVSVYTVRYKSRGWTLPDGARCSRHAPQYCIAGSCQ
Aedes           KSWIFPDGTTCRVRSSDFDDYF......................................
Anopheles       KSWIYPDGTACKVQNGDFDDAY......................................
Culex           KSWIFPDGTACRVQSADFDDYF......................................
Leptinotarsa    RGWTFPDGTACQIRRNRYGIAS......................................
Rhynchophorus   TPRIELNDLGVSPYFPDGTWCN......................................
Periplaneta     RGWNFPDGTACQSHRSRYGKSM......................................
Blattella       ..HFLMVLHAKVIAHVMGN.........................................

                                               820       830       840
Drosophila      ...........................YCISGRCERFSCDNSTSNFFKMDSSFCEAR...
Ceratitis       ...........................YCIAGRCEKFSCDNSTQNYYKLDPNFCQQRKAS
Spodoptera      ................................VCQKFTCSSNPDTEFSLKPSDCEW....
Plutella        VSVYTVRYKSRGWTLPDGARCSRHAPQYCIAGSCQKFSCSPHADSQFSLTPSECEW....
Aedes           ...........................YCISGRCQKFSCNGSLENFYRIDPMFCPES...
Anopheles       ...........................YCVSGRCEKFSCNNTSNNYYRIDGALCTEV...
Culex           ...........................YCIGGRCQKFSCNNSADNFHRIDSTFCPDS...
Leptinotarsa    ...........................YCIKGRCEEFICDSLEEAAFAQSPDLCP.....
Rhynchophorus   ......................................RDNTGMDYYCIQHHCLPEN...
Periplaneta     ...........................YCIAGRCEEFVCDNYDMAPFSTLPEMCP.....
Blattella       ...........................PCIVSVEDEFVCDNYDAATFSTELEQCPGL...

                              850       860       870
Drosophila      .............SVRPTRDSSDQESRQQTNRQRYAERQEGKPQKN..............
Ceratitis       QARAEESESRRYNSVHTTAVAGNNINYKNVNGHSSSNSNNNNNHNNNIPTYSIYAPTTYR
Spodoptera      ...................EALRTATP.PTTSTKTSGTWRR..AVGG.............
Plutella        ...................EALQLQTATPHTSTKSSGTWRR..AVGS.............
Aedes           ..............SISPDRASNSIQQENGRDYKSLNLVHDAHQTTR.............
Anopheles       ..............YQYTERSGNSIHQENVRDYKSLPLNKNAYPIDR.............
Culex           ..............FTSADRVSNSIQQENGRDYKSLNLLKEAQPTSR.............
Leptinotarsa    ...................VDKSDNE.....LSWRTGLRGREGAIR..............
Rhynchophorus   ..................FKFTKIQVPDLDDLPFSQNAMPNKYPIPS.............
Periplaneta     ...................VSSRLQV......SRNKGRRREVPELRR.............
Blattella       .................TRGGGRLQVRDSSLFSRNKGRRREVPEART.............

              880       890                900       910        920
Drosophila      .HYENEVAKRPSYGQG........NHINAPASPYKR..RNYHKPYP.PEIPRPPPPASAS
Ceratitis       SKYENEVAIRGFHGQEREVPYKRKSDIYQPWNKYSYPLRDSAQPIAQPQPPKPPPPVSAS
Spodoptera      ............................................................
Plutella        ............................................................
Aedes           .KLENVTQNSVFYNYP.......ERRRIPEANFYK...ADNYGRWRVRQFEPEKYDKLSQ
Anopheles       .RAESSTIRTALSNYT.......ARKPPPDTEGYSSQAENTFNRWRER..SSSELNRLSE
Culex           .KPENVTQNSVVYNFP.......ERKRFPEANFYK...ADAYSRWRARNFEPVSYDRNSQ
Leptinotarsa    ............................................................
Rhynchophorus   ............................................................
Periplaneta     ............................................................
Blattella       ............................................................

                930                    940       950       960
Drosophila      LIALDRSSSSE...........SEWVAHPGCHSNCMTDSKGVQGVTSRLTGV........
Ceratitis       LVAMDTAVLAQSATAAAAAPESAEWIVKSGCHSSCMAKSMGIQVVTSRRTGA........
Spodoptera      .......................QWRPASGCHYHCISPGAGVRLVTATSRDRPL......
Plutella        .......................QWRAVSGCHYHCVSGGSGVRLVRAAGKQS........
Aedes           LQSRKN.................QWDIKSGCHFSCMEKSKGIQIVASRMDIR........
Anopheles       LQGRKH.................NWEVKSGCHFSCMERGKGVQIVSSTAGTN........
Culex           LRLRKN.................QWEVKSGCHFNCMEKSKGIQIVASKMDIR........
Leptinotarsa    ........................WKSASGCHYNCVSPGSGIRLVMSKKHGR........
Rhynchophorus   ..........................KLKDYFSIGPNGRPLK..................
Periplaneta     .......................LWSPASGCHFNCLAPGRGLRLVVAKDIKCRLEDNCEG
Blattella       .......................LWSPASGCHFNCLAPGRGLRLVVAKDMKCRLEDNCEG

                  970           980       990      1000      1010      1020
Drosophila      ...ESIQLCSYRI....QPCERLQTAAEYAEQTCARYRQKVRGLSGHGAQISASIDEPDR
Ceratitis       ...NNIQLCTHTA....KPCERLLSATEHAEHTCARYKLKVRGLSGYGTQIAASVDEPDR
Spodoptera      ...TSIQLCSPDS.RPESGCTQRKSPYQYATMVCTKYKDRVRRLSGLGMQISPALEDPDR
Plutella        ...TSIQLCRPDTPSARLGCSQRKTPYQYATMVCTKYKERVRRLSGLGMQISPALEDPDR
Aedes           ...TSIQLCTPNT....IACDKVLTTYEYATRLCKRYQQKVRGLSGVGMQIAPSVEDPDR
Anopheles       ...TNIQLCTANT....IACDKVLSTYEFATQLCKRYQQKVSGLSGIGMQIAPSIEDPDR
Culex           ...TSIQLCTPNT....IACDKMQSTYEYATRLCKRYQQKVRGLSGTGMQIAASVEDPDR
Leptinotarsa    ...SSIQLCQPDK....FGCSRVKSPYQHASMVCTKYKERVRRLSGLGMQISPAVEDPDR
Rhynchophorus   ...TSIKISDIRP...PAEEDWETKDYVELPTIEQRFNALIQEMNDLDFF..........
Periplaneta     GRVTSIQLCEPDK....LGCGRLKTPFEHATTVCSKYRERVRRLSGFGMQISPVAQDPDR
Blattella       GSVTSIQLCEPDK....LGCGRLKTPFEHATTVCSKYRERVRRLSGFGMQISPVA.....
```

```
                         1030      1040      1050        1060      1070
Drosophila     SCRVGCQDEFIKYRYYLVNGRNGHFPPGTRCSPVG...KRYCVYGRCLEFGDDDLPLDKT
Ceratitis      SCRVGCQDEFIKYRFYLVNGRYGHFPLGTKCSHTE...ERYCVNGKCLEFGPDNIPLQQS
Spodoptera     PCRIACQDERVSHRFYLVNGHEGWFPLGTSCGKR...NASYCVSGKCLEFGPDNTPVSEM
Plutella       PCRVACQDAGVSHRFYLVHGQEGWFPLGTPCGGG...NASYCVSGKCLEFGPDNTPLSEM
Aedes          SCRIACQDNFIRHRFYLVNGEQGHFPFGARCNHDEP..NRYCVNGKCLRFGEDDTPLNES
Anopheles      SCRVACQDTFIRHRFYLVNGEQGHFPFGTRCNHNES..NRYCINGKCLHFGSDDIPVNES
Culex          SCRVACQDGFIRHRFYLVNGEQGHFPFGTRCNLDEP..NRFCVNGKCLRFGEDDTPVNEL
Leptinotarsa   PCRIACQDDFISHRFYLVNGEDGWFPFGTDCSRGTSGKKAYCVSGKCLEFGGDDTPLSES
Rhynchophorus  ............................................................
Periplaneta    PCRVACQDDEVNHRFYLVNGEDGWFPFGTDCSRGVPEKKSYCVSGKCLEFGSDDTPVHES
Blattella      .CKFLN......................................................
```

```
              1080                                1090      1100      1110
Drosophila     HISLGQLRTRR.......................KRSSISNDLFNITEIFIKPSHSDIS.
Ceratitis      HISLALFRSRRSAPSQDIVERRESAQTAVQSNRQRRSYLYFDPVNITETITQDFINNIVK
Spodoptera     VFTLPLLSRGGNASVSWRTSRS......VSSRHRRRRSLGAQRITVKATLDQTHLDDIIA
Plutella       VFTLPLLNRTTHSAVRRHSSR.............HRRSLQREKVVVTATLKQSHLDHIIK
Aedes          YNSLAHLRTKRT..................ARRIKRHFEFFSPINVTERISQEYIENLIA
Anopheles      YNALPNVRIKRS..................TNRTKRHFEYFAPINVTERLSQEFLENLIA
Culex          YNSLAHLRAKRS..................LSRVRRHFEFYSPINITERINQEYLDNLIA
Leptinotarsa   EFTLPLLSRS........................RRSLYNNGSIRIMTKLNQMELEDIIS
Rhynchophorus  ............................................................
Periplaneta    EYTLPLIGRL........................KRSLVYNSTTMVRTTLDQTTLDHIIS
Blattella      ............................................................
```

```
                                        1120      1130
Drosophila     ....................AENIEFTQPIHVSADELSSKSR..................
Ceratitis      SIMDFEQQGVED....ELLMRDHIEFNNPIHISTAEVTTTQPQPQQTTLKVA........
Spodoptera     RLNLTHNLNDHITYFDTIPEHIELDFSNPIHIAPEDTLLRKKPRPTWD............
Plutella       HLNFTENKNEPITYFEHIPEHIEVDFNNPIHIAPEESMFGKIPKPRWDRQVGTLSQYTHV
Aedes          N.MNFARNTDE.......IFEDNIDFRNPIYFSP..........................
Anopheles      N.IDFGRTTNEN....NIVLEEHIDLKNPIYVDP..........................
Culex          H.IDFEKTTGD.......VYEDSIDFRTPIYFSP..........................
Leptinotarsa   GLHRTLS......GNPSYPVKYDIDLNNPIHVNMDMEQQHHPQRYFRSEDYVNFA.....
Rhynchophorus  ............................................................
Periplaneta    KLNSTDNCNH...QYGSYGNQFSIDLNNPVHVEVKTGTGQFNNEARGYEDIWRSPG....
Blattella      ..............YISYPNKFM.....................................
```
