## Supplementary Alignment Files for "Assessing target genes for homing suppression gene drive": sxl.pdf

```
              1        10        20        30        40        50
Drosophila    MYGNNNPGSNNNNGGYPPYGYN.NKSSGGRGFGMSHSLPSGMDTEFS..FPSSSSRR..G
Ceratitis     MYG......NMNNGGHAPYGYNGYRPSGGRMWGMSHSLPSGMDTDFTSSYPGPSMNRRGG
Spodoptera    ..........MCERYPGYLFG..GTPVSSWDRSFNSLENQIQQQYR............
Plutella      .........MELNLEYP.DISS..LTSLGLLNPKDQNDFLQIQQQYL............
Aedes         MYS......KMN..GYPSSTFR...SSGGRLWGMSHSLPSGMDSDFS......YSSRPRM
Anopheles     MYS......KMN..GYPSSSFR...SSSGRLWGMSHSLPSGMDTDFS......SYSRPRL
Culex         MYS......KMN..GYPSSTFR...SSGGRLWGMSHSLPSGMDSDFS......YSPRTRM
Leptinotarsa  MNIFCLQVNGYYIWELDIKMFG...SQLGNFPFSVNPFLPED.TMNN............
Rhynchophorus ...................MFA...TQLGSFPFASNTQNTDIPHFNN............
Periplaneta   ............MDGYPRTLHY..VNQETSWEGEGPS.HSSARNSYQ............
Blattella     ............................................................
```

```
                 60        70                       80        90        100
Drosophila    YNDFPGCGGSGGNGGSANN.........LGGGNMCHLPPMASNNSLNNLCGLSLGSGGSD
Ceratitis     YNDFS..GGGGGGGGSA.............MGSMNNMVNAASTNSLN..CG...GGGGGD
Spodoptera    .......GRAGAGARCW.......................GSMSEGGG.....VSAAGGD
Plutella      .......GRH..GVRRW.......................GSMSEGG.......ACAG..
Aedes         F.DMPSSSTFGGNAPQT.....................SSSSTSSLS...GMNFGPGSNN
Anopheles     YGDMPSSSTFGGGNPSTPHHHHHHHNHLSQPASHHHTSSSSTSSLT...GMNFGPGSSG
Culex         Y.DLPSSNTFG.NVPNT.....................SSSSTSSLS...GMNFGPGGNN
Leptinotarsa  .......SSE..............................GTANTSN.......GAGDSN
Rhynchophorus .......NSA..............................NIMEQNN.......GAGDSC
Periplaneta   .......NHR...............................PMAEYQ......QKNEG..
Blattella     .........................................MAEYQ......VKNEG..
```

```
                110       120       130       140       150       160
Drosophila    DLMNDPRASNTNLIVNYLPQDMTDRELYALFRAIGPINTCRIMRDYKTGYSFGYAFVDFT
Ceratitis     ..GHGGGSNGTNLIVNYLPQDMTDRELYALFRTIGPINTCRIMRDYKTGYSFGYAFVDFA
Spodoptera    S.......CKTNLIVNYLPQTMTEKDLYAMFMTIGPIESCRVMKDFKTGYSYGFGFVNFT
Plutella      .........RTNLIVNYLPQSVAEKDLYAMFTSVGPVESCRVMKDFKTGYSYGFGFVNFA
Aedes         ........AGTNLIVNYLPQDLTERELYSMFSTMGPIETCRIMRDVKTGYSYGFGFVNFL
Anopheles     T.......AGTNLIVNYLPQDMTEREMYSMFSAMGPIESCRLMRDLKTGYSYGFGFVNYL
Culex         Y.......SGTNLIVNYIPQDMSERELYSMFSTMGPIETCRIMRDMKSGYSYGFGFINFL
Leptinotarsa  Q....YQFDKTKLIVNYIPQFATENDLAQIFGPLGQLENIKIMKDFKTGYSFGFGFVKYS
Rhynchophorus S....GGLDKTKLIVNYIPQFATEGELLQIFAPIGTIESLKIMKDYKTGYSFGFGFVKYF
Periplaneta   .........KTNLIINYLPQNMTEKELYSMFVTIGPVESCRVMKDFKTGYSYGFGFVNYS
Blattella     .........KTNLIINYLPQNMTEKELYSMFVTIGPVESCRVMKDFK.............
```

```
                170       180       190       200       210       220
Drosophila    SEMDSQRAIKVLNGITVRNKRLKVSYARPGGESIKDTNLYVTNLPRTITDDQLDTIFGKY
Ceratitis     AETDSQRAIKSLNGITVRNKRLKVSYARPGGESIKDTNLYVTNLPRTITDDQLDTIFGKY
Spodoptera    REEDAARAIDTFNGYQLRNKRLKVSYARPSGDDIKDTNLYVTNLPRAITEEQLETIFGKY
Plutella      REEDAARAIDTFNGYQLRNKRLKVSYARPSGDDIKDTNLYVTNLPRAITEEQLETIFGKY
Aedes         SEDAAQRAIRCLNGYSVRNKRLKVSYARPQSDDIKETNLYITNLPRTITDEQLDIIFGKY
Anopheles     NEEAAQRAIKCLNGYPLRNKRLKVSYARPQSDDIKETNLYITNLPRTITEEQLDIIFGKY
Culex         TEEAAQRAIRCLNGYTVRNKRLKVSYARPQSDDIKETNLYITNLPRTITDEQLDIIFGKY
Leptinotarsa  SPEEAAKAIAVLNGFNFRNKRLKVSYSRPPGTDMKDSNLYITNLPKDVTEQDVDILFGEY
Rhynchophorus KEEDAAKAIEALNGFNFRNKRLKVSYSRPPGTDMKDSNLYITNLPKDVTEQDVENLFKEY
Periplaneta   KAEDASRAIGTLNGLQVQNKRLKVSFARPSGDEIKETNLYVTNLPRNITEQYLDMLFGKF
Blattella     .AEDASRAIGTLNGLQVQNKRLKVSFARPSGDEIKETNLYVTNLPRNITEQYLDMLFGKF
```

```
                230       240       250       260       270       280
Drosophila    GSIVQKNILRDKLTGRPRGVAFVRYNKREEAQEAISALNNVIPEGGSQPLSVRLAEEHGK
Ceratitis     GMIVQKNILRDKLTGKPRGVAFVRFNKREEAQEAISALNNVIPEGASQPLTVRLAEEHGK
Spodoptera    GRIVQKHILRDKSNGTPRGVAFVRFDKREEAQEAIAALNNVIPEGGTEPLSVKVAEEHGK
Plutella      GRIVQKHILRDKNNGTPRGVAFVRYDKREEAQEAIAALNNVIPEGGTEPLSVKVAEEHGK
Aedes         GTIVQKNILRDKLTGFPRGVAFVRFNKREEAQEAISALNNVIPQGGTQPLIVRVAEDHGR
Anopheles     GTIVQKNILRDKLTGQPRGVAFVRFNKREEAQEAISALNNVIPQGGNQPLIVRVAEDHGR
Culex         GTIVQKNILRDKLTGHPRGVAFVRFNKREEAQEAISALNNVIPQGSNQPLNVRIADDHGR
Leptinotarsa  GEIVQRTVLKDKITGMPRGVAFVRYSKGEEAQAAIANLDGKLLENAMLPLSVRVAEDHGR
Rhynchophorus GEIIQRTVLKDKHTGLPRGVAFVRYARGEEAQAAIANLDGKMLENAMMPLSVRVAEDHGR
Periplaneta   GQIVQKNLLKDKLTGMPRGVAFVRYDRRDEAQQAINSLNGVIPENGTEPLVVKIAEEHGK
Blattella     GQIVQKNLLKDKLTGMPRGVAFVRYDRRDEAQEAINSLNGIIPENGTEPLVVKIAEEHGK
```

```
                290       300       310       320       330       340
Drosophila    AKAAHFMSQMGVVPANVPPPPPQPPAHMAAAFNMMHRDGAMEKLRSLFDAICDAIFGLDS
Ceratitis     AKAQHYMSQLGLIGGGGG....GGGGGGGGGGMGGPPPPPMNMG..YNNMVHRGF....
Spodoptera    QKAAYYAGWAAGFHHNRAS....VAPQRHHSLDQLDMLHSRQDYTTNYGMYGGLSNRYLR
Plutella      QKAAYYAGWAAGFHHNRG......................EPCNEDSGMHD.......
Aedes         AKAALYVPSYNSIVHNNR.......GRVR....MRNNPY....................
Anopheles     AKAALYVPSYNSIVHNNR.......ALLPRQNTMKYNDRSSGLLRSHHSLIFY.......
Culex         AKAALFVPSYNSIVSNNR.......GRVR....MRNNPY....................
Leptinotarsa  QKAQYLEAWNPMA..........YN..RDSNN...DLSTVGN...SNQYK........
Rhynchophorus QKAQYLDNWNPMGGGFGGR....GYPGLRTSIMPFRDLTVPTRPILPNRYTRFPTFTSPN
Periplaneta   QKAAYYAGWQAGYNQSRD..........................................
Blattella     QKAAYYAE....................................................
```

```
                350       360       370       380       390       400
                 .         .         .         .         .         .
Drosophila    ENFADLLDGLYRRK.YHYPYLXLTPQQQQQLLQHQQQALGFTSSSNNSIGNGNGNDNNML
Ceratitis     .NYSGLIDGLYRNKSYHYPYL.......................................
Spodoptera    RDYFNQPSYMSHHRPF............................................
Plutella      ATFMSQYDDEFH................................................
Aedes         ............................................................
Anopheles     ............................................................
Culex         ............................................................
Leptinotarsa  .........FIHG...............................................
Rhynchophorus IRNLGAGDTFFQSPFYY...........................................
Periplaneta   LSLMSSKLAVFK................................................
Blattella     ............................................................

                410       420       430       440       450       460
                 .         .         .         .         .         .
Drosophila    LYHQQYHQQQTQQQRLGNVAAHNISPNGSNNNINTSNTNNINFNTIRQNGVAALHYLQEQ
Ceratitis     ............................................................
Spodoptera    ............................................................
Plutella      ............................................................
Aedes         ............................................................
Anopheles     ............................................................
Culex         ............................................................
Leptinotarsa  ............................................................
Rhynchophorus ............................................................
Periplaneta   ............................................................
Blattella     ............................................................

                470       480       490       500       510       520
                 .         .         .         .         .         .
Drosophila    LQLQQPQDQQSQQQQPLTMPSSPPFQQQSRQSHHNGSSSTLGNQLLAISNNNSFNNNSNQ
Ceratitis     ............................................................
Spodoptera    ............................................................
Plutella      ............................................................
Aedes         ............................................................
Anopheles     ............................................................
Culex         ............................................................
Leptinotarsa  ............................................................
Rhynchophorus ............................................................
Periplaneta   ............................................................
Blattella     ............................................................

                530       540       550       560       570       580
                 .         .         .         .         .         .
Drosophila    SNSFTGNYSNGSAFTSNGAISGSNFPNNPTSSGNFTNNSTNSNPTNSGHFASNLAGSSNF
Ceratitis     ............................................................
Spodoptera    ............................................................
Plutella      ............................................................
Aedes         ............................................................
Anopheles     ............................................................
Culex         ............................................................
Leptinotarsa  ............................................................
Rhynchophorus ............................................................
Periplaneta   ............................................................
Blattella     ............................................................

                590       600       610       620       630       640
                 .         .         .         .         .         .
Drosophila    TNHLSGSNNYTNSNGNFTSNAASSSNFSNNAASSTNYSKNCSSGVVGNSDPDSCSRKHTT
Ceratitis     ............................................................
Spodoptera    ............................................................
Plutella      ............................................................
Aedes         ............................................................
Anopheles     ............................................................
Culex         ............................................................
Leptinotarsa  ............................................................
Rhynchophorus ............................................................
Periplaneta   ............................................................
Blattella     ............................................................

                650       660       670       680       690       700
                 .         .         .         .         .         .
Drosophila    NTNNHHTSPQHDFSFNMSTTEQELHQQTLKLQQLHLNNSFNNTTAAATTSAAASTSTSTT
Ceratitis     ............................................................
Spodoptera    ............................................................
Plutella      ............................................................
Aedes         ............................................................
Anopheles     ............................................................
Culex         ............................................................
Leptinotarsa  ............................................................
Rhynchophorus ............................................................
Periplaneta   ............................................................
Blattella     ............................................................
```

```
                  710       720
                    .         .
Drosophila     ATASSTNSANVGFLWRT
Ceratitis      .................
Spodoptera     .................
Plutella       .................
Aedes          .................
Anopheles      .................
Culex          .................
Leptinotarsa   .................
Rhynchophorus  .................
Periplaneta    .................
Blattella      .................
```
