## Supplementary Alignment Files for "Assessing target genes for homing suppression gene drive": tra.pdf

```
Drosophila   ............................................................
Ceratitis    MNMNITKASATTRKIRIEQNVPSGSVRKGPYAIERSVNPSEVVIKRRFGEGSKPLFQRDD
Periplaneta  ............................................................
Blattella    ............................................................

                                                                1
Drosophila   ........................................................MKMD
Ceratitis    IVVNPDNVVSNVGAHFETQPKDRSNNSKEEVENQWRKERHKSTDSSSPERFRKHHSSNKS
Periplaneta  ...............................................MTRNRIFYGEGLY
Blattella    ..........................................MRSRSPSRRSTPPPPRIS

                      10        20        30        40        50
Drosophila   ADSSGTQHRDSRGSRSRSRREREYHGRSSERDS.......RKKEHKIPYFADEVREQDRL
Ceratitis    EHSNSGNNITTRHTKTHHPSQENLNTASKRRDSSPPTNRRHRTPEKVPYFIDEIRERDRI
Periplaneta  G.DYEKIEDNWRHPTHSGIQPPGFPFPRRSRSP.......LQREPKARSFSPGKRKLSPK
Blattella    RRSPGAFRQRGKSPERRRRMVPPAGVPKRSRSP.......VQREPKGRSFSPSKRKLSPK

                                        60        70        80        90
Drosophila   R...................RLRQRAHQSTRRTRSRSRSQSSIRESRHRRHRQRSRSRN
Ceratitis    RRKYGKRSTKSPSPPVMSSKFRRRRSYSRSISRSRSHSPARSKNRTHVYGSLSRRSSSVD
Periplaneta  R...................ERE........RERDRSREQDRERSSRPIIPPRKRTRS
Blattella    R...................DRERIRQRDPERERMRTRDPDRERGRSRERPPMREREPS

                       100        110        120        130
Drosophila   RN....RSRS....SERKRRQHSRS.....RSSERRRRQRSPHR.YNPPP.........
Ceratitis    RYIGGGRKRRRENLRTERDRDGQYRHHGHRSEEQERSRRGRSPRARTRSRTRSRERSKHV
Periplaneta  KS...PETAVS....SSRE.GGHMGS....YAGNVVGPPEVSPGRSYQGPSCPYPQGG.S
Blattella    HMDLVPPVKRT....RSRSPDGPMGSRLSNYAGSVCGPSDLSPARSYQGPPGMYPQG..P

                    140       150          160                170
Drosophila   ....KIINYYVQVPPQDFYGMS...GMQQSFGYQRLPRPPP........FPPAPYRYRQR
Ceratitis    RARNDERNKNLHGNHDELTNAELNQRNLTQPQIITIPVPVPADFLN.YAYSTWPTQTQWS
Periplaneta  GPDGDHPNRPMVSLSERFHGSGGPGSYKKEPSDHPVFRGPEGTGFDVNELKKITVDIRRN
Blattella    RSDREETNRNVVSLSERFHSSG.PGSYKKE.SEHPVFRGLEGSGFDLTDLKKITMVIRRN

            180                190
Drosophila   PPFIGVPRFG.......YRNAG..RPPY................................
Ceratitis    HPMTPPPRYGAPAYHMPTILPATVMPPMRPALPPYGLPPQPMRYGGRGLRFPQQHGPRPW
Periplaneta  LPVDMVPIERNIINPEDVVLVR..RPATEDNAYHYKGERKLIEFSKDGRTMDEPDSKACL
Blattella    PPVEMVPIERNILNPEDVVLKR..RPGM................................

Drosophila   ..............
Ceratitis    RPNFRPKTHK....
Periplaneta  GLITAEWPNMAPTT
Blattella    ..............
```
