## Supplementary Alignment Files for "Assessing target genes for homing suppression gene drive": vir.pdf

```
                 1        10        20        30        40        50        60
Drosophila       MADVDDGSELLFFDTFSHEEVTDINLDLVQFPKPVFITQVRIIPLGARVQADFPGGVRLG
Ceratitis        MVETDDAAELLFFDTFSHEVDTDINLDLVQFPKPVFITQVRIIPLGARVQADFPGGVRLG
Spodoptera       .MSTNEQPDLLFFDTFSHDTSEELNLDLVQFPKSVYVREIRIIPLGARVEGDFPGGVRLG
Plutella         .MSSSDQPDLLFFDTFSHDTSEELNLDLVQFPKSVYVREIRIIPLGARVEGDFPGGVRLG
Aedes            ..MDEETPELLFFDTFSHDTYEKLNLDLVQFPKPVYITEVRIIPLGARVQADFPGGVRLG
Anopheles        ..MDSELPELLFFDTFSHDTYEQN.LDLVQFPKPVYITEVRIIPLGARVHADFPGGVRLG
Culex            ..MDEETPELLFFDTFSHDTYEKLNLDLVQFPKPVYITEVRIIPLGARVQADFPGGVRLG
Leptinotarsa     ..MSEQIIELLFFDTFAHDNTEEINLDLVQFPKPVYVTEVRIIPLGARVQADFPGGVRLG
Rhynchophorus    ..MTENIVELLFFDTFAHDNTEEINLDLVQFPKPVYVTEVRIIPLGARVQADFPGGVRLG
Periplaneta      ...MADSTELLFFDTFSHESSEELNLDLVQFPKPVFISEVRIIPLGARVQADFPGGVRLG
Blattella        ............................................................

                           70        80        90       100
Drosophila       ATNPSKFDLEFFVNDLGMPAASAFENLGLLRYNQNDCIHLDCSQ................
Ceratitis        ATNPSKFDIEFFVNDLGMPGASTFENLGQLKYNQNDCIHLECTQ................
Spodoptera       ATNPTKFHIDFFVNDLSKPGASTFEALGSLDYCQNGQIHMECGSGLDQ............
Plutella         ATNPTKFHIDFFVNDLSKPGASTFEALGSLDYCQNGQIHMECAAGPED............
Aedes            ATNPSQFKIALFVNDLGKPGAPTFECLGDFEYNQNNCIHLECGKPDDGS...........
Anopheles        ATNPSNFHIQLYVNDLGKPGAPIFESLGELEYNQNNCIHLQCGGGGSGAVGTGPGTEEFV
Culex            ATNPSLFKIELFVNDLGKPGAPTFECLGDFEYNQNNCIHLECGKPDDGA...........
Leptinotarsa     ATNPSQFDIEFFVNDLGKPGASSFETLGGFKYDQNGCINLECTPEETV............
Rhynchophorus    ATNPSQFQIEFFVNDLGKPGASTFETLGAFEYDQNGKINLECTPDDSV............
Periplaneta      ATNPSQFEIEFFVNDLSKPGASTFESLGGLEYKQNVHIQLECD.................
Blattella        ............................................................

                     110       120       130       140       150
Drosophila       EKIVTDGLVLRGWYSTITLAIYGIFTNSVTEPIASPTLP.CEPVGP.EIAN.........
Ceratitis        EKIPTDGLVLRGWYSTITLAVYGILTNSMSEPIASPPPPPCEPTGPEEICN.........
Spodoptera       PRIPTDGLVLRGWYTTITLAVYGNLTQVLPETP.VG...ANPPPVVQRP...........
Plutella         THTPTDGLVLRGLYTTITLAVYGNLTQVIPETPPIGT..SVPPPSVPKQ...........
Aedes            RRIPTDGLVLKGFYTTITLAVYGILTANIAEPIVSPRETTPLPPQEAPV...........
Anopheles        RRIPTDGLVLKGWYTTMTLAVYGILTTNIAEPIVSPRETTPQPAEQLLVGPDHHHHLLHH
Culex            RRIPTDGLVLKGFYTTITLAVYGILTANIAEPIVSPRETTPQP.EVAPV...........
Leptinotarsa     RKIPTDGLVLKGWYTTITLAVYGTLTNNITEPIVQP...VANPSLPTQP...........
Rhynchophorus    RKIPTDGLVLKGLYTTITLAVYGTLTNNIAEQLVQP...VVNPALPNPP...........
Periplaneta      RRIPTDGLVLRGWYTTITLAVYGVLTKSVQEAPPPPPPGLITHSVPPPG...........
Blattella        ..IPTDGLVLRGWYTTITLAVYGVLTKSVQEAPPPPPQVLIPHGVPPPA...........

                               160       170       180              190
Drosophila       .....LSG.....EVLLQEDVLKDEWQEPMQA..ELLTA.......HKGNVSDYDPEDME
Ceratitis        .....LSGSGETREPSIPEETNKDEWKEPLPS..EVPTA.......HKTNIGDFERDDME
Spodoptera       .....VVTREVPPAPAPTVPPNP.EWSQEPSVPIPAYTG.......NVASANPDVYG...
Plutella         .....ALPREVAAVPQTSIPPPPVEWTQEAPNPIPTYTG.......QVAAAMPEVYG...
Aedes            ...EIVVSDVMTGGIAEP.GGVGPEWPEEPPT..VTVVP.......AEVFVQKEYPEAD.
Anopheles        HHHHHHHHLHHGGGIAEPSCGPGGDWIQEPPLPGLVAAA.......ETVFVQKEYPEDD.
Culex            ...EVVP..AVDGPIAEP.GGVGQEWPEEPP...AVVVA.......TEVFVQKEYPEPAD
Leptinotarsa     .....NTIAD..AQQICVVPPVETEWQPEAIPTVPVE.......YSAQPAAPYQAYN...
Rhynchophorus    .....NTIAD..AQQIISGSSVDAEWSQDNISTQPLE.......YPNQAATNYQQYN...
Periplaneta      .....PCPADPRVTDNIVTNSATAEWVQQHAQVAELAPPVDRG.YMVEPALT.DPYNNNY
Blattella        .....PCPTDPRVTDSIVANSATAEWVQRHAQVAELPPPVDRNNYLVEPALAPDPYNNNY

                          200
Drosophila       YG.........MSRDHYHQH...................AE...................
Ceratitis        YVGGVGRSGAGVGGDHYQHS...................GD...................
Spodoptera       ...PNNYP.ENYDGQMYRGDYY............EAEQPKDPRTYQHMEEN.........
Plutella         ...PPNYPPDSYDNQMYRADYY............DAEAPKDPRTY.HMEGP.........
Aedes            .GEDIPKDPRPYARRVAAAVVVAS..............PAEPSSP...............
Anopheles        .SEDAPKDPRPYARRAAAAAAATSSSIA.....TAHPLPASAASPPEPIYE.........
Culex            GGEDIPKDPRPYARRALHATAVVC..............PAEPSSP...............
Leptinotarsa     .......PPE..QYVQDFEYYP..............DVPKDPRSYHHTPET.........
Rhynchophorus    .......QPESYQHQEFSEYYN..............DVPKDPRSYHHTPET.........
Periplaneta      TG.EYHPPPPEYPPHSYREEWPQQPRTVLHGPGESYETPKDPRSIPPYEEERLPWESGG.
Blattella        SGEPYHPPPPDYPPHPYREEWPQQPPTVLHGPGDTYETPKDPRPVPPYEEERLPWESGGA

                                 210       220       230       240
Drosophila       .................EQEQREMRRLRRSTHSTDHSPPPPRRSHTHSESNDREYIR...
Ceratitis        .................ERE....RGLRKTSHSSERS..LPTRNRTHSESNEREYVRGRN
Spodoptera       ...........AWEK.RDLSCEREPDRMRHSPRSIDHDRDRRDARSRRRSLDRGRSRESS
Plutella         ...........EWDKDRRLSAERDRQSPR.SEHDRDRDRDRRDGRSRRRSLDR.RSRESS
Aedes            ..............KMMECDVVDKRGK.RVSRSTERQMHYETGVRSRGRS..HEYS.RSP
Anopheles        .........PLEAGKILPGECEPPPLADKRTKGAGPPYAYERSPRARRHSSREEFSSRSP
Culex            ..............KHVVVECEGAGGKKRVSRSTER..HYEP.RGARGRS..HEYS.RSP
Leptinotarsa     ...........EWET...KPRPRVSDAERDRSRESTYQKPAAPERDHVRSEPARSSREKR
Rhynchophorus    ...........EWD....KGRNRDSDRERNRD...MYQ.....DRDHER.......RDRR
Periplaneta      .....GVEDKLPWERDRAKDRDRDRERDRDRERDRERER.RDRDRDRDRD..RDSYRDRS
Blattella        GGGGSGVEEKLPWERDRVKDRDRDRDRDRDRERERDRDR.RDRERERERDSYRDSDRDRS
```

```
                 250                                     260
Drosophila     .CSRDKGS.......................RDWSRSPEYSSHR................
Ceratitis      EREREKVS.......................RDWSRSPDYSRH.................
Spodoptera     RLRDRSREIDR..............MDRLRSRSRSRDRDYVVKGEYRP............
Plutella       RLRDRSRELDR..............HDRLHSRSRSRDRDYIVKGEYRP............
Aedes          SLCKEYGG.................VGVGPKREWSRSPDYGN..................
Anopheles      SLCKDYGPMPGSGSSGVGGGGGASAAASKQRDQWSRSPDYIEPPSQQQQQQVGPPAHPPL
Culex          SLCKDYQGG.............AAPPVGGQKREWSRSPHRGS..................
Leptinotarsa   YSRSESRDQ.................ERQQ.REYREADAKEEA.DWPD............
Rhynchophorus  YSRSESRDR.................DRPI.KDYHEVARERDG.YDRE............
Periplaneta    RDRDRSRDR...............DADRDRDRDWEDRSRREDRERERD............
Blattella      RDRDRSREREREFRDR..DRDSERDRDRDRERDWDERSRRDDRDRERD............

                         270                           280          290         300
Drosophila     ..............SRRKRS...ER.......SRS.VVDEHKWP.RTPPASIDSPTRPRS
Ceratitis      ..............SRRKRS...ER.......SRSDAEESHKWPPRTPPASIDSPTRPRS
Spodoptera     ............LSRSRSRSID.RLRGISTDRDRDRDWDRGAYKKDDYRRHREPSYDRSR
Plutella       ............RTRSRSRSIDGRLR......SSERDWDRSSYKKDDYRRHRDGSYERSR
Aedes          .............RSRRIRSSSYDR..VRRGSSYD..LEKDDR..RRPRSPIDSPRRPRS
Anopheles      PPPTPSSSSGGHRSSRRLRSGSYER..PRREPSFER.LEKEERR.PRPRSPIGSPRRPRS
Culex          .............SSRRIRSSSYDRGGVRRGSTDDRGTTRDDRSGRRPRSPLDSPRRPRS
Leptinotarsa   ............HDRQDYRSERN..DHDWSDRDRDRGHDRD...............RDRS
Rhynchophorus  ............HERTDYRTERDRIDHEWQESERDRDRTHD...............RERT
Periplaneta    ............RERDRERERSTSSAREYQDVSRDRDWDRGGSGGGGGGSGWGGGRRERS
Blattella      ............RDRDRDRERSTSSAREYQEVSRDRDWDRG..SGGGGGSGWGSSRRERS

                     310
Drosophila     PDTMDYEDEDSRS...............................................
Ceratitis      PD..NYSEDD..V...............................................
Spodoptera     GGSYDKDAGSYERRSPYDKRAPSYERKIPY..............................
Plutella       AGSYDKRSGSYERRSPYDKRAPSYER..................................
Aedes          PDPMDSSNDYEEE...............................................
Anopheles      PDRNESSLEEEDEPEELVDDVEDVEDDLDE..............................
Culex          PDPHD...DYDED...............................................
Leptinotarsa   ERCYK.......................................................
Rhynchophorus  .RDRR.......................................................
Periplaneta    SLSMERLPWADRGDHPQSK.GPPLLPIPPDHVVPPPHHHPHHHHEGPGPGNLHRLPSHMH
Blattella      SLSVERVSWGERGEHSQSMKGPPLLPLPPD.VGPPPHHHPHHHHEGPGPGNLHRLP.HLH

Drosophila     ............................................................
Ceratitis      ............................................................
Spodoptera     ...EKRLSPYEKRASSYERRAPSYEKQSSYDRKR...............HSPYSRIRGSS
Plutella       ....KRLSPYDKRGSSYERRAPSYEKQSTYDKKR...............HSPYNRIRGSS
Aedes          ............................................................
Anopheles      ......EMELDGGYHRSNGAKYAGSSRKRRSMLMMMEPAIESGRPQQQQQQPPLPPLPVH
Culex          ............................................................
Leptinotarsa   .........HDDNYRRSYNRGE........DREE..........................
Rhynchophorus  .........HDDGYRRSERYRD........DREE..........................
Periplaneta    PHPPPGPSHHHDMA.RPHSRGERKPRDGSWERSQGPPPGPQGSLLHQPPDQPWEIGPGRE
Blattella      PHVPPGPPHHHDIGSRPHSRGERKPRDGSWERSQGPPPGPPGPLLHQPPEQPWDIGPGRE

                                                         320
Drosophila     .....................................HYKMQSSH................
Ceratitis      .....................................HYKLKTRS................
Spodoptera     YSSRSPSRDDPRKRPRTPPGES.......SRRPLSPRDVDATS.................
Plutella       YSSRSPSREDPRKRPRTPPGE.........RRPLSPR.EGKVS.................
Aedes          ..........................DFRHGAPGGGHGAKYEKR................
Anopheles      PSASAKHHSSAASSRSSSHHHHQRDHHYQHGHSHAHHGGQHQQHG...............
Culex          ..........................DFRR....GGHGKYGDK.................
Leptinotarsa   ...........RKRPRTPP.........MQS.PKRPHTPQVVENV...............
Rhynchophorus  ...........RKRPRTPP.........MQMSPKKPHSPPHIEQS...............
Periplaneta    MSQERDEVLSIRKRPRSPPSQPSP....ALRSPKRPHTPPLQQLPGGPITPEDKLPQQQM
Blattella      ISQERDDMSQGRKRPRSPPSQPSPAPGIALRSPKRPHTPPPPQQP.....PEDKLPQQQM

Drosophila     ............................................................
Ceratitis      ............................................................
Spodoptera     ............................................................
Plutella       ............................................................
Aedes          ............................................................
Anopheles      ............................................................
Culex          ............................................................
Leptinotarsa   ............................................................
Rhynchophorus  ............................................................
Periplaneta    APRWPSSASPSRSEAGKVKK........................................
Blattella      APRWP...SPSRSEGGKVKKGTFFVVDLIVSVFDSDRGPMLCLRVVGRRLMSRWLHVPDD
```

```
Drosophila     ............................................................
Ceratitis      ............................................................
Spodoptera     ..............................PINSIRSEEGADYDRSGKQIPRVDFYHQS.
Plutella       ..............................PTTSIRSETG.EFDRSGKQIPRVEFYHQS.
Aedes          ............................................................
Anopheles      ................................SSHHHHHHHQHHQQQQQQQQQQHVPER.
Culex          ............................................................
Leptinotarsa   ...............................ETVEAVQSEVQEAARGE............
Rhynchophorus  ...............................KDILSPNGHTDINVREEEII.........
Periplaneta    ...........EVSSPTPSSGAQQKFESLSPGDVESIS.EGEIPETEADLEGDTLALAKT
Blattella      LRPKGYAGPIAEVSSAPQSSGPPQKFESLSPGDVESISSEGEIPEAEGEMDGETHSLTK.

                                                     330
Drosophila     ............................YRHSSESLHRGERD..................
Ceratitis      ............................YMRSEESLTNGAGDTQT...............
Spodoptera     ............................YRHK..TSIRSPSQEAENAAYAEQQHNSLVTV
Plutella       ............................YRQKAVASIRSPSQEPENPPFIEAQHSSLVTV
Aedes          .......................TVEGRIKAAAGGATGSPASASATPVAS..........
Anopheles      ......................VVSPPPLVCDAVSVSVTPVASPTTMLPAGAATVP....
Culex          .........................PPRIKTTSVSAATPPGVGGVE..............
Leptinotarsa   ...........................KVEECAKSPGREAKD..................
Rhynchophorus  ...........................KKDENVQEEIDNSKE..................
Periplaneta    AETSELMVAGDIAGAAE......DEPSLKLEPAEDEIPPARPGSADRKIQPSTPPSLSAS
Blattella      PESSELLVADKNAGEVAGISDEASESALKLEQAEDEIPSARSESVERKAPSTPPTLISNS

                                                                340      350
Drosophila     ..........................................RDDEDRSCTPQEQFEPIL
Ceratitis      ..........................................LDDDETPGTPGEQYEPIL
Spodoptera     QIVDN..TTAKPIESPNR.........................NIDEDRSMDAEQFEPIL
Plutella       QIVDTPPVVKLPIESPGR.........................NVDDEKSMDAEQFEPIL
Aedes          ........................................PALIPEEDALSATEQFEPIL
Anopheles      ........................................AATLPDEDAASVPDQFEPIL
Culex          .............................................EEGLAPPTELFEAIL
Leptinotarsa   ............AHTP...........................PAEEPAPMGVEEFEPIL
Rhynchophorus  ............VKSP...........................INEEITHMDVEEFEPIL
Periplaneta    IPATGSPAKASAVSSPSNIIHGPPSTPPPTTVSNVATPVGSPIPPDEPSMMDVEPFEPIL
Blattella      APATSSPAKVSVVLSPLNISHGPPSTPPLTAMSSVPTPVGSPIPPDEP.SMDLEPFEPIL

                    360           370                 380       390       400
Drosophila     SDDEIIGDD....EEDDAVDAAAIAEYER........ELEAAAAAAPPAIDAFEPWQKPL
Ceratitis      SDDEIIGDD....DSNTAIEPVMGDELET........ELAAAANAVGPAILDFDPFTMPI
Spodoptera     SDEDICDDMEGPSYMEVDYDVNDYCGVDDIIKYYNPFKCDWNKYEYVNKIHLLGPNK.EG
Plutella       SDEDICDDLEAPAYMDVDYDVNEYCGVDDIIKYYNPFKAEWVQYENMNKIHLVAPVKKEG
Aedes          SDEEITEDY....DFDMDYEP........................TDEYYKAFNPFVAVL
Anopheles      SDEEIPEDF....DYDLESATGGGGGAGAPAPTSGSSGPAAYENLDEPPAKQFDPFGATL
Culex          SDEEIVEDF....DYDVEYEMGGGGAA.................GGEDYGKGFNPFAAVL
Leptinotarsa   SDEDILDD..GDHYQDLEYDYSTYTNNDDIIKLFVPGTTPLSKYPKAKSFTIK......N
Rhynchophorus  SDEDIVDD..GEHYQDSDYDYSAYTNNDDIIRLFIPGVTELKKYSKSKKYVIR......D
Periplaneta    SDEDIVDET.DPQFQDMDYDFPEYCD.DQLSKTFNPFACELQPLICLPDPGLS......P
Blattella      SDEDIVEET.DPQY..IDYEFPDYCD.ELLSKTFNPFACELQPLNCLTDPSLS......P

                      410       420       430       440         450       460
Drosophila     LVFEGDMAAHFCKELETLKLLFKKLVLQTRCENVNAFS..EEHGASVDEREQFVYLGEQL
Ceratitis      SRYECEMQTLCRKELDTLAAIMQKFDIQTKCASLEAFN..DAS.TTIEERENFVYLCEQL
Spodoptera     AKDVLTATFDELCDASPELFKISELAKSNKKLDQKFFASTFLEIDNTTREEWVHQCEQLF
Plutella       VKDVMNATFEELCSANADLFKISELTKACK...GKFFASTFAEIDNASREDWVHQCEQLF
Aedes          QKYTP......DEGADAIPAVVKHLENLLKNTKYGAKATIDFGGESADFKEGFVDYTEQC
Anopheles      ERYVPAGADPVRLLGPGEQKLLRQLEQLLGQERYGRAATGAFEREPAEWKERFVDYAEQL
Culex          KKFELG.....VEKGGDETNAVRQLEIVLKNNRYNKKTTLEFGAESGDFKEGFVDYTEQI
Leptinotarsa   NSIVIDDCLKTAIGIADDYFKSSITKYDLGTFHKLNTEIKEEFIHLCEKVTPMIGSSAIF
Rhynchophorus  NSIELEEGLKTIIGIADDYLKSSITKYVLDNFNKHNTEIKEEFVHLCEKLICTIGEPPIF
Periplaneta    CEVEMKRLLR....LRTDPCLRPTVEMPTEASRLEDIILQQGKIISVQAENTCSGEPQVH
Blattella      CEVEMKRLLR....LQSEPCLRPTVEPTGEATKLEEIILQHEKMFNCNTEDSNSGEPQVH

                        470       480           490       500       510
Drosophila     NNQLGYLAQHYKRRNFVLQQFF....GNDELHLRQAANVLQIALSFQAACMQPQPAFKIR
Ceratitis      INQLSYITQNFKRRNYVLKQFF....NTNPNFLTKSYNILRIALEFEAACSQLQPAFKIR
Spodoptera     VSITTLFKNTDIVLRIFKNPNRNDSSDEFTEIYQLLISFCKIGLTFDLALAQQQPTYKIR
Plutella       LSLTSLCKTSDIILRIFKNLDKEEP.DEFNDIYNLLIIFTKIGLSFDSALAQQQPTYKVR
Aedes          VQLLNQIFAHDEWKDKFEKSVNGWIDGKSEGV.EILMNCIKLGLDWDRAMTQPQPASKLR
Anopheles      VHLLGQLLAKDDARDACQQQLDRWIQTGWPGVTDRLVNCICVGLDFERAMVQPQPAFKLR
Culex          VLLLNQIFSETPQRVKFERTLEDWIAGKSEGV.EIMLNCIKLGLDVDRAMTHPQPAYKLR
Leptinotarsa   CSIAKLYVDTKSMNPQSLSAQDKELTEQVLFIVETLVDWLKVALSYEMALIQDQPAYKTR
Rhynchophorus  VDIMKVHNVVRSLITPKMSVEDRELANQTNFIIETITDWLKISLNYDMANAQDQPAYKIR
Periplaneta    EDWVHLAEQVPPILIKALPYISNREG.....VLQTLFRWVEIGLDFNLALSQPQPGYKVR
Blattella      EDWVHLAEQVPPLLMKALPHISNRED.....ILQTLFKWVEMGLDFNLALGQPQPGYKVR
```

```
                  520       530       540       550       560       570
Drosophila        HIKLGARMAELLG...SSEELFQHLLKEH...KFDIFEAVFRLYHEPYMALSIKLQLLKA
Ceratitis         HLKIGARLTELLS...SAPQFLDHLLNVQ...NFDPFDVLFKLYQENYMAVSIKLMLLKA
Spodoptera        HMKCGIRLAEALMGHRHNGEVTKLLLSSG....VDMPMLLLDMYSREYMALSIRLMILKA
Plutella          HMKCGIRLAEGLLSHKNNGQVTKILLSIG....INIPMQLLEMYSKEYMALSIRLMILKS
Aedes             HMKAGIRLTEFLG...STEAFIQVILDQK...DFNIFDKLLGLYSQSYMALSIKLMIIRS
Anopheles         HMKSGIRLAEFLG...PHEPFIALLLRRARPDEFNLFDRLLGLYSQPHMALSIKLMIIRT
Culex             HMKAGIRLIELLG...ATDAFIKVVLEQE...DFNIFDKLLGLYSQSYMALSIKLMIIRS
Leptinotarsa      HIKCGVRLADWCC...NSIDFVRLLWQND....FNVHNALLNLYDQEFMALSIKLMILKA
Rhynchophorus     HIKCGVRLADWLC...HSEEYIQYIWNKN....YNLHRELLDLYRQEFMALSIKLMILKA
Periplaneta       HIKAGVRLVDALC...HCGETVTEQLLEA....QDIHSKLLDLYHRDYMALSIKLMILRA
Blattella         HIKAGVRLVDALC...HCGESVSERLLEA....QDIHSKLLDLYHKDYMALSIKLMILRA
```

```
                        580       590                             600
Drosophila        VYALLDTRMGIEHFMG.AKNN..........................GYQMIVEAIKTAK
Ceratitis         IYALLDTKEGIRYFLIGGKVN..........................GYQLLLNTLQQSH
Spodoptera        LNACLSSKESIDHFMKETVFP...KFDKSDDRK...........PRNGYQALITMMQTNP
Plutella          LNAALSSKEAVEHFMQKAVFP...RFGKNDDSESG.........PKNGYQALLAIMQTNP
Aedes             IYACLDTKRGLDYFLS.PAYN..........................GYQILLQLIQENP
Anopheles         IYACLDTRTAVHYFLS.PSYN..........................GYQILLDLLQENP
Culex             ICACLDTKRGLDYFLS.PEYN..........................GYQILLDLIQENP
Leptinotarsa      LDTYLQHKFAIEKFLLGGIDDSICQNGFNDSTPVP.........HTNGYKVLVGFMLQQP
Rhynchophorus     LDTYLLHKFAIEKFLISNSVSNSVQNGYTDTVLPN.........QSNGYKIIIEALDKNP
Periplaneta       LDSSLSCQQAVQHFLGHRTRGHKIAQNRNSFVKVKDNSVKEIEGSTNGYRTLVKMIQGNQ
Blattella         LDSSLSCQQAVQYFLGFRTR......DKNSLVKVK.....EIQGSTSGYRTLVKMIQGNQ
```

```
                     610       620       630       640
Drosophila        LTRTKYALQAIIKKLHLWEGLESVQIWCRRLFVDRIIIPGNRDQME..............
Ceratitis         LTRTKFALQAIVKKLHLNEALESVRETCNKLFVLTNLPTS....................
Spodoptera        LARLKFSISALIKKLNIYELLQKLRNLVIDFSKS........................GD
Plutella          LARIKFSVSALIKKLNMYELLEKLHELVLKMNKSTAEDKP..................DE
Aedes             STRISFALKALLKKINMYENLDLIRSTISDMYAIRSEPNFKP..................
Anopheles         STRISFALKSLLRKISLYENLELIHATVEELYVRAVEEGGEPQQQQG............G
Culex             STRISFALKSLLKKINLYENLDLVRSTVSDVYATRNDPAFKP..................
Leptinotarsa      LARVKFALTSILKKLNLYEILHKMHSILIKLRDVS.........................
Rhynchophorus     LVRIKFALSSILKKLNLYEVLFKIYNILIKLRNVS.........................
Periplaneta       LARVKFAISCLLRKLHAYEVLERLKESVGQLIDASPPPGAASPEDERRRSSDTSPEPMPT
Blattella         LARVKFAISCLLRKLHAYEVLERLKECVGQLIDASPPP...SSEDERRRSTEASPD..PI
```

```
                  650       660       670       680       690       700
Drosophila        ...DTVITCQQIEFAFEMLMDALFSSQLSYLQPRRFLPVSKKFEVVTDPTAQRSFGNALQ
Ceratitis         ...ALAEQYVAIEQAIEQIMDALVVNSLSYQQPKRFLPVAKKFEIANDPAAQRSFANALQ
Spodoptera        DSELSEADSHFIVNSFEEVLHMYRSQCFHLSQPKRFLPVSAQFEISKEC.....SNETLL
Plutella          EVELTEKETDFIVDSLGEILYMYRSQCFHISQPKRFLPVSAQFEINKEC.....SNEILL
Aedes             ....DPAMVSILENCFAELSKALTWDAIAYSQPKRFLPVSAKFEIHKDIKSALAGNRSFV
Anopheles         AKRTDQELMELLRSSLTELTSAISLHEVSLAQPKRFLPVSAKFEIHKDIASSLNVSRTVV
Culex             ....DPAQLSVLENCFAELTKAITWDAIAYSQPKRFLPVSSKFEIHKDIRSSVTANRSFV
Leptinotarsa      .HDISAEEINLITKSLNEVLNYCRAGPFVLSQPKRFLPVSSQFEIMRN.....YSYNVLV
Rhynchophorus     .HDISAEEINLITKSLNQILSYCNSGPFTMSQPKRFLPVSAQFEIMRA.....DSRNVLV
Periplaneta       NVDVSDQEVETIVSCLEEVLRVYREAPLAISQPKRFLPVCAQFEISPCSQ...DPYPALY
Blattella         TAEVSDQEVETIVSCLEEILRVYREAPLAISQPKRFLPVCAQFEISPCSQ...DPYPALY
```

```
                  710       720       730       740       750       760
Drosophila        SYLGQNSLAESLLVMLANCKELPATTYLS.MLDLMHTLLRSHVGIDYFVDDAFPVTQTIV
Ceratitis         SYFVEHALGESLLLMLVNAGELPSSLLLR.VLDLLQALLQTHVGVDYFVDDCFETTQLLI
Spodoptera        EFFNIHQILEVCLYLLTCPGTCNNLVVVSPIHDLIYELINSQNGLNFLYNN.IEMSGLLF
Plutella          EFFNIHRILEVCLYLLTAPTTCNNLVVTSPIHDFLYELIHSRNGLNFLYKN.MDLSELIF
Aedes             AYFNVHKLLETIILLLEMKLPFSS.LIGA.VYDLLSALLKNHQKLHYLADH.IDSVNVLL
Anopheles         QFFKIHRLVESLVLLLDLPGTPPA.LCEL.IYNLLACLLKSVPKLRYLVDS.LPYTNLLL
Culex             AFFNIHHLLETIIFLLEMKLPSSS.LIGA.VYDLLSALLKNHQKLDYLMEN.VDAVNVLL
Leptinotarsa      VYFKMFNLLQCFVLLLNYPSTMNLPLIKTPIFEIISILLDKSEGLEYLSEN.VETITVLL
Rhynchophorus     HYFGMFNLLQCFLLLLTYPSTLNLPPIKTPIFEIVSALLDNTEGLQYLSEN.SSTVNVLL
Periplaneta       AFFRCHRLLEVCLVLLSCPATAGYSAITVPVQEILAALLDSQGGMQFLASN.ADISNAIL
Blattella         AFFRSHRLLEVCLVLLSCPATSGYSAITVPVQEILAALLDSQGGMQFLASN.AHITNAIL
```

```
                    770       780       790       800         810
Drosophila        AILLGLDEVPRNPEEKEEKAEKSDAEDKAMEVENEAVEAG...GEKPTPPTADE......
Ceratitis         AVLLGIDEVPELVTTTEEETTESKVKEELLIPKTEAVSDDSAVKKETTASTAEQTDTLPK
Spodoptera        KTLTLPYS..QQNTE.............................................
Plutella          KILVQPYG..QQNAE.............................................
Aedes             KILFQHCPTPD...........ESHPEDLLH.............................
Anopheles         RWLFQRLPASAGGEQSPGGRSAPTVDEVQVS.............................
Culex             KILFQHCPSS.............SADEDFLH.............................
Leptinotarsa      KCLLRTDEEMQYSAA.............................................
Rhynchophorus     KCLLRTDEEMQYSLS.............................................
Periplaneta       RVLLQQAPPVSPGPP.............................................
Blattella         RVLLQQSPPVTPGPP.............................................
```

```
                      820       830       840       850       860
Drosophila     .......EGKPVAAPISVPAPAAAPQVRPRPILRPVLPRLARLGIEMSYKVQTRYHLDAI
Ceratitis      DNGTKESEKQSESAVPSKESTPASVEPNVQPLRRPVLHRLQQLAVELANKVQTRYHLDAI
Spodoptera     ..........................DFVYPHDLSNYSDLQILGLELAYRLKALYYLQSI
Plutella       ..........................DFVYLHDTSNYTELQIIGLELAYRLKSLYYLESI
Aedes          ...................................ELTRGQLLGLEMSYKLQTVFYLDAI
Anopheles      ...................................ELSRPQLLGVELVYKVQTIYHLDAI
Culex          ...................................DLTRGQLLGLEMAYKLQTLYYIDAI
Leptinotarsa   ..........................EFV........LKSHQLGLKIAYKLQALYHVEFL
Rhynchophorus  ..........................DCIE.......VKSHNLGLRIAYKLQCLYHIECL
Periplaneta    ..........................EDVLDEGGISAPPAQQLGLYMAYRLQALQYIDTL
Blattella      ..........................EDVLDEGAMSAPPAQQLGLYMAYRLQALQCIDAL

               870                 880       890       900       910       920
Drosophila     AYAAA........APEYDAVKLATHMHAIYSQTCDPAGRQHTVEVLGLNNNLKIFMDLIK
Ceratitis      MYLTR........ADNYDVIKLATHLHALYSQTCQSPGRQHTVDVISMNNNMQLFMDLIQ
Spodoptera     CDQQAGKA.........DENELIDRLQSLYCLSFGSIGRTAVPDVIVMGDNAECLMEVFE
Plutella       CDIQAGQP.........DENELIDRLQSLYCLSFGTVGKTAVPDVIAMGDNAECLMEVFE
Aedes          G............HTKDDLDKLVEHLQGLFYLANS.FGKRYVVEVLSMDGNLKLFLDLIE
Anopheles      ANASTGSVAPSNDQLEEQRDRLVDGLHALYELSCG.PGRRFVVEVLSMDDNLAVLLELLE
Culex          G............QTREDVDRQVESLQALFQLACS.FGKRYVVEVICMEQNMKLFLELID
Leptinotarsa   TANGKLHN......LDCDAPEIIDHLHGLLCLTFSHFGKVFCAETLGLEDNIRCLLQFLE
Rhynchophorus  LDSAKSNS......FDIDAFDIIDQLHGLFCLTFSPVGKISVSEMLGMNENARCLLQFFD
Periplaneta    LDLGSSDPS....YLDPDQTDVLDNLNGLYCLGFSPVGKLSLVHVLSRGYNIRVLLRFFT
Blattella      LYLGSSDPT....YLDPDQTEVLDNLNGLYCLGFSPVGKLSLVHVLSRGNNIRVLLRFFT

                       930       940       950       960       970
Drosophila     KEQRLQTQR.....QLSSPGTKYKSPVLSYAVDMVDACVRYCEQLDYLIEHGGVILELAK
Ceratitis      KEQRLQAQR.....QIASPGAKYKSPVLSYAVDMVDCCVRYCEQMDYLIEHGPMILELAK
Spodoptera     NDLKSKTKNE........SPLKQKSPAKGYAIELIVSAVRFAANVPFLKKYGQRLINVSR
Plutella       RDLKGKTKTD........SPPKQKSPAKGYAIELIVLAVRFSADVPFLKKFGPRILNVAR
Aedes          SEKRALMKD.......GSPGVKHKSPILSYSVDLVDCVVRHCPNVGYLVKHGETLLNLVK
Anopheles      REQRTVAKES..VGGTNAAATKHRSPIVGYCVDLVDCTVRHGEAVGYLAQHGPALLALAK
Culex          GEKRALAKE.......GSPGMKHKSPVLSYSVDLVDCAVRHCEHVGYLVQHGEKLLNLVK
Leptinotarsa   PVLAKEKTET......FVGRLKDSVGIPYIVDLVLLAVATVSNIPILEKHSKVLLHLVN
Rhynchophorus  NVP...KSES.......YLQKLKKSSGTGYVLDLLYYTIVVVPNIKFLEKNTKTINYILN
Periplaneta    DDNSHEQSQTQTLKYDLSDSKIKKSPGKSYSADLIAMTVKYSDYVPFLQKFGKEILEAAL
Blattella      GDSSHEQSQSQSLKYDINESKLKKSPGKNYAADLIVMTVKYSDYVPFLQRFGKEILNAAI

                 980       990       1000      1010      1020      1030
Drosophila     NHETFEPSVSAVLQEMYVYMKPLEAINVFVYDDIMPLVEVIGRSLDYLTTFPGDLIMAMR
Ceratitis      NHETFEPSVSAVLQEMFVYMKPLEAINIFSYDNITPLVEVISRSMEYITTFPGDLIMGLR
Spodoptera     EHDRFEPSVSSVLQEVIPYLKPVEKPTVLSGEDMTECVEILKASLEHAPDLPGELMTCLR
Plutella       EHDRFEPSVSSVLQEVVPYLTPIEKTNLLSGDDMVECVEIIKTSAEHAPDLPGELMTCLR
Aedes          QHEQFESSVSAMLQEMAVYLKPLEIHELFTNEDINPLVELMTRSLEFITTFPGDLIMALR
Anopheles      QHEQFEPGVSATLQELAVYLKPLELHDLFAYVDIGPLVELMRRSLEYVTAFPGDLITALR
Culex          THEQFESSVSAMLQEMAVYLKPLEIHELFTYEDINPLVELMKRSLEFITTFPGDLIMSLR
Leptinotarsa   QQDVFEDSVCLKLSELRLYLAPLENVTKFAYDDISLFVEAVDKSVENLPYTMSTAITALR
Rhynchophorus  ..LIYDDQYAFKINEIRLHLTPLEQAVSLNYDNISTFVEIMEKQMENVISHPGPLITALR
Periplaneta    K..DEQQSPSCRLFEVVPWLKPVENPSSFAYDDIGMLSEIIKRNVENATSLPGELVTAVR
Blattella      K..VEHQSPYCRLFEVVSWLKPVENPLSFAYDDIGMLSDIIKRNVDNATSLPGELITAVR

               1040      1050      1060      1070      1080      1090
Drosophila     ILRYLSISK......PLAGQKAPPVTEELKHRFVALQLYAADGVQLCIQIMERLCAYFEQ
Ceratitis      ILRHLAIGP......SNKAFRSS.STEELKHRFVTLQFYAADGVQTLIQILEKLCSYFEQ
Spodoptera     ILRHYCISDYENNVTIACES.ATEEYVELKYKYNVLQLFSLDGVAHLAGVLDRLATHFDQ
Plutella       ILRHFCISDYNANITVATES.STDEYVELKYKYSILQLFSLDGVAHLAGILDRIATHFDQ
Aedes          LLRYFGIPE......FDGQG..QPSHIELKYKYTILQFYSADGKTLLISILEKLNTYFDQ
Anopheles      LVRLLGIPP......YDDYE.QPGGHAEFKYKYTLLQFYSADGQALLLAILDKLTHHFDQ
Culex          LLRFLGIPE......HDGIVGANPSHVELKYKYTILQFYSADGKTLLISILEKLNTYFDQ
Leptinotarsa   IIQHLGISKHQPS...SSDN.PISNFVELKYKHVILQLFSLEGTTILSKVLQKICDHFEQ
Rhynchophorus  IVQYLGISNYQNKDIILLEN.PLTNYVELKYKHVILQLYSLDGTIVLVKILQKICNHYEQ
Periplaneta    ILKFLGIPLRDKDLAAVHETVDAEEYVELKYKYVILQLFSQEGVANLTAILQKLCEYYEQ
Blattella      ILKYLGIPLRDKDLNAMHETGDAEEYVELKHKYVILQFFSQEGVTNLTAILQKLCEFYEQ

                     1100      1110      1120      1130      1140      1150
Drosophila     PGAHAPALMTIQGVHCCQIMLPTLQILRELLSYAILCRDGTYKDLTAIDHLVKVYYLLYY
Ceratitis      PGLHTPALMTIQGLHCCQILLPALQVLRAMLSYAIQCRDTEFKDLTAIDHLMKAYFLLHY
Spodoptera     PSMHTSLFASVKGLQLAQILLPCLRLLDSMLARVVRCRGPRFRDLTAAPVLLKTYGIAKA
Plutella       PSMHTAFFASVKGLQLAQMLLPCLRLLDEMLARVVRCRGSRFRDLTAAPVLLKTYGIAKT
Aedes          PAIHASSLASMQGILLTQILTPLIQVLRKLMTYLIECRNTEFKDLTVIEPLLRTYALMQS
Anopheles      PAIHTSSLASQQGVLLTQLLCPLVYILRKLVTYLIECRNTEFRDLTLIEPLLRTYTLMQC
Culex          PALHASSLASMQGVLLTQILTPLVQVLRKIMTYLIECRNTEFKDLTVIEPLLRTYALMQS
Leptinotarsa   PSLHSSFFVSQQGWCLLQVIHPAVLLLKQMLAYAIQCQNTNFKDLTAISVFLQTHNLLSC
Rhynchophorus  PGLHSSILASNHGLLLVSVIYPCIVLLKQMLAYVVQCRNTNFRDLTTIPVLLQTYNFLNS
Periplaneta    PMLHSTSFVGYQGAMLVSFILPAIQLVRRMLTYVIHCRNTEFKDLTAVPVLLQTYSLMHA
Blattella      PMLHSTSFVGYQGAMLVSFILPAIQLVRRMLTYVIHCRNTEFKDLTAVPVLLQTYSLMHA
```

```
               1160      1170      1180      1190      1200
Drosophila     FPTRCQAGPEVEQCKMEVVQTLLAYTQPNEQ....DEESLHKSLWTLMIREVLKN.VDGP
Ceratitis      FPTHSQAANEIAQAKQEIVQLLLAYTQPNEQ....DEESLHKSLWTLMIREILKN.IDGP
Spodoptera     YPVGSVGYRTAAKAAEASVRALLAYAQPIADDAN.DGDSIRRGPWTSLCSEVISYITTAP
Plutella       FPVGSVGYATASKAAEAAVRALLAYSQPIADDAK.DGDSIRRGPWTSLCSEVISYTMTAP
Aedes          VPPTATAINDAREIQKDIIKILLAFTQPTPMDGL.DTNNIHKSLWTQMVGELIKFILFGP
Anopheles      VPSVAAAGPDARQVQRETVRILLAYTQPTPPEGL.DTNNVHRSLWTQMVGELVRFTLAGP
Culex          VPVNAAAINDAREVQKDIIKILLTFTQPTPTDGL.DTNNIHKSLWTQMVGELIKFILYGP
Leptinotarsa   FPRSCAGYGLAQEVSACIVEALLVYTQPVSEQIS.EKDSLTKALWTQMCGEVIKFVTFSP
Rhynchophorus  YPITAIGAELANKSQIAIIDTLLVYTQPLSDEIH.EKDSLNKTLWSQMCGEVIKFITTAP
Periplaneta    FPASALSYVDSQRVCREIIETLLAYTQPISSAPSSETEALNKSLWTMMMSEVVKYVTIGP
Blattella      FPASALSYIDSQRVCREIVETLLAYTQPISSAPSSETEALNKSLWTMMMSEVVKYVTTGP
```

```
               1210      1220      1230      1240      1250      1260
Drosophila     AHFIPGLKLLAELLPLPLPMPQPLCDQLQQQHKQRLITERKLWSAHLHPQSGQIAKLVEA
Ceratitis      ATFMPGLHVLAELLPLPLPIPLPVDAQLPETQSQRLITERKLWSAHLHPQSAQIAKLIET
Spodoptera     YTFVPGLLVFSELLPLPLPMQT..KTPPTDRELADASNERRLWSAHLHALSNDLTDMIQI
Plutella       YTFVPGLLVFSELLPLPLPMQT..KIPPTERELADAFNERRLWSAHLHALSNDLIDMIQI
Aedes          NYFIPGLLVFSELLPLPLPIQC..RQELTPVEINKLVNERQLWSAHLHPQSPSITEMIQT
Anopheles      NYFLPGLLVLSELLPLPLPIHC..RQPLAPPELSRLIQERQLWSAHLHPQSPSIGELIQT
Culex          NYFIPGLLVFSELLPLPLPIQC..RQELTPTEISKLINERQLWSAHLHPQSPAITEMIQT
Leptinotarsa   HTFLSGLLVFSELLPLPLPIQT..RDEHTAEEVSWVINLRKLWSAHLHPHSAIIQDMVNK
Rhynchophorus  YTFVSGLLILSELLPLPLPIQT..RDELTKSEITWAINLRKLWSAHLYPYNNAIQEMVNR
Periplaneta    HTFVPGLLVLSELLPLPLPVQT..RTPLPEDEILRAVNGRKLWSAHVHSLGSGLQELIAT
Blattella      HTFVPGLLVLSELLPLPLPVQT..RTPLPEDEVLRAVNGRKLWSAHLHSLGSGLQELIAT
```

```
               1270      1280      1290      1300      1310
Drosophila     LAPSSFPQLSELLQRVCMQLSDLAPNMTLLIAKTITELLCNEYQTSNC............
Ceratitis      IAPSTFPPLVDLMTKVCLQLADLAPNMTLLISKTLSDLVCAEWQSASA............
Spodoptera     ICMSTYRPVVHMLRRVCVQIADLAPNTAATVARAAVGAVMRELKP...............
Plutella       ICMSTYRPVVHMLRRVCVQIADLAPNTAATVARAAIGAVTKELKA...............
Aedes          ICPSSYTPLLVIFSRVCLQLSDLAPNMTLLISKAITDLIVQEPLLPGN............
Anopheles      LCTSSYTPMLAVLSRVCLQLSDLAPNMTLLVSKSLADLIVQEPLLPGG............
Culex          ICPSSYTPLLVIFSRVCLQLSDLAPNMTLLISKAITDLTIQEPLLPNTPPDD........
Leptinotarsa   LCISTQPQLLNLLRRVCVQISDLSANSALMIARGVLDNVYDALIP..............K
Rhynchophorus  ICISTQPQLLNLLRRICVQISDLAANSALIIARGMLDAVYKALLT..............K
Periplaneta    MSGSSFQPLLQLLRRVCVQLADLAAPTALIVSRGVLDTVLASLQHVPSSSPQ.SLPMNNS
Blattella      MSGSSFQPLLQLLRRVCVQLADLAAPTALIVSRGVLDTVLASLQHIPPNSPQPNNCLQTS
```

```
                          1320      1330      1340                1350
Drosophila     ......IPTTNLERLLRFSTRLCAFAPLKSSMLSILSGK.............FWELFQSL
Ceratitis      ......TPTPHLARLLYFFARLTSFAALKISSISILSGK.............LWELFQSV
Spodoptera     ....GEPATASVARVLGFLACLVSHAPVKCAVLHAMNSG.......GPRATDVQTALCSV
Plutella       ....GEPANGSAARVLSFLACLVAHAPVKCAVLHAINSG.......GPKAADVQTALCGV
Aedes          ......LASTSHARLFSFLGSLVCHASVKVSVLSILSGR.............IMELMTSI
Anopheles      ......LASTSHARLVSFLATLVCHASVKISVLSILPGR.............IMDMLMGV
Culex          QKPLKGLATTAHARLFSFLGSLVCHASVKISVLSILSGR.............IMDLMTNI
Leptinotarsa   EDVKMGPCNAHIARLLNFLACLVTHNTIKCAVLHLMHTNSTVTLKTEERYVSLIQAFSQV
Rhynchophorus  DD...GPCTSHVARLLNFLACLVTHNTVKCAVLHLMHTNSNVTGNQENKYSSLILAFSNI
Periplaneta    LSTPESPCSGHTARLLNFLACLVTHASVKSAVLHLLAKGGSVK..SDERYPALVTSLCAI
Blattella      NVTPQFPCSGHTARLLNFLACLVTHASVKSAVLHLLAKGGSMK..SDERYPGLVTSLCAI
```

```
                    1360      1370      1380      1390          1400
Drosophila     LALN.EFNDVVSNCQEAVHRILDSFLDSGISLISHKST.............ASPALNLAA
Ceratitis      LCYTGQASGVVQHCQLAIQQILKSFIDAGISFVSPKST.............ATPELNLAA
Spodoptera     LG.LANSSNEHAAAQEYAAHALAAFCDAEITLTPVGG...............TTDIILSN
Plutella       LG.LSNASHEHATAQEFAALALAGLCDAEVTLTPLSA...............PAEKIVAN
Aedes          LLNPGQTP.AHQQTQENIYMVFQNLLDSEFAMMFGN...............IAPMMQLAC
Anopheles      LLNPGQTP.AHQQTQQHIYLTFQSLFDAEIALMFGDT..............IDPALQLAC
Culex          LLNPGQTP.AHQQTQENIYIVFQNLLDSEFSMVFAN...............ISPMMQLAC
Leptinotarsa   LK.NESTSNSHIQSQECVLSIIQSFCDVEITLSQN.............TGITSEEYLAN
Rhynchophorus  LKRSDVTSTSHIQAQECILSILQSFCDVEITLMQNPI.........EGTLAINSETYLAN
Periplaneta    LR.SPSDSPSHIQAQECVVSVIQSLCDTEVTLLPPPGTTLVGGGAGDPGSMPTPDVYLAN
Blattella      LH.SPSESPSHIQN..............................................
```

```
                    1410      1420      1430      1440      1450      1460
Drosophila     ALPPKELIPRIIDAVFSNLTSVEVTHGISILAVRNLVILTEHDFTFYHLAQLLKQKITEF
Ceratitis      ALPPKELIPRIIEAILTNLFSSDPHQDVCEVALNNLNLLTELDISMHHLTQQLKQRRAEF
Spodoptera     SLPNKDALVAFLDSTADCLESTTKTCSVASSILRAYFVLTEHEYGFQQFRKFVSKRRESL
Plutella       SLPNRETLSAILDATADCLASTNRTCSVASSILRAYFVLTEHEYGFQQFRKFAAKKRDLL
Aedes          SLPPKEIIESCCATIVENISSEEVGAGVMTAALRTMLLLTEHDITFTTLIAAIEKRKDPI
Anopheles      SLPPKDIIEAFCGACATNVTGAEIGLGVLMAALRTMLLLTEHDTTFVTLMANLMRKRDGW
Culex          SLPPKEIIESCCGTIVENIASEEVGMGVLVAAVRTMLLLTEHDITLSILLTAVEKRKDAM
Leptinotarsa   ALPVKEHLLMFIHMIWEHLTT.DNSFMTCLPIFRTLLMLTEHDYGFYHLKEHLLKKNDAF
Rhynchophorus  AMPIKEHLLTFIDMILDHLTS.DNSYVTYLPIVRTLLLLTEHNYGFYHLRERLLKREKLF
Periplaneta    ALPPRDLLITMCTIMLEHVANIEHSFTTLLPTVRTFLMLTEHDYGFYHLKSCLEKKTDAL
Blattella      ..................................LWLSASKREIQEEKRSSCLEKKTDAL
```

```
                1470      1480      1490      1500      1510
Drosophila     QAWMERVILHN.ETVEYN.ANIESLILLLRSLTQIE.PPP.AMSAMPHRTLKLGATELAQ
Ceratitis      QQWLEKFLKYA.EAEHARLAPIELFIGFLHSLSHIA.EYLQQQQTIPIRSMKWSIGEVAH
Spodoptera     GKFFKWALEDQGEDKAECLSLYIDLIRILKMEE.........GEGSVGRKTMLTVSEMAD
Plutella       GSFFKWVLETAGEDKLECLSLYVDLIRILRLEE.........GEGAVGRRSTLTPHDVAD
Aedes          MAIVQKFADKARENPDTYRQIFILLGEFWRSLVMID.ETSLVGFNLPKRTVKLTPEQL.A
Anopheles      CAVMHKFWRKARDGPDAYRALFVLLSDFWRSLAAYE.TDGCV..NLPKRTVKLPLDQL.G
Culex          LAVVQKFADKAREAPDTYRQLFILLTEFWRSLITID.ETALVGFNLPKRTIRLTPEQLGA
Leptinotarsa   LVLLEKLANNFSKSIAECLSTLNILVEFLRVCVTTDVA..EESLLYTPRKITLSVLELKK
Rhynchophorus  LTVLNKLSQNFSKTSAECLSTLNTLVEFLRVCLITEEN..DATLLYSSRTIKLTVQEVKN
Periplaneta    LNLFVKFCSGFSKDSSDCLSTLSTSLELLRVCISLEEDDAESGLGLPARSLVMSVPELAV
Blattella      LNLFVKFCSGFSKDSSDCLSTLSTSLELLRVCISLEEDETESGLGLPPRSLVMIVPELAV

               1520                                1530      1540
Drosophila     LVEFQDIEL...........................AKPPVLSRILTVMEKHKAVANEAA
Ceratitis      LLGFQKN.............................ADPHLIQRLSAAYTN....VKNKA
Spodoptera     MVGYS.....PSDEE...............................HPVLKLDKIFRPRE
Plutella       MVGYT.....PGEES...............................HPALAMESILK...
Aedes          SLVFLPSRKEGQEHS...................VADCTAMLKELLVFFKNDIKPIEAPE
Anopheles      MLMPLCDSEEDALLA.....................QMLPFVDELLDLYGGG.EAKEQPM
Culex          MLVPKVIKKEGQELPPSE................PGDCGQLLKELLVFFKPEPKPDAEPE
Leptinotarsa   LIGWR...TDGKPEN...............................HPLLSLETILKAVV
Rhynchophorus  LMGWEDASVDGEGNN...............................HPFLQLEEVLKKET
Periplaneta    VLGWDRAQAETQEEEPTDGASEGDAVESAEVKDSIETNCATRNKKKHPIFCLEKLLQDCS
Blattella      VLGWNCSSVEGQEEE.TECAMETDINESADIKESIDTNGAIRNKKKHPLFCLEKLLQDCS

          1550      1560      1570      1580      1590
Drosophila     LSDLKQLILLQASKQEILAGTSTETPPEAEGEANPSASSCSASLTVEP............
Ceratitis      LTQIKEEDDSNSP.IKLLAEIATEFEALEGSASPKTDEDGAGAQTSEP............
Spodoptera     QHSNFNHRFQDRNSDEDALANAGFLASHLQKLKTLEEPGTPPEVQDMV............
Plutella       ..........ERNIDEEVISNCSLIPPLLHDLPPLS.PRAGADPPSPP............
Aedes          PEVVEDSSQSVPAVPIVAADTLEENILYLMDQLKDVAPGVMETQFDFG...........V
Anopheles      QLDSGAAAEPPKPG....TDSLKDNLSFLRTQIELAVAAAAATATTEAAKSAPIDWSAVT
Culex          .....EAAQLVPPV....VDNLEDNLLMLLDQLKEAKSTGGECLFDFG...........V
Leptinotarsa   E.........EDNTFENFLEDLTSLLKLLDGDSSDTNKEMKDTCIDTV............
Rhynchophorus  E.........ENTSFETILEGLSLFLKQLQTLDQESENEIP...QEVL............
Periplaneta    S.........EEEALESLHENVASLIRLLEVEGNKPVENRE..LLEPT............
Blattella      S.........EEEALESLHENVASLIRLLEVEATKPIENKE..LPEPI............

                 1600      1610      1620      1630      1640      1650
Drosophila     ..YLPQAEGIVTQYEARPIFTRFCATAENAQLTARYWLDPLPIELIEDMNEPIYERIACD
Ceratitis      ..TLPQSEGIVMQFAARPIFVEFDAVADEQLLLTRYWLQVS....CVDLDHSKYERVPCD
Spodoptera     ...LPAPETLVAQFQARTVYSLGEASDE..RLTSSYWLNVPAAGGE..DDVNDSELVPCD
Plutella       ...AAPADPLVAQFAGRAGHAGGAGAEE..RLTSSYWLDVPPRHHE..EEVNDGELVSCD
Aedes          IEPLPQAEGIVTQYSSRNVFIVSEEFED..QLKASYWLSFSP.....CEDDLEREQIPCD
Anopheles      VDPLPQAEGIVTQYGSRSVYLLSEDCDD..QLKATDWLGFTP.....CDDDLEREQIPCD
Culex          VEPLPQAEGIVTQYASRSVFIVSEEFED..QLKASYWLSFSP.....CDEDLEREQIPCD
Leptinotarsa   ...LPVPESLLIQFSRRTIFSSSDACDE..RLTANYWLAIPAVD.....GETDSENVQCD
Rhynchophorus  ...LPPPESLLTQYASRIICSSSDACDE..RLTDRYWLTIPADE.....GEGELETVPCD
Periplaneta    ...LPAPEPLLSQFAARPVFVVGEVDDE..RLSSTYWLAPPPVDE....ADQDMDQVGCD
Blattella      ...LPAPEPLLSQFAARPVFVVGEVS.................................CD

Drosophila     L
Ceratitis      L
Spodoptera     V
Plutella       I
Aedes          L
Anopheles      L
Culex          L
Leptinotarsa   I
Rhynchophorus  L
Periplaneta    L
Blattella      L
```
